# Immune characterization and profiles of SARS-CoV-2 infected patients reveals potential host therapeutic targets and SARS-CoV-2 oncogenesis mechanism

**DOI:** 10.1101/2021.02.17.431721

**Authors:** Martine Policard, Sidharth Jain, Samantha Rego, Sivanesan Dakshanamurthy

## Abstract

The spread of SARS-CoV-2 and the increasing mortality rates of COVID-19 create an urgent need for treatments, which are currently lacking. Although vaccines have been approved by the FDA for emergency use in the U.S., patients will continue to require pharmacologic intervention to reduce morbidity and mortality as vaccine availability remains limited. The rise of new variants makes the development of therapeutic strategies even more crucial to combat the current pandemic and future outbreaks. Evidence from several studies suggests the host immune response to SARS-CoV-2 infection plays a critical role in disease pathogenesis. Consequently, host immune factors are becoming more recognized as potential biomarkers and therapeutic targets for COVID-19. To develop therapeutic strategies to combat current and future coronavirus outbreaks, understanding how the coronavirus hijacks the host immune system during and after the infection is crucial. In this study, we investigated immunological patterns or characteristics of the host immune response to SARS-CoV-2 infection that may contribute to the disease severity of COVID-19 patients. We analyzed large bulk RNASeq and single cell RNAseq data from COVID-19 patient samples to immunoprofile differentially expressed gene sets and analyzed pathways to identify human host protein targets. We observed an immunological profile of severe COVID-19 patients characterized by upregulated cytokines, interferon-induced proteins, and pronounced T cell lymphopenia, supporting findings by previous studies. We identified a number of host immune targets including PERK, PKR, TNF, NF-kB, and other key genes that modulate the significant pathways and genes identified in COVID-19 patients. Finally, we identified genes modulated by COVID-19 infection that are implicated in oncogenesis, including E2F transcription factors and RB1, suggesting a mechanism by which SARS-CoV-2 infection may contribute to oncogenesis. Further clinical investigation of these targets may lead to bonafide therapeutic strategies to treat the current COVID-19 pandemic and protect against future outbreaks and viral escape variants.

## Introduction

In December 2019, a cluster of pneumonia cases resembling viral pneumonia was detected in Wuhan, China, and led to a rapid outbreak^1^. Severe acute respiratory syndrome coronavirus 2 (SARS-CoV-2) is a novel coronavirus that was determined to be the causative agent^1^. The global spread of infection led the World Health Organization to declare a pandemic on March 11, 2020^2^. SARS-CoV-2 and the resulting disease, COVID-19, has caused a worldwide health crisis. As of February 14, 2021, over 108 million cases and 2.3 million deaths have been reported worldwide^3^. COVID-19 is highly transmissible, and increasing mortality rates create an urgent need for vaccines and treatments. Currently, the only antiviral treatment approved by the U.S. Food and Drug Administration (FDA) is Veklury (remdesivir) for use hospitalized COVID-19 patients and combination with Barcitinib was authorized for emergency use, after being shown to reduce recovery time and serious adverse effects^4^. Remdesivir, hydroxychloroquine, lopinavir/ritonavir, and interferon regimens were promising treatment prospects until October 2020. WHO concluded these treatments had little or no effect on mortality from the interim results of a large international trial investigating the effectiveness of repurposed drugs for the treatment of COVID-19^5^. Thus, there is no effective treatment as of date. Although vaccines have been authorized for emergency use in the U.S and globally, complications with distribution, compliance, and efficacy against novel variants make developing effective treatments crucial to our ability to combat the current pandemic and prepare for future outbreaks. To develop safe and effective therapeutic strategies, understanding how the mechanisms of infection impact host immunity are paramount. Increasing evidence from several studies suggests the host immune response to SARS-CoV-2 infection may have a critical role in the severity of COVID-19^6^. Consequently, immune mechanisms are becoming more recognized as potential biomarkers and therapeutic targets for COVID-19^6^. However, the immunopathology of COVID-19 is not well understood and presents challenges to treatment development.

Coronaviruses (CoVs) belong to the family *Coronaviridae* and are named based on their surface’s crown-like appearance^7^. CoVs are enveloped, positive-sense, single-stranded RNA viruses with the largest known RNA genome of 30 to 32 kilobases^7^. The outer envelope, made of phospholipid bilayers, is covered by two different types of spike proteins: the spike glycoprotein trimmer (S) that can be found in all CoVs, and the hemagglutinin-esterase (HE) that exists in some CoVs^7^. Inside the outer protein coating, the virion has a nucleocapsid composed of genomic RNA and phosphorylated nucleocapsid (N) protein^7^. The family Coronaviridae is divided into four genera, the Alphacoronavirus(α), Betacoronavirus (β), Gammacoronavirus(γ), and Deltacoronavirus (δ)^8^. Phylogenetic analysis of the viral genes molecularly characterized SARS-CoV-2 as a new β-CoV^9^. It is now considered one of the seven CoV family members that infect humans^9^. Until the discovery of SARS-CoV-2, six CoVs were known to infect humans, including HCoV-229E, HCoV-NL63, HCoV-OC43, HCoV-HKU1, SARS-CoV-1, and MERS-CoV^9^. SARS-CoV-1 and MERS-CoV have resulted in significant disease outbreaks with high mortality in 2002 and 2012, respectively; however, the other human CoVs remain associated with mild upper-respiratory-tract illnesses^9^. Coronaviruses impose a continuous threat to global public health, and therapeutic strategies are needed to counteract current and future infections.

SARS-CoV-2 belongs to the same lineage that causes SARS-CoV-1, but is genetically distinct and more infectious^8^. SARS-CoV-2 has structural differences in its surface proteins that enable stronger binding to the ACE 2 receptor and greater efficiency at invading host cells^10^. Also, SARS-CoV-2 has a greater affinity for the upper respiratory tract, which permits the virus to infect the upper respiratory tract and airways more easily^10^. The first step in SARS-CoV-2 infection is receptor-binding to the host. The S1 subunit of the S protein contains the receptor-binding domain that binds to the peptidase domain of angiotensin-converting enzyme 2 (ACE 2)^10^. Therefore, S protein is the mediator of host cell binding and entry. SARS-CoV-2 binds to ACE 2 as the host target cell receptor in collaboration with the host’s transmembrane serine protease 2 (TMPRSS2), a cell surface protein primarily expressed in the airway epithelial cells and vascular endothelial cells^10^. Binding to the host receptor leads to membrane fusion and releases the viral genome into the host cytoplasm^10^. Afterward, viral replication occurs, leading to viral assembly, maturation, and virus release^10^. Host factors influencing coronavirus replication and protein expression are crucial for regulating coronavirus infection^11^. Previous studies identified important host factors associated with coronavirus cell entry and fusion, including CEACAM1, ACE2, APN, DPP4, TMPRSS2, cathepsins, and furin^11^. Host antiviral, pro-inflammatory, translation, and unfolded protein response factors influenced by SARS-CoV-2 infection include eIF4F, GCN2, PERK, PKR, RIG-I, MDA-5, TLR3, IRF3, IRF7, type I interferon pro-inflammatory cytokines, and chemokines ^11^. These results and others underline the diversity of molecular pathways involved in SARS-CoV-2 infection.

The host immune response against SARS-CoV-2 is critical to control and eliminate the infection. Several studies have shown that SARS-CoV-2 dysregulates normal immune responses, resulting in uncontrolled inflammatory responses in severe and critical patients with COVID-19^6^. Increasing evidence demonstrates that immune patterns are closely associated with disease severity^12,6^. SARS-CoV-2 infection and replication trigger cytokine storm and a hyperinflammatory response, involving increased secretion of the pro-inflammatory cytokines and chemokines IL-6, IFNγ, MCP1, and IP-10 into the blood of afflicted patients^13^. Studies show multiple viral structural and non-structural proteins antagonize interferon responses contributing to inflammation^13^. SARS-CoV-2 is able to inhibit type I IFN responses in infected cells, leading to delayed or suppressed type I IFN responses and allowing the virus to replicate unchecked and induce tissue damage^14^. Lymphocytes are seemingly active but dysfunctional T cells in severe COVID-19 seem to be more activated and may exhibit a trend toward exhaustion based on the continuous expression of inhibitory markers^15^. A robust B cell response is observed in COVID-19 patients; however, high titers of antibodies are associated with severe clinical cases. Study data suggests crosstalk with monocytes might impair NK cell recognition and killing of SARS-CoV-2-infected cells^15^.

In summary, severe COVID-19 patients exhibit lymphopenia, lymphocyte activation and dysfunction, granulocyte and monocyte abnormalities, high cytokine levels, and an increase in immunoglobulin G (IgG) and total antibodies^6^. Thus, evidence from several studies suggests that the host immune response to SARS-CoV-2 infection plays a critical role in disease pathogenesis, thereby making efforts to characterize the host immune response critical to understanding COVID-19 and identifying potential therapeutic targets. In this study, we investigated the immune characteristics of the host immune response to SARS-CoV-2 infection that may contribute to the disease severity of COVID-19 patients. We curated four publicly available datasets from the NCBI GEO database and performed differential gene analysis^16^. We then applied pathway analyses of differentially expressed genes to identify significant pathways and upstream regulators to identify potential immune-related therapeutic targets for further studies.

## Materials and Methods

### Acquisition and Processing of Data

To understand the host immune response to SARS-CoV-2 infection, four publicly available gene expression datasets were selected from the Gene Expression Omnibus (GEO) repository. Relevant datasets were identified from the queries “single cell COVID19” and “COVID19 SARS Cov 2”. Selection was primarily focused on sample sources upper respiratory system, lungs, and blood. Two RNA-seq and two single cell RNA-seq datasets were acquired for data analysis (**Table 1**). For each dataset, a count matrix table containing the raw gene counts was imported into *Partek® Flow®* software (v10.0). The standard workflow within the software was used to process and quantify the gene counts **(Figure 1**).

**Table 1.** Publicly available datasets from the NCBI Gene Expression Omnibus (GEO) database included in analysis. Datasets were identified from the queries “single cell COVID19” and “COVID19 SARS CoV 2” in GEO DataSets.

| GEO Accession | Platform ID | Study Design | Sample Source | Sample Size | Infected | Controls | Disease State |
| --- | --- | --- | --- | --- | --- | --- | --- |
| GSE145926 | GPL23227 | Characterized the lung immune microenvironment in the bronchoalveolar lavage fluid (BALF) using 10x genomics to measure single-cell RNA sequence (scRNA-seq) | BALF | 12 | 9 | 3 | (3) Healthy<br>(3) Moderate<br>(6) Severe |
| GSE154567 | GPL24676 | Examined immune response using 10x genomics to measure single-cell RNA sequence (scRNA-seq) of blood buffy coat from COVID-19 patients | Blood buffy coat | 17 | 12 | 6 | (6) Convalescent<br>(5) Moderate<br>(6) Severe |
| GSE152418 | GPL24676 | Systems biology approach to analyze immune responses of COVID-19 patients using RNA seq-analysis | PBMC | 34 | 17 | 17 | (17) Healthy<br>(1) Convalescent<br>(4) Moderate<br>(8) Severe<br>(4) ICU |
| GSE15207<br>5 | GPL1857<br>3 | Examined host gene expression across infection status, viral load, age, and sex among RNA-sequencing profiles of nasopharyngeal swabs | Naso-pharyngeal swab from upper respiratory tract | 484 | 430 | 54 | N/A<br>(Infected vs Control) |
| **Totals** |  |  |  | **547** | **468** | **80** |  |

**Figure 1.**
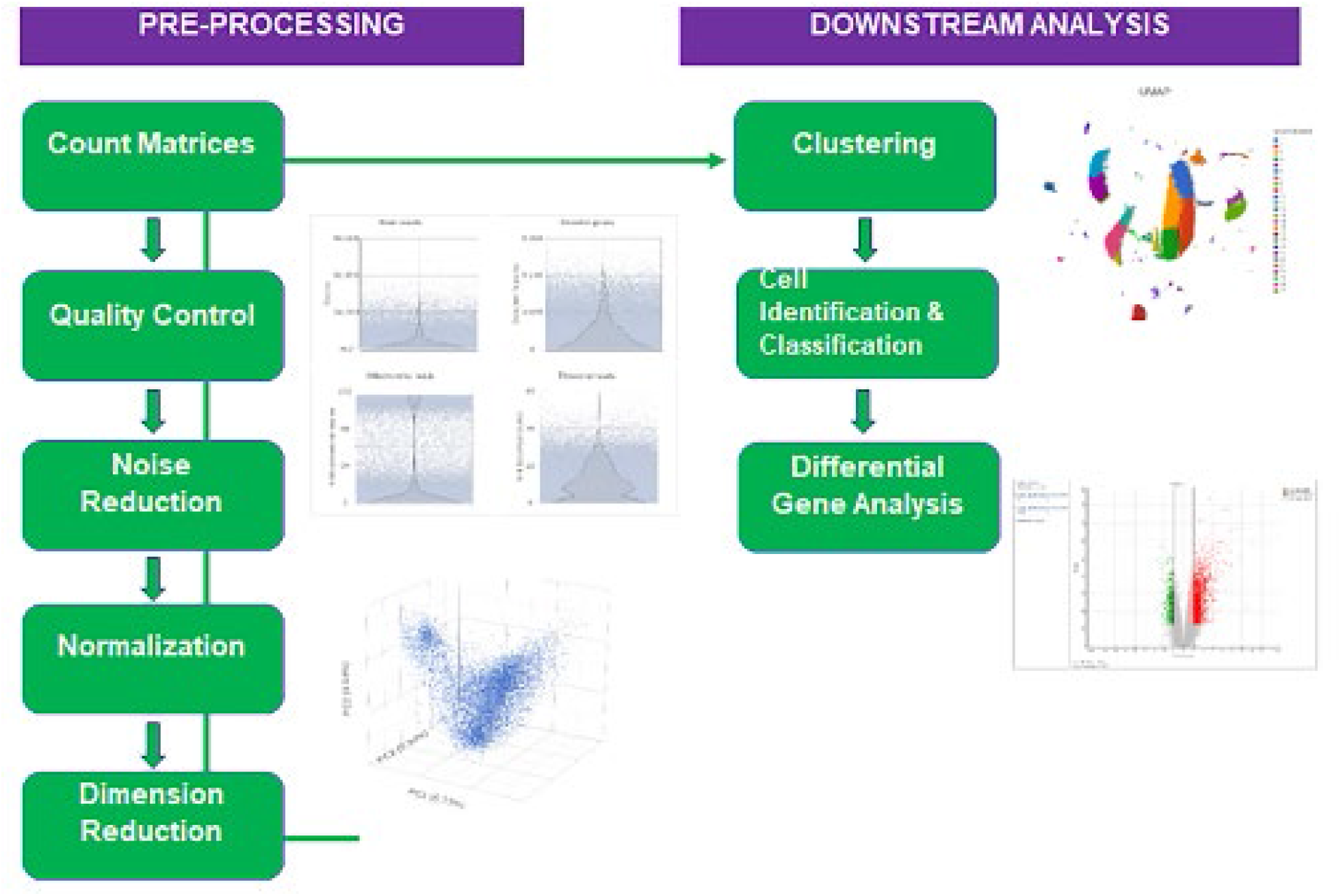
Schematic of standard single-cell RNA-seq pipeline in *Partek® Flow®*. Single cell count matrices were imported from GEO and processed for downstream analysis.

### Differential Gene Expression Analysis in RNA-seq datasets

GSE152075 and GSE 152418 count matrix tables were pre-processed by the authors. Therefore, no additional processing was necessary before performing differential gene analysis. Gene-specific analysis (GSA) was applied to the quantified gene counts in *Partek® Flow®* software (v.10.0) to identify differentially expressed genes in each dataset. Samples were labeled by the presence or absence of SARS-CoV-2 infection. For GSE152418, multiple comparisons of COVID-19 patients were analyzed by performing GSA twice. Similarly, to GSE152075, the first analysis was based on the presence or absence of detected SARS-CoV-2. The comparison included the following: SARS-CoV-2 infected patients (described by 4 moderate, 8 severe, and 4 ICU cases of COVID-19) vs healthy patients and one convalescent COVID-19 patient vs SARS-CoV-2 infected patients. A second GSA was performed to examine severity and the following comparisons were analyzed: Moderate vs Severe, Moderate vs ICU, and Severe vs ICU. Differentially expressed genes from each comparison were considered significant if the adjusted p-value (FDR) < 0.05 and log2 fold change ≥ ∓ 2. Genes are indicated by GeneCard codes, but more information about each gene is listed in **Supplementary Table 1**.

### Pre-processing scRNA-seq data

The recommended scRNA-seq pipeline in *Partek® Flow®* consists of several tasks to process single cell counts for further data analysis (**Figure 1**). First, low quality cells were filtered out using the single cell QA/QC task. Next, a noise reduction filter was applied to exclude genes considered as background noise. Genes that were not expressed by any cell in the dataset but were included in the matrix file were filtered out. Finally, data was normalized using counts per million (CPM), the recommended method in *Partek® Flow®*.

### Clustering Analysis and Cell Identification

The dimensionality of the data was reduced by performing principal components analysis (PCA) using 50 principal components. To identify similar groups in patient samples graph-based clustering analysis was performed and results were visualized in a 2D scatter plot using Uniform Manifold Approximation and Projection (UMAP), a visual dimensional reduction technique. Cell identification was accomplished by using UMAP to classify immune cell subsets by their phenotypic markers. NK cells were classified by NKGT (Natural Killer Cell Granule Protein 7) and NCAM1 also known as CD56. T cells were classified by TRAT1 (T Cell Receptor Associated Transmembrane Adaptor 1) which stabilizes the T-cell antigen receptor/CD3 complex at the surface of T-cells. Macrophages were characterized by CD68 and B cells were characterized by CD19. The classification result included a list of top featured biomarkers for each immune cell subset using an ANOVA test. Significance was determined by p-value < 0.05 and fold change ∓ 2. The immune cell classifications were applied to normalized gene counts and a differential gene analysis was performed to compare immune cell subsets in mild and severe COVID-19 patients.

### Differential Gene Analysis of scRNA-seq data

After immune cells were classified for each sample, the one-way analysis of variance (ANOVA) was performed to determine differentially expressed genes. For GSE145926, SARS-CoV-2 infected patients, including 3 mild cases and 6 severe cases, were compared to healthy patients. Additionally, severe versus mild COVID-19 patients were compared. For GSE154567, the following comparisons were analyzed for differential genes: severe versus mild COVID-19 patients, and convalescent patients versus severe COVID-19 patients. Furthermore, for each dataset, immune cell subsets were compared in mild versus severe COVID-19 patients. To immunoprofile genes, the 10xGenomics Human Immunology Panel was applied to the DEG output to filter and sort genes directly related to immunity. The panel contains 1,056 genes to profile innate and adaptive immunity, inflammation, and immuno-oncology from curated content including recent publications spanning key biomarkers, pathways, and lineage and tissue markers. Differentially expressed genes from each comparison were considered significant if the adjusted p-value (FDR) < 0.05 and log2 fold change ≥ ∓ 2.

### Pathway Analysis and Interaction Mapping

Differentially expressed genes identified by GSA or ANOVA in *Partek® Flow®* imported into Qiagen Inc. Ingenuity Pathway Analysis (IPA) software application. To identify significant pathways associated with the patterns of differential gene expression, core analyses were run in IPA separately for upregulated and downregulated genes in each dataset. Associated networks determined by the analyses were used to identify host-viral protein interactions and upstream or downstream regulators. Networks maps included an overlay of either the Coronavirus Pathogenesis Pathway or Coronavirus Replication Pathway. The comparison analysis function in IPA was used to compare the upregulated pathways and downregulated pathways across datasets.

## Results and Discussion

### RNA-seq profile of nasopharyngeal swabs of COVID-19 patients show upregulation of IFN-stimulated genes

The RNA-seq profile of nasopharyngeal swabs collected from the upper respiratory tract of 430 SARS-CoV-2 infected patients and 54 uninfected patients were retrieved from GSE152075. To identify differentially expressed genes (DEGs), gene-specific analysis (GSA) was performed in *Partek® Flow®* software. Genes were defined as significantly downregulated or upregulated if the threshold FDR value < 0.05, and fold change was ∓ 2. The volcano plot showed a greater number of genes were downregulated in SARS-CoV-2 patients, suggesting the infection has an inhibitory effect in the host’s upper respiratory tract **(Figure 2A)**. The topmost significantly downregulated genes (FDR <0.01) were DPM3, ROMO1, MT3, MRPL53, SCGB3A1. These are protein-coding genes that do not directly relate to the immune response, suggesting that other processes affected by SARS-CoV-2 infection may overlap with the immune system (**Supplementary Table 1**). In contrast, the topmost significantly upregulated genes (FDR <0.05) were found directly related to immunity or immune response in SARS-CoV-2 infected patients. These genes, including IFI44L, IFIT1, OAS3, and RSAD2, are primarily related to interferon-induced proteins that play important roles in the antiviral response (**Supplementary Table 1**). The significant upregulation of interferon stimulated genes suggests an activation of the interferon response in COVID-19 patients. The actions of interferon induced proteins may be a major component in antiviral host defense to SARS-CoV-2.

**Figure 2.**
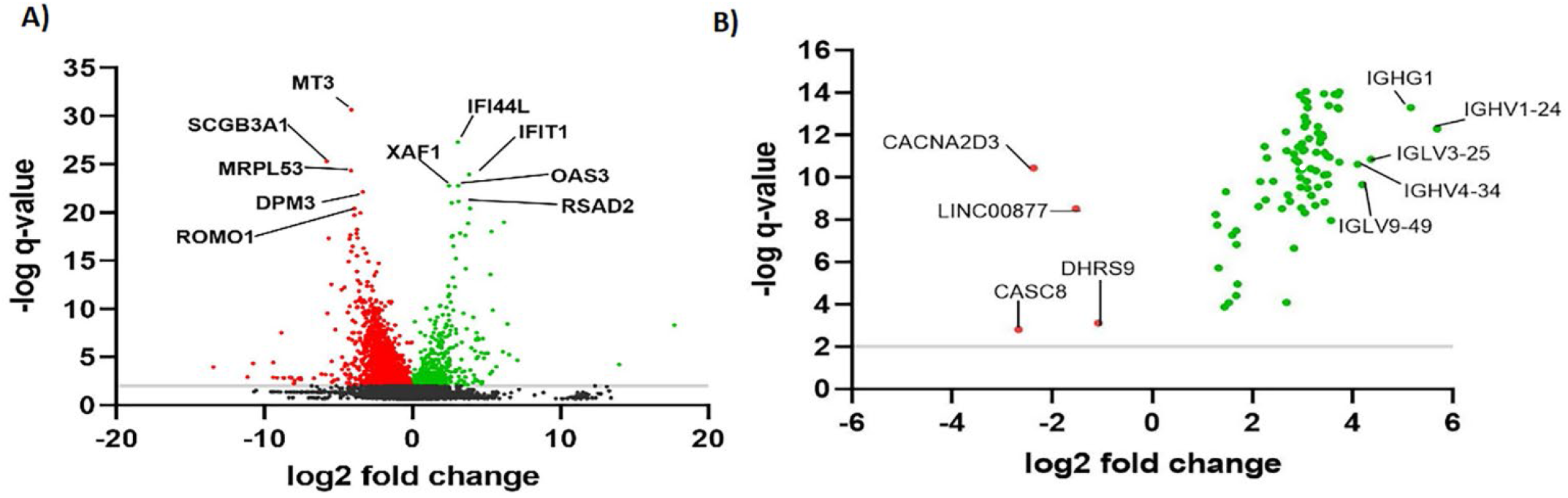
RNA-seq profile of nasopharyngeal swabs and PBMCs from COVID-19 patients. (**A**) Volcano plot of differentially expressed genes in SARS-CoV-2 infected patients versus uninfected patients from GSE152075 analysis. **(B**) Volcano plot of differentially expressed genes in SARS-CoV-2 infected patients versus uninfected patients from GSE152418 analysis. An adjusted p-value (q-value < 0.05) and fold change (log2 fold change ≥ ±2) were used to determine significantly downregulated or upregulated genes. The log2 fold change of the five topmost significantly upregulated and downregulated genes (FDR <0.01) are highlighted.

### Pathway analysis of significantly downregulated genes from nasopharyngeal swabs reveals indirect interactions that may overlap with the host immune response

Next, we applied IPA core analyses on DEGs to identify significant pathways and associated networks. The lists of significantly upregulated and downregulated were uploaded to IPA separately. In IPA, the core analysis for downregulated genes ranked “eIF2 signaling” as the most significantly affected pathway **(Figure 3A)**. Viruses hijack host machinery to complete viral replication and protein synthesis. In response, host cells shut down these systems, which is regarded as an integrated stress response^17^. The stress response induces translational shutdown via the phosphorylation eIF2α^17^. Therefore, sustained phosphorylation of eIF2α inhibits host or viral protein synthesis. Our IPA core analysis shows eIF2α downregulated in the eIF2 signaling pathway, as well as other mediators of the eIF2 complex, including eIF2γ. It remains unclear from the analysis which upstream regulator of the integrated stress response (PERK, HRI, PKR) plays a major role. It was previously reported that PERK is an important host factor that influences viral protein expression^11^. SARS-coronavirus infection activates PERK and PKR by phosphorylation, leading to sustained phosphorylation of eIF2α in 293T/ACE2 cells that inhibits protein synthesis^17^. SARS-CoV has also been shown to block the action of PKR to evade detection and suppress IFN production^18^. Thus, PERK and PKR may be potential targets for therapeutic strategies that prevent SARS-CoV-2 from hijacking the host’s machinery for protein synthesis, while targeting PKR may also help modulate the interferon response. IPA analysis indicated “Mitochondrial Dysfunction” as the second most significantly downregulated pathway, and “Oxidative Phosphorylation” as the third, based on p-value. Two of the most significantly downregulated genes from the differential gene analysis, MRPL53 and ROMO1, are potentially related. MRPL53 is a protein-coding gene for a mammalian mitochondrial ribosomal protein involved in protein synthesis within the mitochondrion^19^. Interestingly, mitochondria can play a regulatory role in the activation, differentiation, and survival of immune cells^20^. Mitochondrial dysfunction may be an indirect cause of the lymphopenia that is often observed in severe COVID-19 patients. While the analysis of downregulated genes in this RNA-profile does not directly relate to the immune response, there is a relationship between these genes and the host-viral interaction, which requires further investigation.

**Figure 3.**
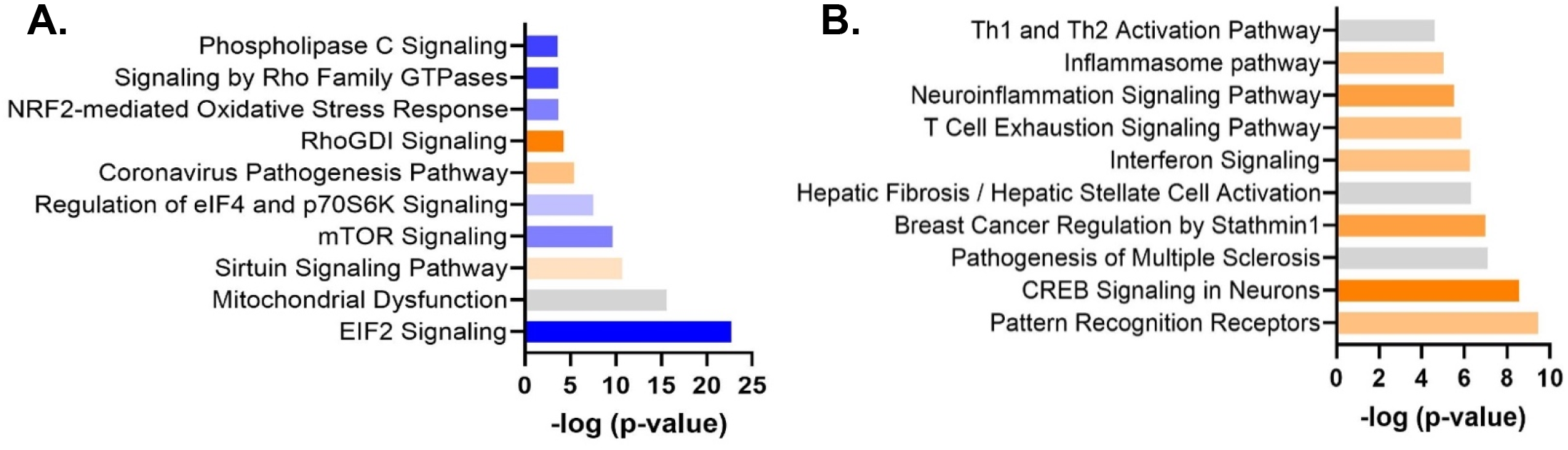
Pathway analysis of DEGs in nasopharyngeal swabs from COVID-19 patients. **(A)** Most significantly downregulated canonical pathways in IPA, determined by –log(p-value). The orange and blue-colored bars in the bar chart indicate predicted pathway activation or predicted inhibition, respectively, based on z-score. Gray bars indicate pathways for which no prediction could be made by IPA. **(B**) Most significantly upregulated canonical pathways in IPA.

### Pathway analysis of significantly upregulated genes from nasopharyngeal swabs reveals interferon-alpha as an upstream regulator

IPA core analysis of upregulated genes ranked “Role of Pattern Recognition Receptors in Recognition of Bacteria and Viruses” as the most significantly activated pathway **(Figure 3B)**. This pathway comprises numerous genes related to the innate immune response, such as toll-like receptors, TNF, complement proteins, and RNA-activated protein kinases. One of the activated networks observed from the IPA analysis was a “cell-mediated immune response” network highlighting interferon-alpha. **(Supplementary Figure 1)**. Interferon-alpha was identified as an upstream regulator in the pathway analysis. Several genes involved in the innate immune response appear related, such as ISG15, IFITM3, IFITM2, which are interferon-induced proteins. mTORC1, a regulator of cell metabolism in immunity, is also upregulated. These results show evidence of an active innate immune response. The reasons why this innate immune response is inadequate to combat viral infection and disease progression in COVID-19 patients remains to be determined. Several studies suggest an association of defective interferon activity with more severe disease in COVID-19 patients^21^. Thus, a dysregulated interferon response may cause a weakened innate response. It is unclear from the IPA analysis whether interferon signaling contributed to mild or severe disease in this RNA-seq profile because disease severity was not an available attribute in the original study imported from GEO. To characterize the immune characteristics that result in COVID-19, disease severity is an important attribute to study and analyze.

### Differential gene expression in the RNA-seq profiles of PBMCs collected from COVID-19 patients suggests host-virus interactions via EIF4E

GSE152418 consisted of the RNA-seq profiles of the peripheral mononuclear blood cells (PBMCs) collected from 17 SARS-CoV-2 infected patients and 17 uninfected patients. SARS-CoV-2 infected patients were compared based on disease severity for one convalescent, 4 mild, eight severe, and 4 ICU COVID-19 cases. Differentially expressed genes (DEGs) were identified by comparing RNAseq profiles of SARS-CoV-2 infected patients with uninfected patients. Genes were defined as significantly downregulated or upregulated if the threshold FDR value < 0.05, and fold change was ∓ 2. The volcano plot showed a more significant number of genes were upregulated in SARS-CoV-2 patients, suggesting the infection leads to activation in PBMCs. (**Figure 2B**). The most significantly upregulated genes (FDR <0.01) were not directly related to genes relevant to the immune response. However, the genes with the greatest differential expression (fold change ≥ 5) were constant or variable regions of immunoglobulin heavy and light chains (**Figure 2B**). Among the most significantly upregulated were IGHG1, IGHV4-59, IGHV1-24, IGHV1-46, IGKC. This result reflects the immunopathology of COVID-19 described in previous studies in which high titers of antibodies are correlated with severe disease^6^. This suggests that successful antibody response is not enough to counteract severe disease. The only genes significantly downregulated in SARS-CoV-2 infected patients in comparison to uninfected patients were CACNA2D3, LINC00877, DHRS9, and CASC8. The most significantly downregulated gene (FDR <0.01) out of the four was CACNA2D3, which encodes a protein in the voltage-dependent calcium channel complex. LINC00877 and CASC8 are non-coding RNA genes. In a study investigating the interaction between the cardiovascular system and the host antiviral defenses to SARS-CoV-2, non-coding RNAs reported to be dysregulated by transcriptomic profiling in cardiovascular diseases were also involved in viral innate immune responses^22^. DHRS9 is a dehydrogenase that has been shown as a stable marker for human regulatory macrophages^23^. Thus, although these genes are not directly related to the immune system, DHRS9 and non-coding RNAs have the potential to be indirectly related to the host immune response. Additionally, DEGs were identified by comparing the single convalescent patient with SARS-CoV-2 infected patients. Genes were defined significantly downregulated or upregulated if the threshold FDR value < 0.05 and fold change ≥∓ 2. All four of the significantly downregulated genes in SARS-CoV-2 infected patients were significantly upregulated in the convalescent patient. Similarly, all the significantly upregulated genes in SARS-CoV-2 infected patients were significantly downregulated in the convalescent patient. When other comparisons of disease severity were analyzed, no statistically significant genes were generated. Therefore, disease severity was not investigated in this dataset. Instead, the focus for the analysis of this dataset was the differential gene expression based on the presence or absence of SARS-CoV-2 infection, which excluded the one convalescent patient.

### Pathway analysis of differentially expressed genes in RNA-seq profiles of PBMCs reveals relationship between Coronavirus Pathogenesis Pathway and CSF2

Next, we applied IPA core analyses on DEGs to identify significant pathways and associated networks for GSE152418. Because there were few downregulated genes, the upload was not separated by upregulation or downregulation. All significant genes were imported into IPA. The analysis ranked “Kinetochore Metaphase Signaling Pathway” first and “Mitotic Roles of Polo-Like Kinase” second as the most significantly upregulated pathways. Interestingly, multiple pathways related to cell cycle regulation were among the most significantly affected, with “Cell Cycle: G2/M DNA Damage Checkpoint Regulation” indicated as the single most significantly inhibited pathway (**Figure 4**). While the genes annotated within these pathways do not appear relevant for host immune response to viral infection, the pathways indicated suggest dysregulation of the cell cycle, a possible connection to oncogenesis.

**Figure 4.**
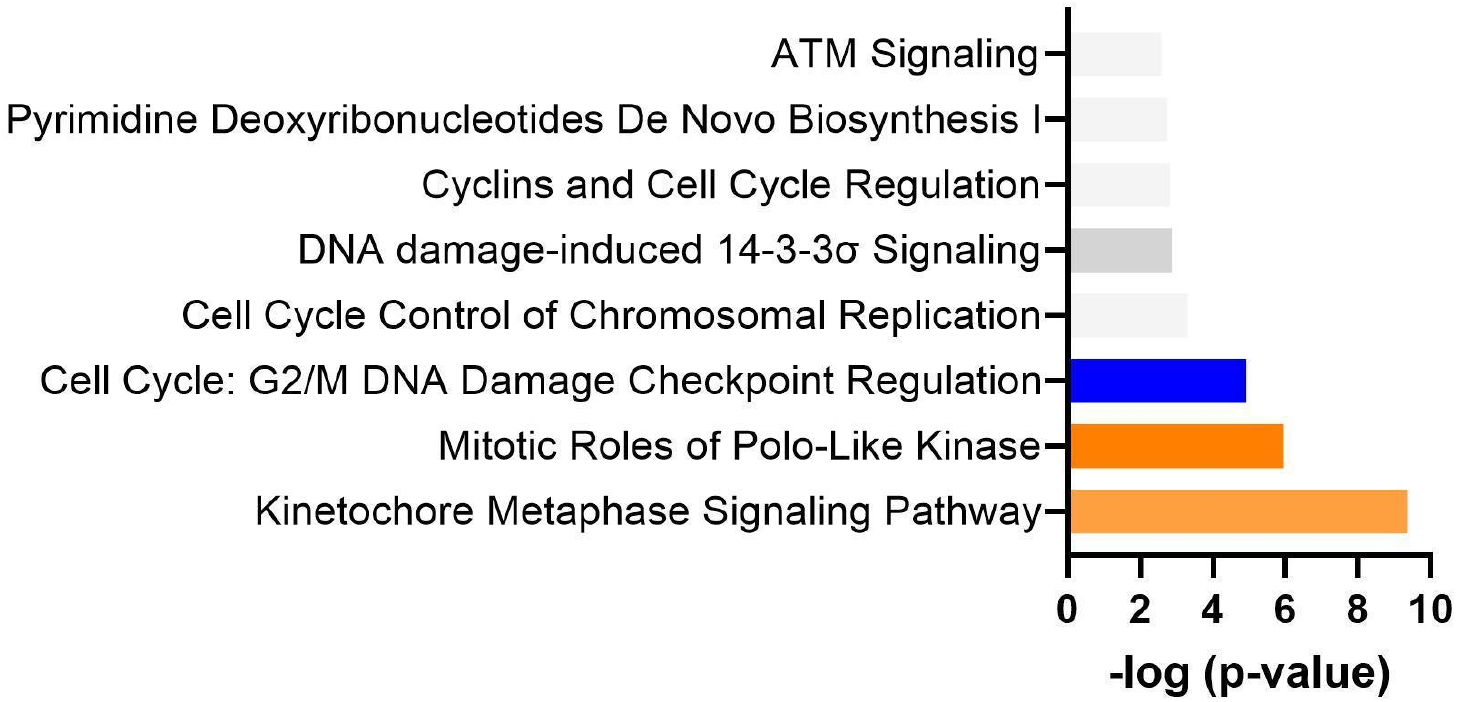
Pathway analysis of DEGs in PMBCs from COVID-19 patients. Kinetochore Metaphase Signaling Pathway was ranked first in the significant IPA canonical pathways for GSE152418 based on –log(p-value). The orange and blue-colored bars in the bar chart indicate predicted pathway activation or predicted inhibition, respectively. Gray bars indicate pathways for which no prediction can be made for the data provided.

Deeper analysis of the significant networks and pathways from this dataset indicated connections between SARS-CoV-2 infection and potential mechanisms of oncogenesis. One of the networks associated with this dataset revealed relationships between the Coronavirus Pathogenesis Pathway and E2F7 and RB1 (**Supplementary Figure 2**). E2F transcription factors control the transition from G1 to S phase in the cell cycle. Rb is a tumor suppressor gene that regulates the cell cycle by modulating the activity of E2F. Rb is often targeted by viral oncoproteins to inactivate its function and dysregulate the cell cycle, promoting entry into S phase and cell proliferation^24^. RB1 was significantly downregulated in COVID-19 patients while multiple E2F transcription factors were significantly upregulated, suggesting that SARS-CoV-2 may inactivate Rb tumor suppressor genes to promote the activity of E2F transcription factors, similar to other tumor viruses (**Figure 5**). This would inhibit regulation of the cell cycle and promote cell proliferation, suggesting a mechanism by which SARS-CoV-2 infection could contribute to oncogenesis. While these results do not directly implicate the immune response, they reveal more potential damage SARS-CoV-2 infection could cause, such as cancer.

**Figure 5.**
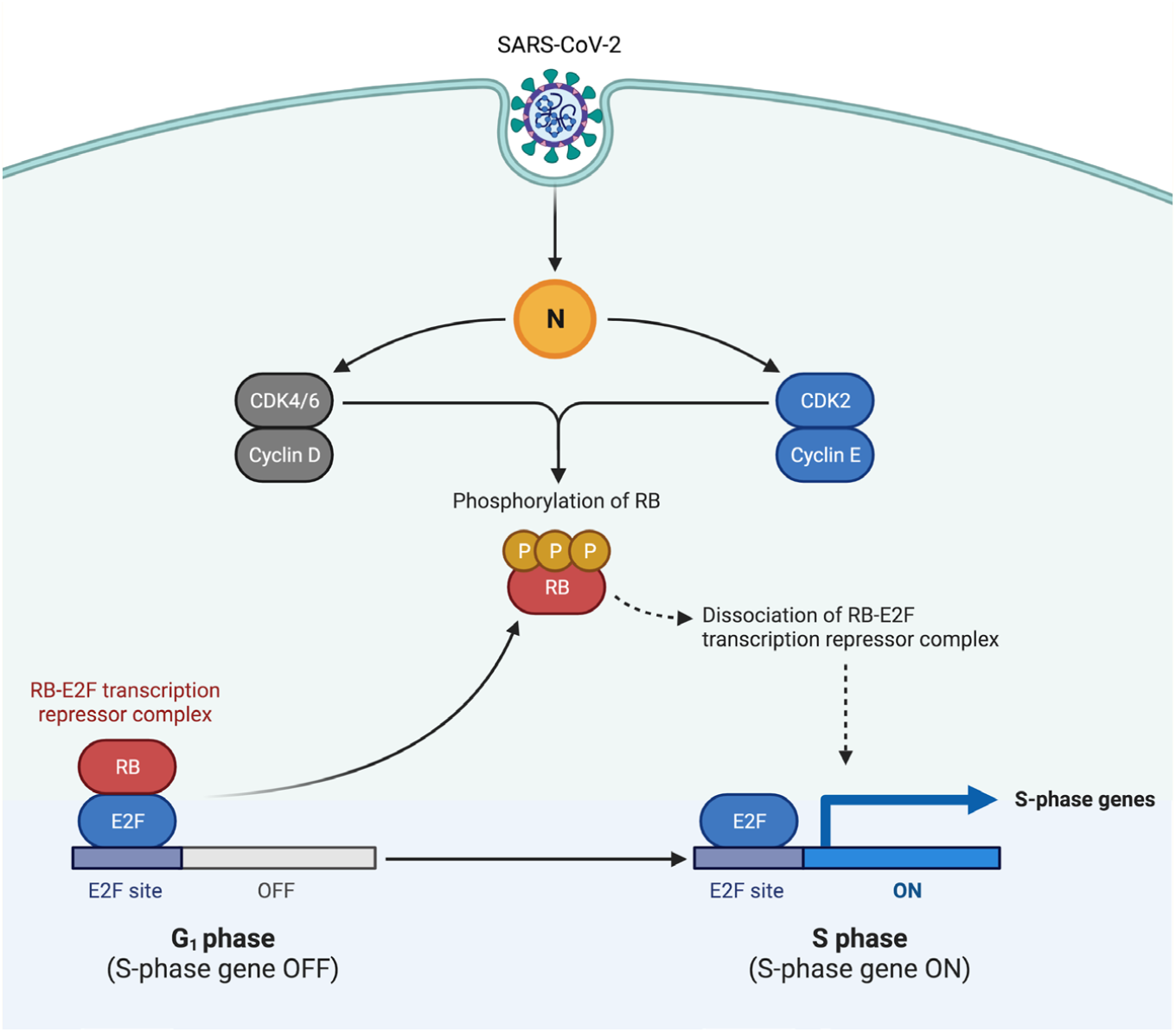
Potential oncogenesis mechanism of SARS-CoV-2 through interaction of the N protein with the Rb-E2F complex. This figure was created in BioRender based on the Coronavirus Pathogenesis Pathway from IPA and the G1/S Checkpoint in BioRender, modified to highlight the key genes most significant in the dataset GSE152418. Genes significantly downregulated are indicated in red and genes significantly upregulated are indicated in blue. Genes not significant in this dataset are indicated in gray and viral proteins are indicated in yellow. Adapted from “G1/S Checkpoint”, by BioRender.com (2020). Retrieved from https://app.biorender.com/biorender-templates.

### Classification and cell identification of scRNA-seq profile of BALF suggest T cell lymphopenia in severe COVID-19

GSE145926 consists of scRNA-seq data for the bronchoalveolar lavage fluid (BALF) of 9 SARS-CoV-2 infected patients and three uninfected patients. SARS-CoV-2 infected patients were compared based on disease severity, including three mild and six severe COVID-19 cases. We performed a graph-based cluster analysis to collate similar groups and classify four immune cell subsets (**Figure 6C**). Top biomarkers featured in this classification task were identified using ANOVA test with a threshold p<0.005. The output was used to verify markers for cell identification. (**Figure 6D**). We compared COVID-19 patients by disease state (**Figure 6A**) and the classified immune cell groups, including NK cells, T cells, B cells, and Macrophages (**Figure 6B**). A more significant number of T-cells were observed in the cluster of mild COVID-19 patients than severe. Conversely, a larger population of macrophages in the cluster of severe COVID-19 patients compared to mild. Considering the number of severe patients in this analysis is more significant than mild patients, this result suggests T cell lymphopenia in the severe COVID-19. Furthermore, when the classification counts were examined, there were 951 T cells in severe COVID-19 patients and 872 T cells in mild COVID-19 patients (**Supplementary Table 1**). Since a more significant portion of macrophages was observed in the T cell population, perhaps the macrophages can reduce T-cell activation through a dysregulated macrophage response. Numerous studies have reported lymphopenia as a common characteristic of immunopathology in severe COVID-19 patients, but the underlying mechanisms are not well understood^25^. Careful immune monitoring during clinical studies should be considered in future studies to help reveal these mechanisms.

**Figure 6.**
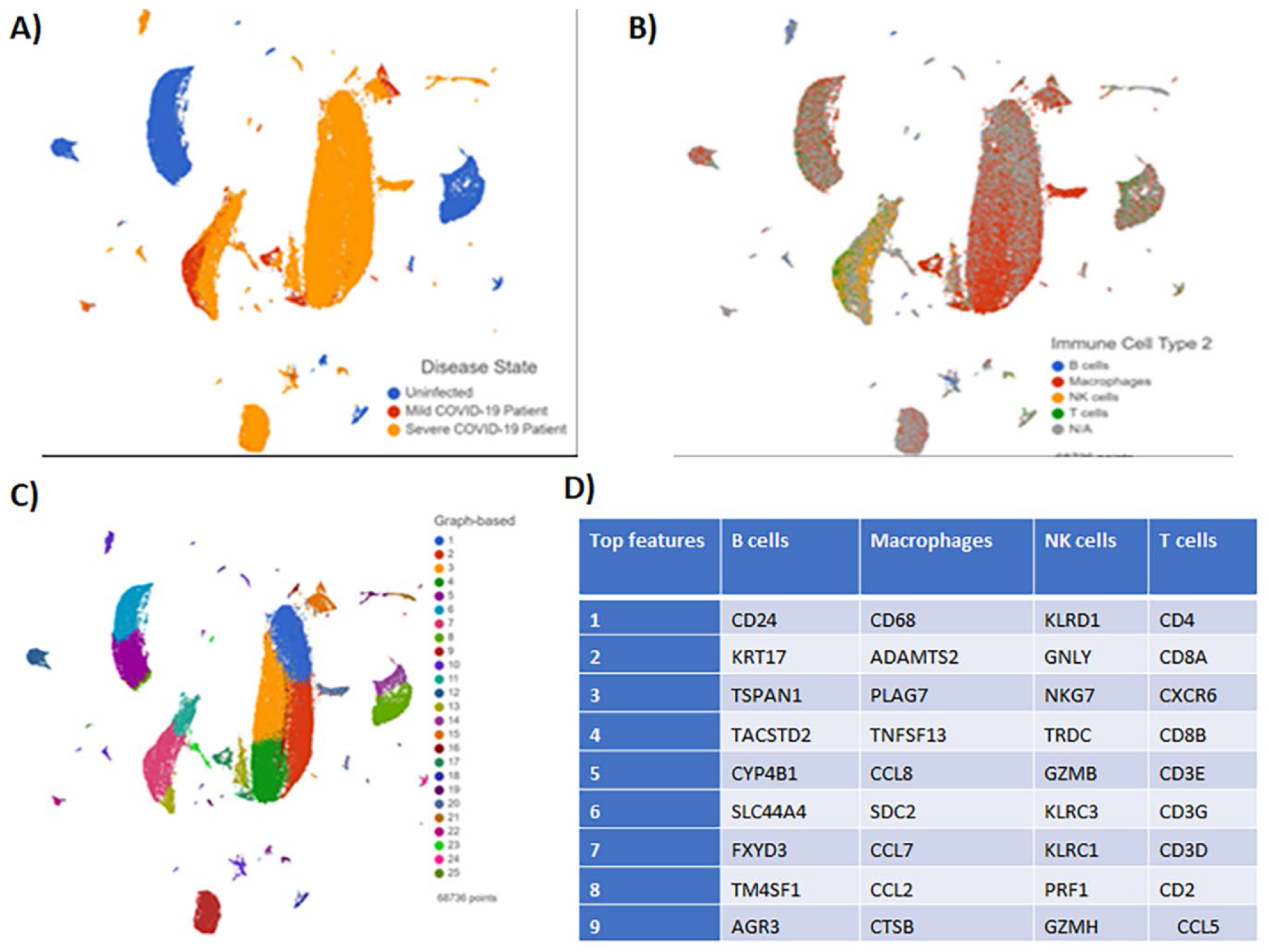
BALF Clustering Analysis results visualized by Global UMAP. **(A)** Patient samples were grouped by disease severity. **(B)** Cells were classified by four immune cell subsets: B-cells, Macrophages, NK cells, and T cells. N/A defines all unclassified cells. **(C)** Graph-based analysis was performed in *Partek® Flow®*. 25 clusters were reported. **(D)** Top features in each immune cell subset generated by clustering analysis.

### Differentially expressed genes in scRNA-seq of BALF in COVID-19 patients suggests immune markers of disease severity

Differentially expressed genes were identified by comparing infected versus healthy COVID-19 patients and severe versus mild COVID-19 patients to characterize immune patterns that contribute to disease severity. Genes were defined as significantly downregulated or upregulated if the threshold FDR value < 0.05, and fold change was ∓ 2. For several genes, the FDR value = 0. To determine the most significant values, genes with an FDR value = 0 were assigned the same FDR value as the next most significant value and sorted by the largest fold change. The 10xGenomics Human Immunology Panel was applied to the DEG output to filter and sort genes directly related to immunity.

The top ten significantly downregulated genes in infected patients were CD52, JAML, HLA-DPB1, TNFSF12, HLA-DQB2, HLA-DQB1, CES1, VMO1, MME, and CAMP (**Figure 7B**). The top ten significantly upregulated genes in infected patients were IFITM2, LAG3, SLAMF7, CCL3L1, CXCL10, IDO1, CCL2, CCL7, and CCL6 (**Figure 7B**). Three host factors (HLA-DPB1, HLA-DQB2, HLA-DQB1) were significantly downregulated, and several chemokines (CCL3L1, CXCL10, CCL2, CCL7, CCL6) were upregulated. It is well known that SARS-CoV-2 infection triggers high cytokine and chemokine levels. This result was consistent with observations made in previous studies, however, the role of HLA genes is not well established. HLA genes are human MHC II molecules, and their primary function is antigen presentation. A previous study showed downregulation of HLA class II proteins to be especially prominent in severe COVID-19 patients dependent on ventilation^26^. Downregulation of HLA class II molecules may be a potential strategy employed by SARS-CoV-2 to evade elimination by the host immune response.

**Figure 7.**
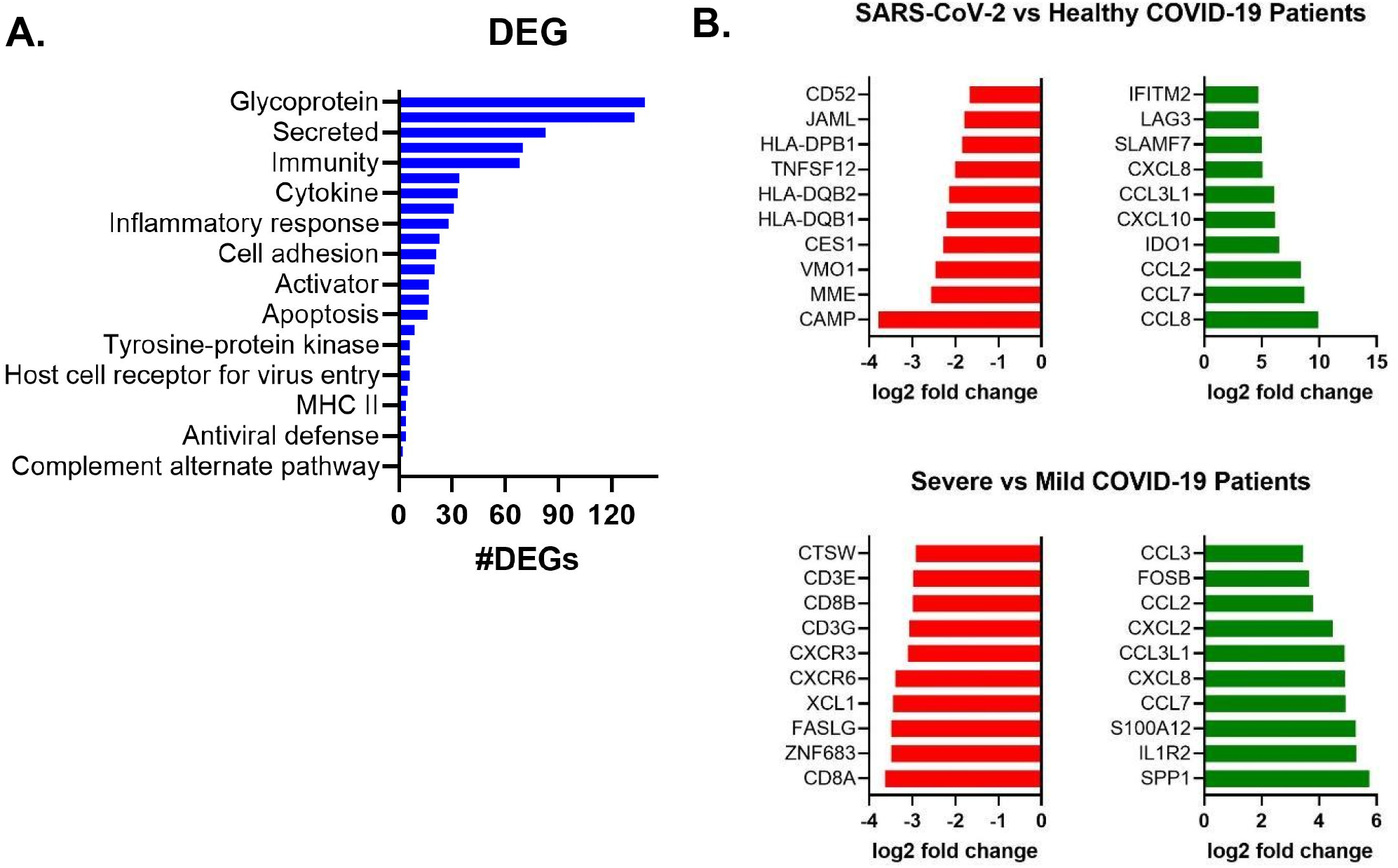
Differential gene expression analysis of RNAseq data from BALF of COVID-19 patients. **(A)** 10xGenomics Human Immunology Panel was used to filter differentially expressed genes for GSE145926 and annotated based on function. **(B**) The top row shows the DEGs for SARS-CoV-2 infected COVID-19 patients (mild+severe) versus healthy patients. The bottom row shows the DEGs for severe versus mild COVID-19 patients. The ten most significantly upregulated genes are ranked in green and the ten most significantly downregulated genes are ranked in red, based on log2(fold change).

The top ten significantly downregulated genes in infected patients were CTSW, CD3E, CD8B, CD3G, CXCR3, CXCR6, XCL1, FASLG, ZNF683, CD8A. The top ten significantly upregulated genes in infected patients were CCL3, FOSB, CCL2, CCL3L1, CXCL8, CCL7, S100A12, interleukin (IL)1R2, and SPP1. CCL7, CXCL8, CXCL3L1, and CCL2 were commonly upregulated between SARS-CoV-2 patients compared to healthy patients and when severe is compared to mild COVID-19 patients. This suggests that severe COVID-19 patients have a greater expression of chemokines. **Figure 7A** shows the 10Gx Genomics Human Immunology Panel gene annotation for function. All the DEGs were tallied and annotated according to the panel. The majority of DEGs in BALF are related to “glycoprotein” or “secreted” (**Figure 7A**). Glycoproteins are located on the cellular surface and can bind to receptors. This is important for innate sensing of pathogens, but more importantly, viruses bind to the host cell receptor to gain access. “Secreted” could correlate to secreted immunoglobulins, but not many of these genes were observed among the DEGs. Overall, these results indicate an activated innate immune response in the scRNA-seq profile of BALF collected from COVID-19 patients.

### scRNA-seq profile of BALF of severe vs. mild COVID-19 patients shows IKZF2 and HLA-DQA2 downregulated across immune cell subsets

To immunoprofile genes and characterize immune patterns that contribute to severity, we performed a differential gene analysis on each immune cell subset in severe vs. mild COVID-19 patients. 10xGenomics Human Immunology Panel was used to filter the gene directly related to immunity. A total of 18 genes for each immune cell subset was generated. Across all four immune cell subsets there was relatively the same level of upregulated genes in severe COVID-19 patients (**Figure 8A**). IKZF2 and HLA-DQA2 were commonly downregulated in all subsets, except for the B cells (**Figure 8A)**. HLA-DQA2 was the only downregulated gene in B cells **(Figure 8A)**. IKZF2 encodes a protein member of the Ikaros family of zinc-finger proteins, a group of transcription factors involved in regulating lymphocyte development. The downregulation of IKZF2 across all four immune cell subsets suggests hindered lymphocyte development in severe COVID-19 patients compared to mild COVID-19 patients. HLA-DQA2, a human class II MHC, was also commonly downregulated in NK cells, T cells, B cells, and macrophages. However, the expression seemed more pronounced in macrophages than the other immune cell subsets (**Figure 8C**.) HLA class II-DQ has been found to play crucial immunological roles, including antigen presentation to T lymphocytes and recognition of self and non-self-proteins^27^. Therefore, HLA polymorphisms could be associated with severe COVID-19.

**Figure 8.**
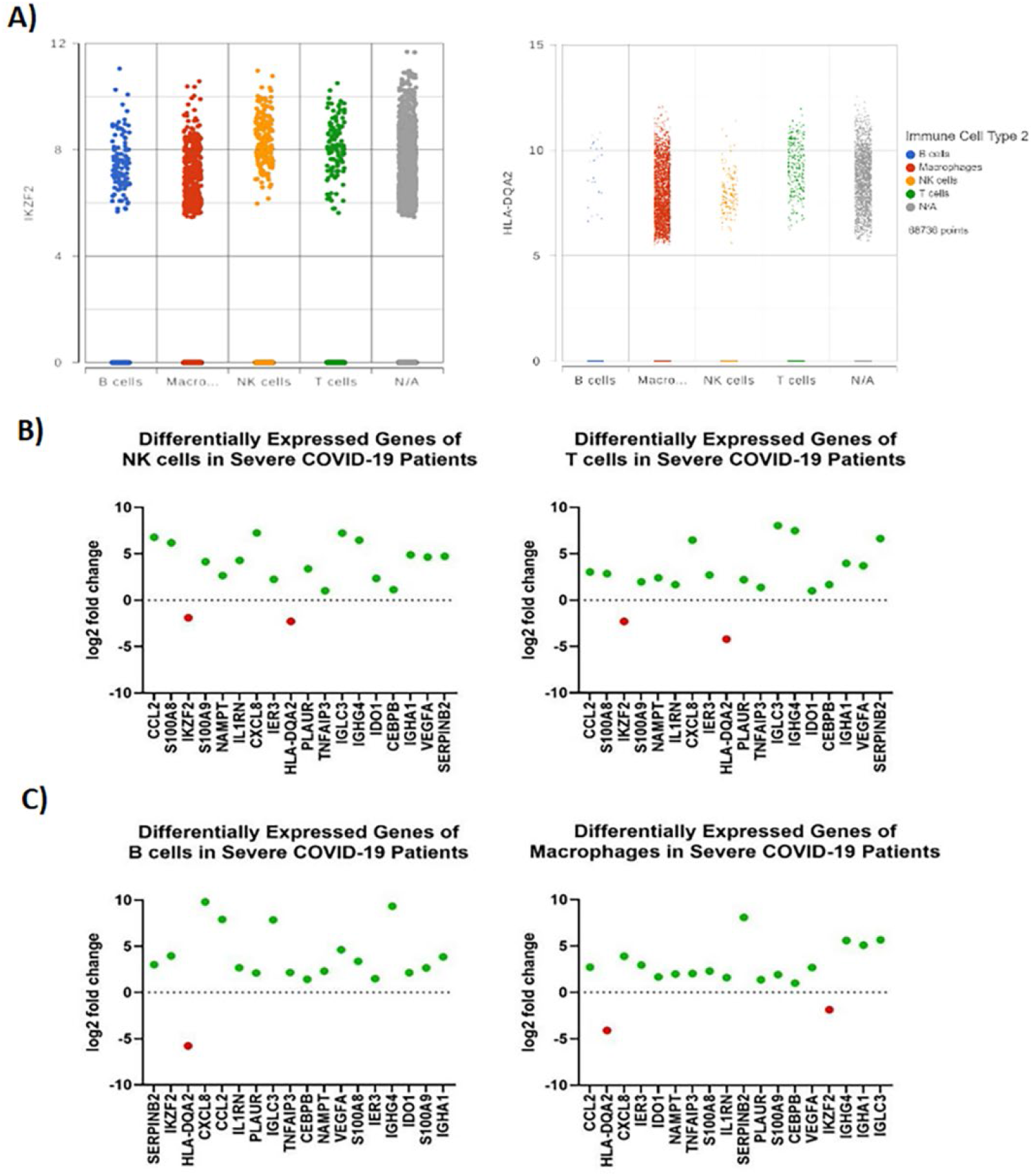
Differential gene expression analysis across immune cell subsets in RNAseq data from BALF of COVID-19 patients. **(A)** 10X Genomics Human Immunology Panel was used to filter differentially expressed genes in each immune subset (NK cells, T cells, B cells, and Macrophages) of severe versus mild COVID-19 patients. **(B)** Scatter plot for IKFZ2, one of the most downregulated genes, in immune cell subsets. **(C)** Scatter plot for HLA-DQA2, one of the most downregulated genes, in the immune cell subsets.

### Pathway analysis of differentially expressed genes in severe COVID-19 patients suggests an upregulated inflammatory response and inhibited Th1 response

Next, we applied IPA core analyses on DEGs for severe vs. mild patients to identify significant pathways and associated networks. The analysis ranked “TH1 Pathway’’ as the most significantly downregulated pathway and “TREM1” as the most upregulated pathway. A graphical summary was generated to provide an overview of the most predicted biological processes in the IPA core analysis (**Supplementary Figure 3**). T helper type 1 (Th1) cells are a lineage of CD4+ effector T cells that are responsible for cell-mediated immunity and protection against intracellular viral and bacterial pathogens^28^. CD4+ T cell responses are primarily based on their cytokine profile, producing interferon-gamma, interleukin (IL)-2, and tumor necrosis factor (TNF)-beta. This promotes neutrophil and monocyte-mediated inflammatory responses leading to phagocytosis^28^. Earlier studies demonstrated the downregulation of cytokines in SARS-CoV-2 infected patients. A reduction in IL-12 expression was observed in dendritic cells to quell the conversion of the Th1 cell phenotype, which is a vital type of cell-mediated immune response that is involved in viral clearance^29^.

Uncommon upregulation of cytokines and chemokines due to unsuccessful infection of dendritic cells can imbalance the stimulation of T-helper cell subsets by affecting the migrating and antigen-presenting function of dendritic cells to distort T-cell activation^29^. Our differential gene analysis indicates an upregulation in the expression of CCL2, an IFN-stimulated chemokine, as well as several other chemokines and cytokines. This suggests the immunological profile of SARS-CoV-2 patients includes a surge of chemokines and cytokines that could induce various immune responses, such as a helper T cell lymphopenia caused by a dysregulated Th1 response. TREM1 activation induces secretion of TNF-α, IL-6, IL-1β, IL-2, IL-12p40 by monocytes, macrophages, and dendritic cells, enhancing inflammation during infections by different pathogens^30^. This may potentially be another immunopathological event of severe COVID-19. Our IPA analysis predicts several upregulated biological processes related to myeloid cells, including chemotaxis of myeloid cells, cell movement of myeloid cells, a respiratory burst of myeloid cells, and the response myeloid cells, and inflammatory response (**Supplementary Figure 3**). Some of the cytokines involved are IFN-gamma, TNF, IL-6, IL-1A, IL-1B (**Supplementary Figure 3**). This suggests an upregulated inflammatory response due to chemokine and cytokine increase, but an inhibited Th1 response, resulting in a dysregulated immune response. Upregulation of the TREM1 pathway may comprise cellular and molecular mechanisms contributing to an impaired immune response.

### Clustering analysis and cell identification for the scRNA-seq profile of blood buffy coat reveals key differences in immune cell subsets in varying disease states

GSE154567 consisted of the scRNA-seq data for blood buffy coat samples of 12 SARS-CoV-2 infected patients and six convalescents. SARS-CoV-2 infected patients were labeled by disease severity, including five mild and six severe COVID-19 cases. We performed a graph-based cluster analysis to collate similar groups and classify four immune cell subsets (**Figure 9C**). Top biomarkers featured in the classification task were identified using ANOVA test with a threshold p<0.005. The output was used to verify the markers for cell identification. (**Figure 9D**). We compared COVID-19 patients by disease state and the classified immune cell groups, including NK cells, T cells, B cells, and macrophages (**Figure 9A, 9B**). A more significant number of NK cells and T-cells were observed in the cluster of mild COVID-19 patients than severe. There was a larger population of T-cells in the cluster of severe COVID-19 patients compared to mild. There were no large gaps in the number of counts per sample (**Supplemental Table 2**). This finding is still meaningful because it raises the question of what immune patterns appear for convalescent or recovering immunoprofile, which is not investigated thoroughly in the literature. It is unclear if the severe patients on the way to recovery or if the recovered patient was a recent severe patient. Again, careful immune monitoring during clinical studies could help uncover the cause and be considered for future studies.

**Figure 9.**
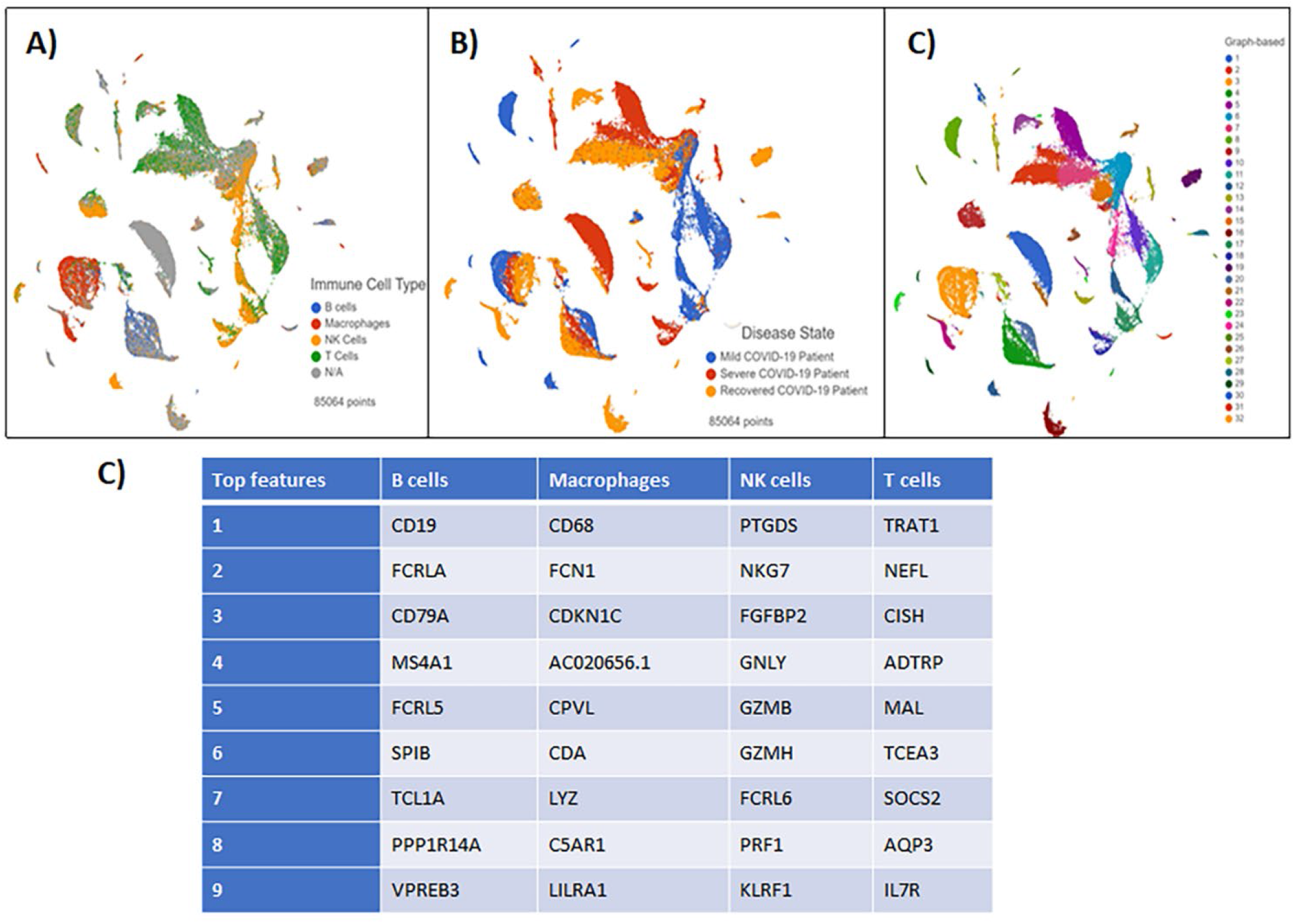
Blood Buffy Coat Clustering Analysis results visualized by Global UMAP. **(A)** Patient samples were grouped by disease severity. **(B)** Cells were classified by four immune cell subsets: B-cells, Macrophages, NK cells, and T cells. N/A defines all unclassified cells. **(C)** Graph-based analysis was performed in *Partek® Flow®*. 25 clusters were reported **(D)** The top features generated by clustering analysis in each immune cell subset.

### Differentially expressed genes in blood buffy coat samples reveals key genes that characterize immunological profiles of severe and recovered COVID-19 patients

To characterize immune patterns that contribute to disease severity differentially expressed genes were identified by comparing severe versus mild COVID-19 patients and convalescent versus severe COVID-19 patients. Differentially expressed genes (DEGs) were identified with ANOVA, defined as significantly downregulated or upregulated if the threshold FDR value < 0.05, and fold change was ∓ 2. The 10xGenomics Human Immunology Panel was applied to the DEG output to filter genes directly related to immunity.

Eight genes were differentially expressed in severe COVID-19 patients in comparison to mild COVID-19 patients: SLC25A37, SNCA, CXCL8, IFIT1, IFI27, HLA-DQA2, MEFV, ARG1 **(Figure 10**). CXCL8, IFIT1, and MEFV were significantly downregulated, and SLC25A37, SNCA, IFI27, HLA-DQA2, ARG1 were significantly upregulated (**Figure 10**). Conversely, the reverse is observed in recovered patients when compared to severe COVID-19 patients, in which CXCL8, IFIT1, and MEFV were upregulated, and SLC25A37, SNCA, IFI27 were downregulated (**Figure 10**). HLA-DQA2 and ARG1 were commonly upregulated in both severe and recovered COVID-19 patients (**Figure 10**). This suggests the immune patterns of CXCL8, IFIT1, MEFV, SLC25A37, SNCA, and IFI27 characterize the immunological profile of severe COVID-19 patients. CXCL8 and MEFV are involved in regulating the inflammatory response while IFIT1 and IFI27 are interferon-induced proteins involved in the antiviral response, reflecting an activation of these processes in severe COVID-19. SLC25A37 and SNCA are less clearly involved in the host immune response (**Supplementary Table 1**). Our analysis does not indicate these genes work in conjunction but are involved in several different processes that may overlap with host immune response.

**Figure 10.**
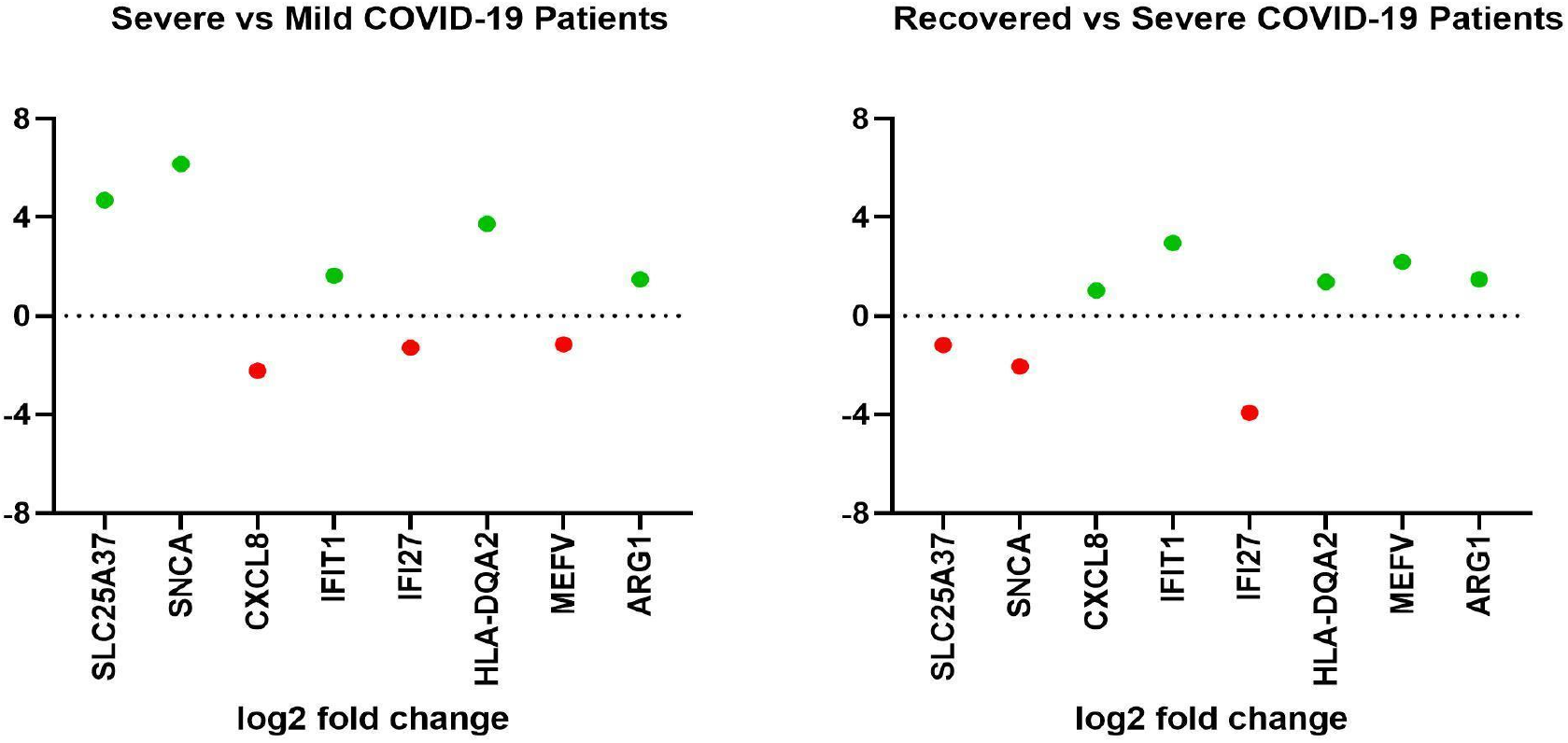
Differential gene expression of RNAseq data from blood buffy coat samples of severe and recovered COVID-19 patients. 10X Genomics Human Immunology Panel was used to filter differentially expressed genes of severe versus mild COVID-19 patients (left panel) and recovered versus severe COVID-19 patients (right panel).

### Differentially expressed genes in blood buffy coat samples reveals consistent upregulation of MX1, IFIT1, and HLA-DQA2 across immune cell subsets in severe COVID-19 patients

Next, we compared the DEGs in each immune cell subset and each disease state (mild, severe, recovered). Three genes were significantly upregulated across the immune cell subsets in severe COVID-19 patients compared to mild COVID-19 patients, MX1, IFIT1, and HLA-DQA2. The scatter plots of the gene expression changes are shown in **Supplementary Figure 4**. Expression across the immune cell subsets was relatively similar for the three genes, however, IFIT1 and MX1 were more enriched in severe COVID-19 patients than mild or recovered patients. HLA-DQA2 is more enriched in recovered COVID-19 patients. This suggests an immune pattern for severe patients’ immunological profile, involving the upregulation of IFIT1 and MX1 genes. Regained expression of the host factor, HLA-DQA2, may characterize the immunological profile of recovered patients. A total of 10 genes for each immune cell subset were generated in recovered patients compared to severe patients. (**Figure 11**). Across all four immune cell subsets, there was relatively the same level of upregulated genes and downregulated in severe COVID-19 patients (**Figure 11)**. Our analyses for severe COVID-19 patients suggest that upregulated interferon-induced proteins, chemokines, and cytokines contribute to the severity of COVID-19 compared to mild patients. It remains unclear what immune patterns the immunological profile of a recovered patient may entail. The recovered patients may retain some of the immune characteristics in early recovery or perhaps a complete reversal of the immunological gene signature. There may be stages of recovery and immunological gene signatures that define the progression of recovery. This requires further investigation to understand how the host immune response overcomes severe COVID-19.

**Figure 11.**
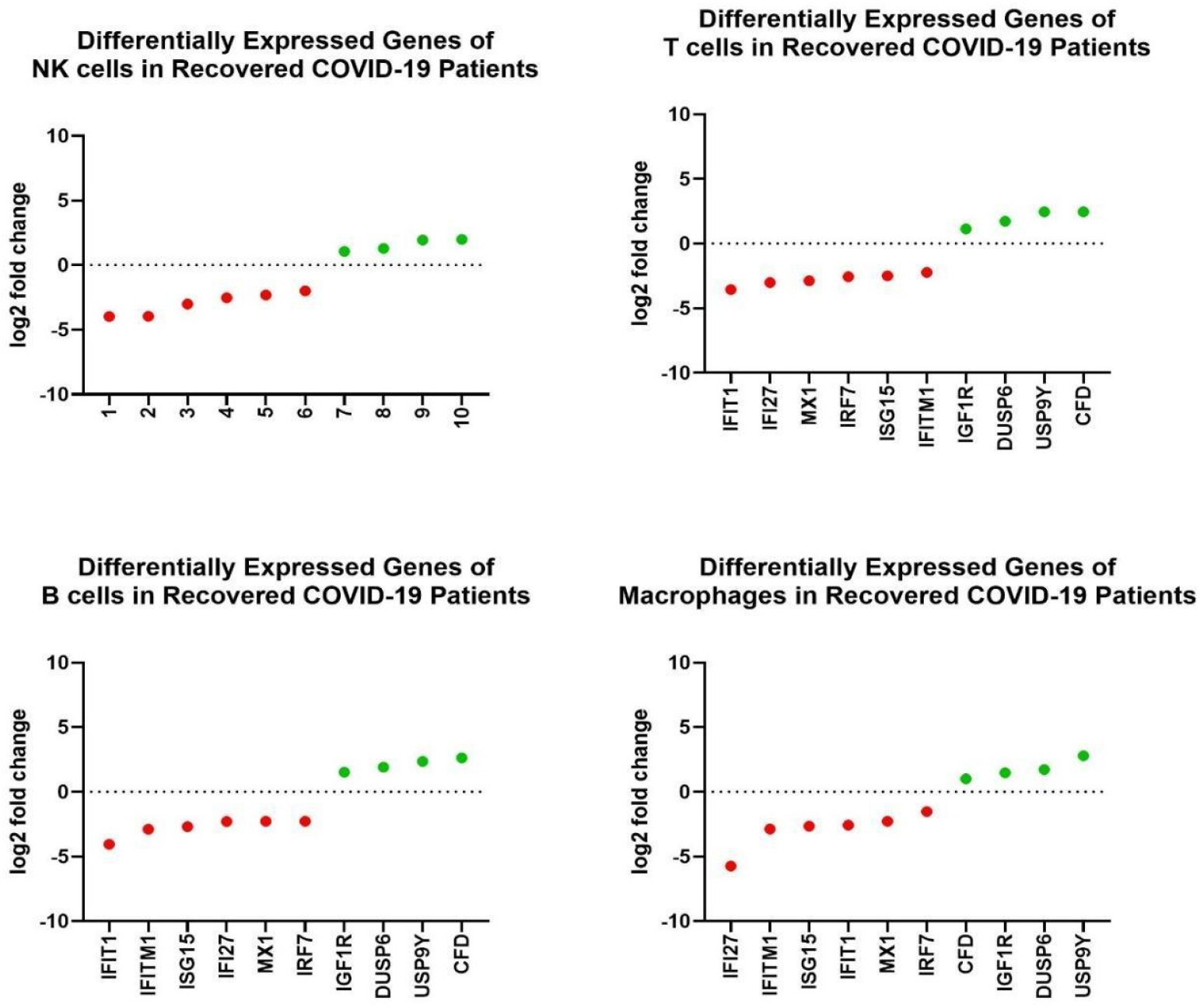
Differential gene expression of RNAseq data from blood buffy coat of recovered COVID-19 patients. Differentially expressed genes across immune cell subsets (NK cells, T cells, B cells, and macrophages) found in recovered versus severe COVID-19 patients.

### Pathway analysis reveals upregulation of MSP-RON signaling and arginine degradation pathways in severe COVID-19 patients

Next, we applied IPA core analyses on DEGs for severe vs. mild patients to identify significant pathways and associated networks. The analysis did not demonstrate any predictions of activation or inhibition; however, MSP-RON Signaling in Macrophages and Arginine Degradation I (Arginase Pathway) were ranked as the top two most significant (p-value <0.05). Macrophage-stimulating protein (MSP), is the only known ligand for recepteur d’origine nantais (RON), a member of the MET proto-oncogene family. Arginine is an amino acid shown in non-clinical studies to be essential in many viruses’ life cycle^31^. As such, arginine depletion may represent a practical therapeutic approach against SARS-CoV-2^31^.

### Comparison across data sets reveals significant pathways affected consistently in COVID-19 patient samples

A comparison analysis was applied to all datasets in IPA to identify common significant pathways across all four datasets. A heat map of the analysis results was generated with IPA Canonical Pathways (**Supplementary Figure 6**). Hepatic Fibrosis Signaling, Osteoarthritis, and Neuroinflammation pathways were associated with activation while Interferon Signaling was inhibited among severe COVID-19 patients. This suggests that these pathways contribute to the severity of COVID-19 disease. Further investigation of each pathway and active molecules should be performed in future studies.

### Upstream immune regulators present potential therapeutic targets for COVID-19

We next sought to identify potential therapeutic targets for COVID-19 based on key genes within the most significant pathways we identified. We applied the “Regulator Effects” function in IPA onto all the pathway analyses to determine the biological impact of upstream molecules according to the genes they regulate. A list of 547 upstream regulators and associated target molecules were generated from the analysis (**Supplementary Table 3**). However, these targets and corresponding drugs require further analysis in our future studies. As previously noted, our scRNA-seq differential gene analyses were filtered based on the 10xGenomics Human Immunology Panel. Therefore, there are several DEGs that were not included in our queries that may not directly relate to the immune system but may also be potential targets. Genes that were noted as the topmost significantly expressed in this study was determined by an FDR value of <0.01, but not by fold-change. It is possible that several statistically significant genes that may exhibit notable variations in gene expression were missed during the analysis. Although our list requires further analysis to determine druggability of each target, many existing drugs have potential to target these key regulators involved in COVID-19 pathogenesis, which is encouraging for drug repurposing efforts or novel therapeutics discovery. We propose many potential host immune therapeutic targets for COVID-19 that our results show have the most significant impact on the gene expression profiles observed in COVID-19 patients.

## Conclusion

In this study, we analyzed the RNA seq-profiles of nasopharyngeal swabs, PBMCs, BALF, and the blood buffy coats collected from COVID-19 patients. We performed differential gene analyses and pathway analyses to characterize the host immune response and severe COVID-19 patients’ immunological profile. Our results showed various immune responses that are consistent with those reported in other studies, including increased action of interferon-induced proteins and increased chemokine/cytokine production in SARS-CoV-2 infected patients^6,10,15^. We observed interferon-alpha to be a major upstream regulator of ISG15 and IFITM-family proteins, important for antiviral defense and adaptive immunity, as well as proteins encoding cell metabolism in immunity. As previously reported, we find that interferon alpha plays a major role in antiviral host defense but over-activation may contribute to the hyperinflammatory responses reported in cases of COVID-19 patients.

IPA pathway analysis identified several pathways significantly affected in COVID-19 patients, including multiple pathways involved in cell cycle regulation (**Figure 4**). These findings suggest that host cells employ defense mechanisms that are either directly involved in or overlap with cell division processes to counteract viral replication and release. There is potential for genes involved in cell division pathways to be adversely modulated in SARS-CoV-2 infection and lead to cancer. SARS-CoV-2 may also directly be involved in mechanisms of oncogenesis. Based on the gene expression changes and significant pathways indicated in our analysis, we propose one potential mechanism by which SARS-CoV-2 may dysregulate the cell cycle through interaction with the Rb-E2F complex, leading to cell proliferation and potentially contributing to oncogenesis (**Figure 5**). Further investigation is necessary to examine the relationship between SARS-CoV-2 infection and the cell cycle in order to elucidate a mechanism for oncogenesis as a result of viral-induced expression changes. It is critical to investigate the potential connection between SARS-CoV-2 infection and cancer. The process of oncogenesis could begin undetected in COVID-19 patients and not become apparent for many years after infection, at which point intervention may come too late.

In our analysis, many differentially expressed genes that were seemingly non-immune related genes were downregulated. For example, DHRS9, a type of dehydrogenase involved in metabolic processes, was one of the most significantly downregulated genes in SARS-CoV-2 patients. This suggests that immune pathways or processes may overlap with metabolic processes during a host immune response. This result was further emphasized in pathway analysis that ranked Mitochondrial Dysfunction and Oxidative Phosphorylation as significantly downregulated in COVID-19 patients. Previous studies report lymphopenia as part of the immunopathology of COVID-19. This lymphopenia may be connected to dysfunctional cell metabolism of different immune effector cells. Recent gene expression studies have also shown evidence for metabolic changes in COVID-19 patients^32,33^. However, few studies have investigated how metabolic processes might be correlated to the host immune response to SARS-CoV-2 infection.

Lastly, a major component of our analysis was examining immune cell subsets in severe COVID-19 patients compared to mild or recovered COVID-19 patients. HLA gene expression was commonly modulated in the two scRNA-seq profiles we analyzed. These are important host factors that are affected in SARS-CoV-2 infection and further investigation of the HLA polymorphisms may provide important insight about the antiviral host defense and susceptibility to severe COVID-19 in patients.

Overall, this study affirms that immune characteristics can be potential biomarkers or therapeutic targets for severe disease. The differentially expressed genes and significant pathways found in this analysis should be further investigated for novel understanding of the mechanisms underlying the host response to SARS-CoV-2, as well as selecting potential therapeutic targets and drug candidates for COVID-19.

## Acknowledgments

The author Sivanesan Dakshanamurthy wishes to acknowledge the support by Georgetown University Medical Center COVID19 Pilot Grant, CCSG grant P30 CA051008/CA/NCI NIH HHS/United States, GUMC Lombardi Comprehensive Cancer Center, and GUMC Computational Chemistry Shared Resources (CCSR).

## Supplemental Information

**Supplementary Figure 1**. Network map associated with upregulated canonical pathways from GSE152075 shows interferon alpha as a key regulator.

**Supplementary Figure 2**. Network map associated shows relationships among significant canonical pathways from GSE152418.

**Supplementary Figure 3**. Network map shows significant biological processes affected in brochoalveolar lung fluid of severe COVID-19 patients.

**Supplementary Figure 4**. 10X Genomics Human Immunology Panel filtered differentially expressed genes in each immune subset (NK cells, T cells, B cells, and Macrophages) of severe versus mild COVID-19 patients.

**Supplementary Figure 5**. Network map shows interactions between differentially expressed genes in severe versus mild COVID-19 patients.

**Supplementary Figure 6**. Comparison analysis across the four analyzed datasets reveals canonical pathways consistently significant in COVID-19 patient samples.

**Supplementary Table 1**. Descriptions of relevant genes mentioned in paper.

**Supplementary Table 2**. Classification counts for GSE145926 quantifies immune cell populations in each BALF sample.

**Supplementary Table 3**. Classification counts for GSE154567 quantifies immune cell populations in each blood buffy coat sample.

**Supplementary Table 4**. Upstream regulators of significant dataset molecules present potential host therapeutic targets for COVID-19.

## References

[1] Huang, C., Wang, Y., Li, X., Ren, L., Zhao, J., Hu, Y., Zhang, L., Fan, G., Xu, J., Gu, X., Cheng, Z., Yu, T., Xia, J., Wei, Y., Wu, W., Xie, X., Yin, W., Li, H., Liu, M., Xiao, Y.,… Cao, B. (2020). Clinical features of patients infected with 2019 novel coronavirus in Wuhan, China. Lancet (London, England), 395(10223), 497–506. https://doi.org/10.1016/S0140-6736(20)30183-5

[2] WHO. Weekly epidemiological update - 24 November 2020. (2020). Retrieved December 08, 2020, from https://www.who.int/publications/m/item/weekly-epidemiological-update---24-november-2020

[3] WHO coronavirus disease (COVID-19) dashboard. Geneva: World Health Organization, 2020. Retrieved February 14, 2021 from https://covid19.who.int/

[4] Kalil, A. C., Patterson, T. F., Mehta, A. K., Tomashek, K. M., Wolfe, C. R., Ghazaryan, V., Marconi, V. C., Ruiz-Palacios, G. M., Hsieh, L., Kline, S., Tapson, V., Iovine, N. M., Jain, M. K., Sweeney, D. A., El Sahly, H. M., Branche, A. R., Regalado Pineda, J., Lye, D. C., Sandkovsky, U.,… Beigel, J. H. (2020). Baricitinib plus Remdesivir for hospitalized adults with COVID-19. New England Journal of Medicine, 0(0), null. https://doi.org/10.1056/NEJMoa2031994

[5] WHO recommends against the use of remdesivir in COVID-19 patients. (2020). Retrieved December 08, 2020, from https://www.who.int/news-room/feature-stories/detail/who-recommends-against-the-use-of-remdesivir-in-covid-19-patients

[6] Yang, L., Liu, S., Liu, J., Zhang, Z., Wan, X., Huang, B., Chen, Y., & Zhang, Y. (2020). COVID-19: Immunopathogenesis and immunotherapeutics. Signal Transduction and Targeted Therapy, 5(1), 128. https://doi.org/10.1038/s41392-020-00243-2

[7] Weiss, S. R., & Leibowitz, J. L. (2011). Coronavirus pathogenesis. Advances in Virus Research, 81, 85–164. https://doi.org/10.1016/B978-0-12-385885-6.00009-2

[8] Li, G., Fan, Y., Lai, Y., Han, T., Li, Z., Zhou, P., Pan, P., Wang, W., Hu, D., Liu, X., Zhang, Q., & Wu, J. (2020). Coronavirus infections and immune responses. Journal of Medical Virology, 92(4), 424–432. https://doi.org/10.1002/jmv.25685

[9] Dhama, K., Khan, S., Tiwari, R., Sircar, S., Bhat, S., Malik, Y. S., Singh, K. P., Chaicumpa, W., Bonilla-Aldana, D. K., & Rodriguez-Morales, A. J. (2020). Coronavirus Disease 2019-COVID-19. Clinical Microbiology Reviews, 33(4), e00028–20. https://doi.org/10.1128/CMR.00028-20

[10] Mortaz, E., Tabarsi, P., Varahram, M., Folkerts, G., & Adcock, I. M. (2020). The Immune Response and Immunopathology of COVID-19. Frontiers in Immunology, 11, 2037. https://doi.org/10.3389/fimmu.2020.02037

[11] Ovsyannikova, I. G., Haralambieva, I. H., Crooke, S. N., Poland, G. A., & Kennedy, R. B. (2020). The role of host genetics in the immune response to SARS-CoV-2 and COVID-19 susceptibility and severity. Immunological Reviews, 296(1), 205–219. https://doi.org/10.1111/imr.12897

[12] Brodin, P. (2021) Immune determinants of COVID-19 disease presentation and severity. Nature Medicine, 27(1), 28–33. https://doi.org/10.1038/s41591-020-01202-8. in Ingenuity Pathway Analysis. Bioinformatics. (2014). 30(4), 523–30.

[13] Tay, M. Z., Poh, C. M., Rénia, L., MacAry, P. A., & Ng, L. (2020). The trinity of COVID-19: immunity, inflammation and intervention. Nature reviews. Immunology, 20(6), 363–374. https://doi.org/10.1038/s41577-020-0311-8

[14] Miorin, L., Kehrer, T., Sanchez-Aparicio, M. T., Zhang, K., Cohen, P., Patel, R. S., Cupic, A., Makio, T., Mei, M., Moreno, E., Danziger, O., White, K. M., Rathnasinghe, R., Uccellini, M., Gao, S., Aydillo, T., Mena, I., Yin, X., Martin-Sancho, L.,… García-Sastre, A. (2020). SARS-CoV-2 Orf6 hijacks Nup98 to block STAT nuclear import and antagonize interferon signaling. Proceedings of the National Academy of Sciences, 117(45), 28344–28354. https://doi.org/10.1073/pnas.2016650117

[15] Vabret, N., Britton, G. J., Gruber, C., Hegde, S., Kim, J., Kuksin, M., Levantovsky, R., Malle, L., Moreira, A., Park, M. D., Pia, L., Risson, E., Saffern, M., Salomé, B., Esai Selvan, M., Spindler, M. P., Tan, J., van der Heide, V., Gregory, J. K., Alexandropoulos, K.,… Sinai Immunology Review Project (2020). Immunology of COVID-19: Current State of the Science. Immunity, 52(6), 910–941. https://doi.org/10.1016/j.immuni.2020.05.002

[16] Barrett, T., Wilhite, S. E., Ledoux, P., Evangelista, C., Kim, I. F., Tomashevsky, M., Marshall, K. A., Phillippy, K. H., Sherman, P. M., Holko, M., Yefanov, A., Lee, H., Zhang, N., Robertson, C. L., Serova, N., Davis, S., & Soboleva, A. (2013) NCBI GEO: archive for functional genomics data sets--update. Nucleic Acids Research, 41(Database issue):D991–5.

[17] Fung, T. S., Liao, Y., & Liu, D. X. (2016). Regulation of Stress Responses and Translational Control by Coronavirus. Viruses, 8(7), 184. https://doi.org/10.3390/v8070184

[18] Kindler, E., Gil-Cruz, C., Spanier, J., Li, Y., Wilhelm, J., Rabouw, H. H., Züst, R., Hwang, M., V’kovski, P., Stalder, H., Marti, S., Habjan, M., Cervantes-Barragan, L., Elliot, R., Karl, N., Gaughan, C., van Kuppeveld, F. J., Silverman, R. H., Keller, M., Ludewig, B.,… Thiel, V. (2017). Early endonuclease-mediated evasion of RNA sensing ensures efficient coronavirus replication. PLoS Pathogens, 13(2), e1006195. https://doi.org/10.1371/journal.ppat.1006195

[19] Koc, E. C., Burkhart, W., Blackburn, K., Moyer, M. B., Schlatzer, D. M., Moseley, A., & Spremulli, L. L. (2001). The large subunit of the mammalian mitochondrial ribosome: Analysis of the complement of ribosomal proteins present. Journal of Biological Chemistry, 276(47), 43958– 43969. https://doi.org/10.1074/jbc.M106510200

[20] Angajala, A., Lim, S., Phillips, J. B., Kim, J. H., Yates, C., You, Z., & Tan, M. (2018). Diverse Roles of Mitochondria in Immune Responses: Novel Insights Into Immuno-Metabolism. Frontiers in Immunology, 9, 1605. https://doi.org/10.3389/fimmu.2018.01605

[21] Jafarzadeh, A., Nemati, M., Saha, B., Bansode, Y. D., & Jafarzadeh, S. (2020). Protective Potentials of Type III Interferons in COVID-19 Patients: Lessons from Differential Properties of Type I-and III Interferons. Viral Immunology. https://doi.org/10.1089/vim.2020.0076

[22] Greco, S., Madè, A., Gaetano, C., Devaux, Y., Emanueli, C., & Martelli, F. (2020). Noncoding RNAs implication in cardiovascular diseases in the COVID-19 era. Journal of Translational Medicine, 18(1), 408. https://doi.org/10.1186/s12967-020-02582-8

[23] Riquelme, P., Amodio, G., Macedo, C., Moreau, A., Obermajer, N., Brochhausen, C., Ahrens, N., Kekarainen, T., Fändrich, F., Cuturi, C., Gregori, S., Metes, D., Schlitt, H. J., Thomson, A. W., Geissler, E. K., & Hutchinson, J. A. (2017). DHRS9 Is a Stable Marker of Human Regulatory Macrophages. Transplantation, 101(11), 2731–2738. https://doi.org/10.1097/TP.0000000000001814

[24] Nevins, J. R. (2001). The Rb/E2F pathway and cancer. Human molecular genetics, 10(7), 699–703. https://doi.org/10.1093/hmg/10.7.699

[25] Merad, M., & Martin, J. C. (2020). Pathological inflammation in patients with COVID-19: a key role for monocytes and macrophages. Nature reviews. Immunology, 20(6), 355–362. https://doi.org/10.1038/s41577-020-0331-4

[26] Wilk, A. J., Rustagi, A., Zhao, N. Q., Roque, J., Martínez-Colón, G. J., McKechnie, J. L., Ivison, G. T., Ranganath, T., Vergara, R., Hollis, T., Simpson, L. J., Grant, P., Subramanian, A., Rogers, A. J., & Blish, C. A. (2020). A single-cell atlas of the peripheral immune response in patients with severe COVID-19. Nature Medicine, 26(7), 1070–1076. https://doi.org/10.1038/s41591-020-0944-y

[27] Lorente, L., Martín, M. M., Franco, A., Barrios, Y., Cáceres, J. J., Solé-Violán, J., Perez, A., Marcos Y Ramos, J.A., Ramos-Gómez, L., Ojeda, N., Jiménez, A., Working Group on COVID-19 Canary ICU, & Annex. Members of the BIOMEPOC group (2020). HLA genetic polymorphisms and prognosis of patients with COVID-19. Medicina Intensiva, S0210-5691(20)30266-7. Advance online publication. https://doi.org/10.1016/j.medin.2020.08.004

[28] Romagnani S. (2000). T-cell subsets (Th1 versus Th2). Annals of allergy, asthma & immunology: official publication of the American College of Allergy, Asthma, & Immunology, 85(1), 9–21. https://doi.org/10.1016/S1081-1206(10)62426-X

[29] Zhang, Y. Y., Li, B. R., & Ning, B. T. (2020). The Comparative Immunological Characteristics of SARS-CoV, MERS-CoV, and SARS-CoV-2 Coronavirus Infections. Frontiers in Immunology, 11, 2033. https://doi.org/10.3389/fimmu.2020.02033

[30] Arts, R.J.W., Joosten, L.A.B., van der Meer, J.W.M. and Netea, M.G. (2013), TREM-1: intracellular signaling pathways and interaction with pattern recognition receptors. Journal of Leukocyte Biology, 93: 209–215. https://doi.org/10.1189/jlb.0312145

[31] Grimes, J. M., Khan, S., Badeaux, M., Rao, R. M., Rowlinson, S. W., & Carvajal, R. D. (2021). Arginine depletion as a therapeutic approach for patients with COVID-19. International Journal of Infectious Diseases. 102, 566–570. https://doi.org/10.1016/j.ijid.2020.10.100

[32] Gardinassi, L. G., Souza, C. O. S., Sales-Campos, H., & Fonseca, S. G. (2020). Immune and Metabolic Signatures of COVID-19 Revealed by Transcriptomics Data Reuse. Frontiers in Immunology, 11. https://doi.org/10.3389/fimmu.2020.01636

[33] Moolamalla, S. T. R., Chauhan, R., Deva Priyakumar, U., & Vinod, P. K. (2020). Host metabolic reprogramming in response to SARS-Cov-2 infection [Preprint]. Bioinformatics. https://doi.org/10.1101/2020.08.02.232645

