## Supplementary Materials for "Immune characterization and profiles of SARS-CoV-2 infected patients reveals potential host therapeutic targets and SARS-CoV-2 oncogenesis mechanism"

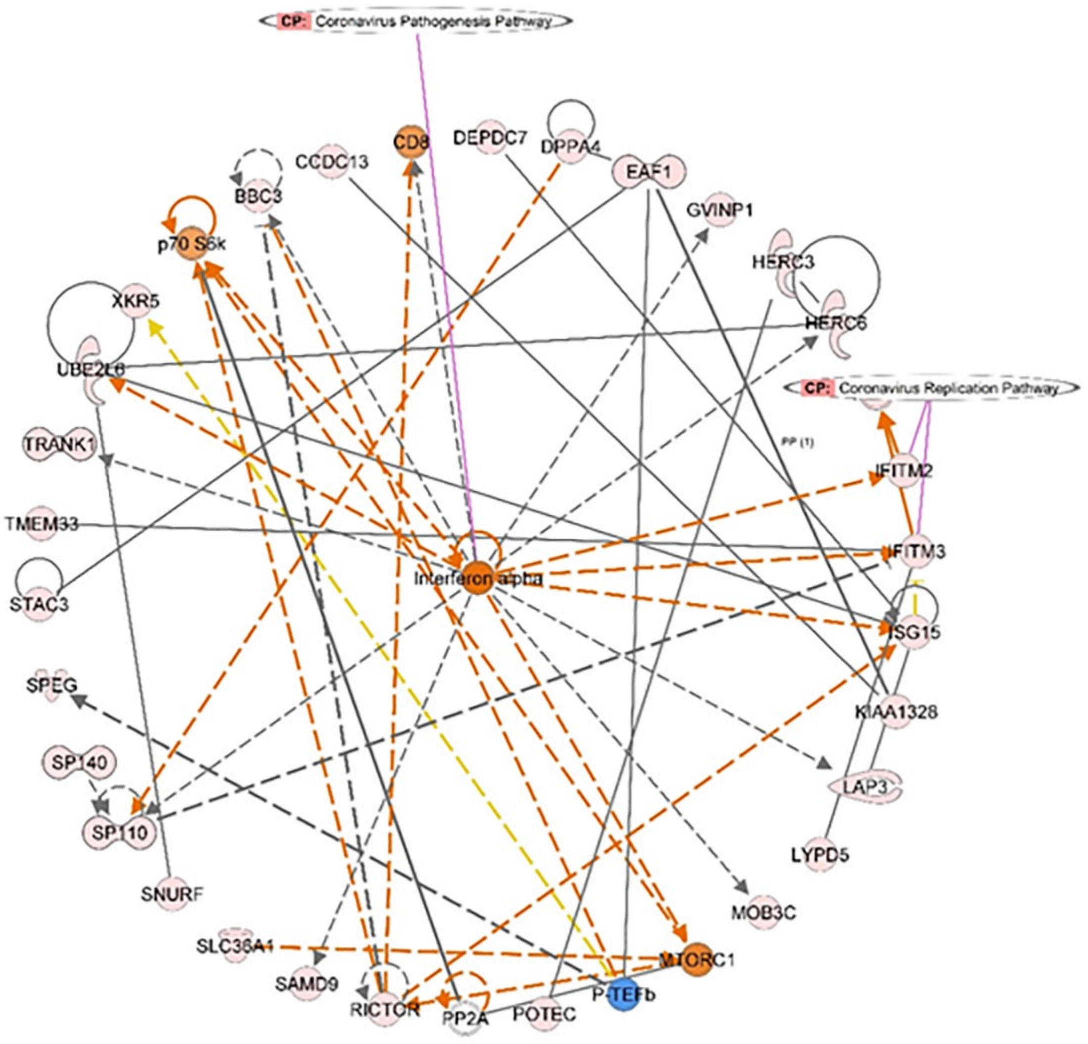


**Supplementary Figure 1. Network map associated with upregulated canonical pathways shows interferon alpha as a key regulator.** IPA core analysis determined interferon-alpha as an upstream regulator in the significantly upregulated genes from RNAseq data from nasopharyngeal swabs of COVID-19 patients (GSE152075). Network map was generated in IPA, overlaid with the Coronavirus Replication Pathway.

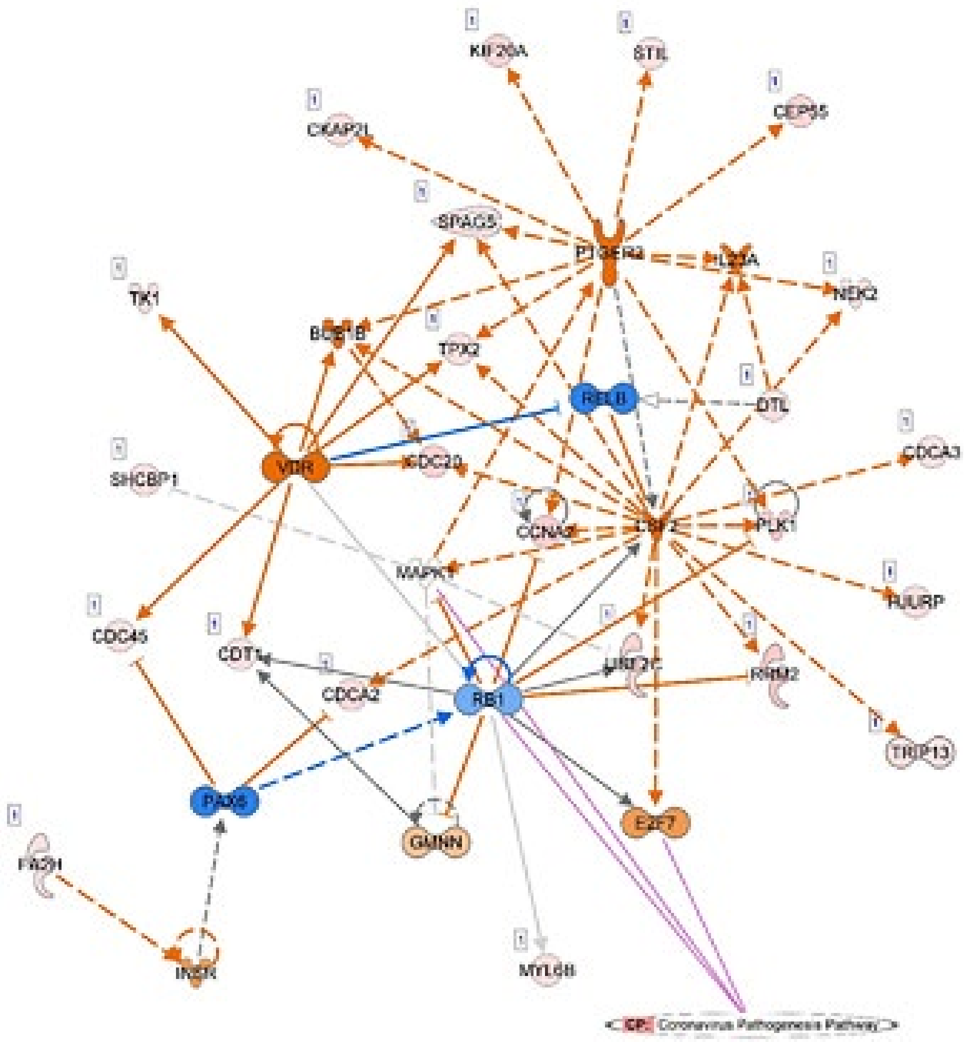


**Supplementary Figure 2. Network map associated with Cell Cycle, Cellular Assembly and Organization, DNA Replication, Recombination, and Repair shows relationships among significant canonical pathways.** Significant pathways were identified from pathway analysis of RNAseq from PBMCs of COVID-19 patients. Coronavirus Pathogenesis Pathway was also overlaid on the network map. The orange and blue colors indicate predicted activation or predicted inhibition, respectively.

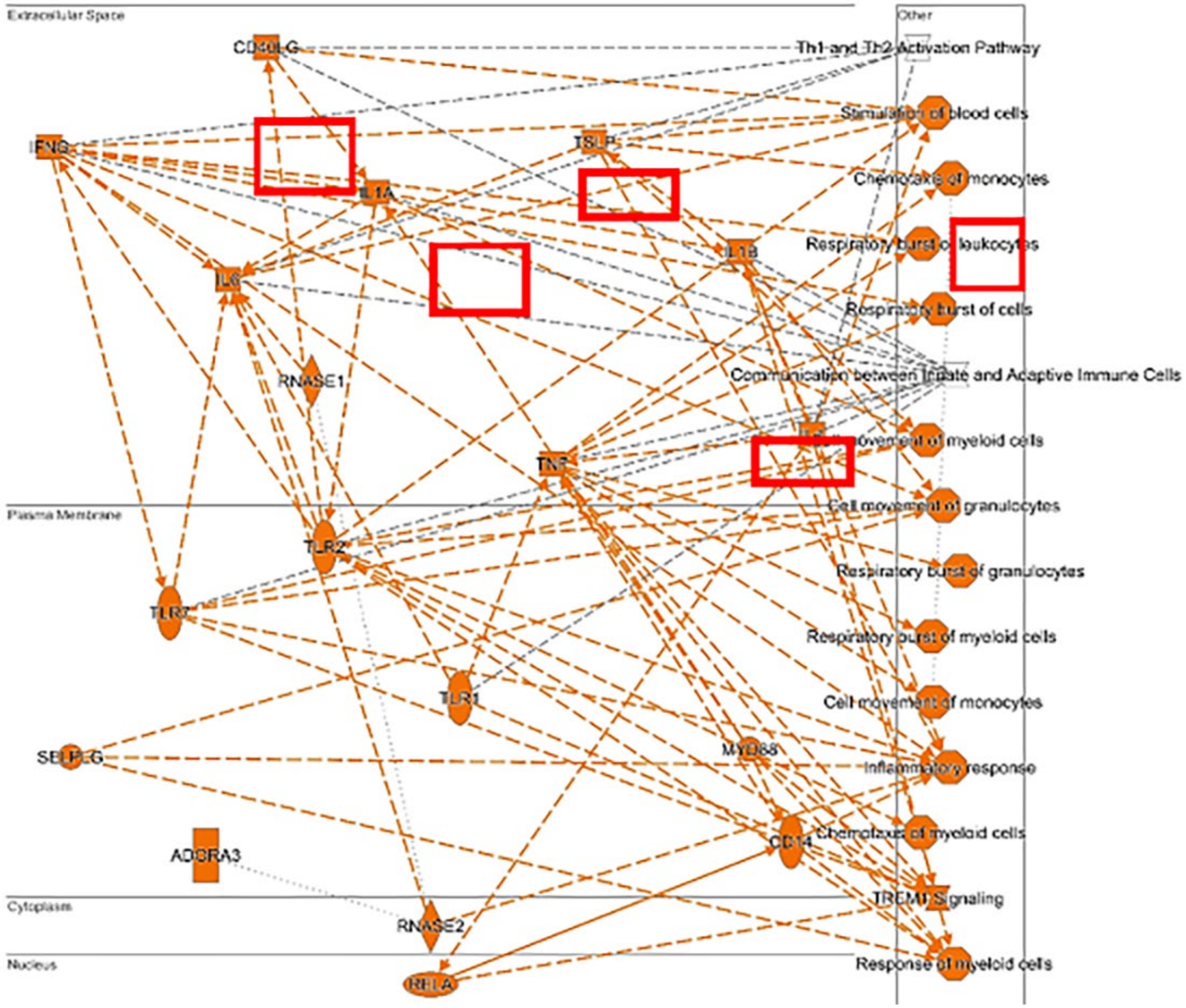


**Supplementary Figure 3. Significant biological processes affected in brochoalveolar lung fluid of severe COVID-19 patients.** Network map was generated by IPA core analysis of differentially expressed genes for severe vs mild COVID-19 patients in bronchoalveolar lung fluid (BALF) from scRNA-seq profile of GSE145926. Orange color represents predicted activation. Red boxes highlight important cytokines involved.

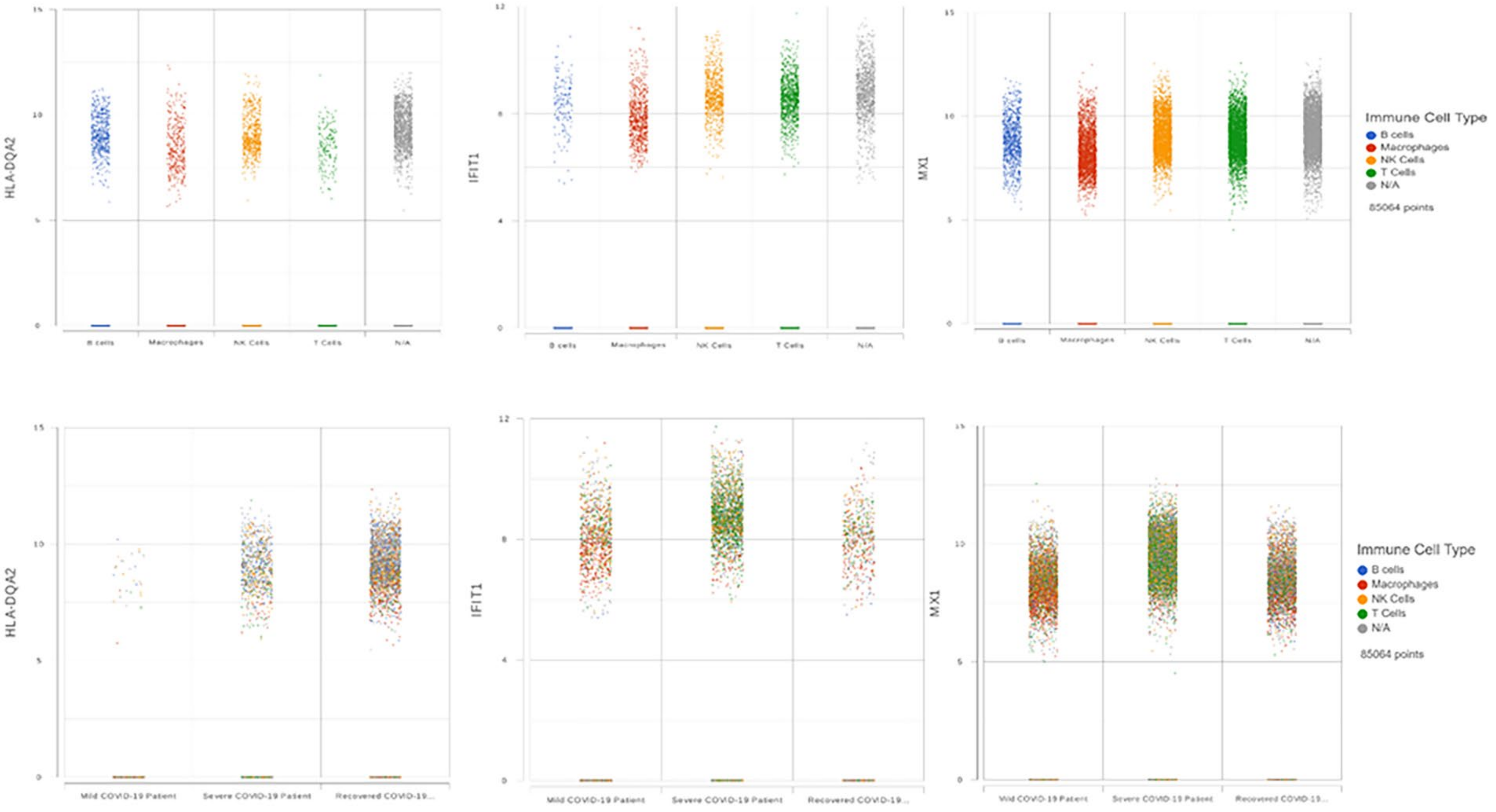


**Supplementary Figure 4. 10X Genomics Human Immunology Panel filtered differentially expressed genes in each immune subset (NK cells, T cells, B cells, and Macrophages) of severe versus mild COVID-19 patients.** Three genes (HLA-DQA2, IFIT1, and MX1) were found significantly and consistently differentially expressed. Gene expression is shown per the disease severity (mild, severe, recovered) is shown on the top row and expression across immune cell subsets are shown on the bottom row.

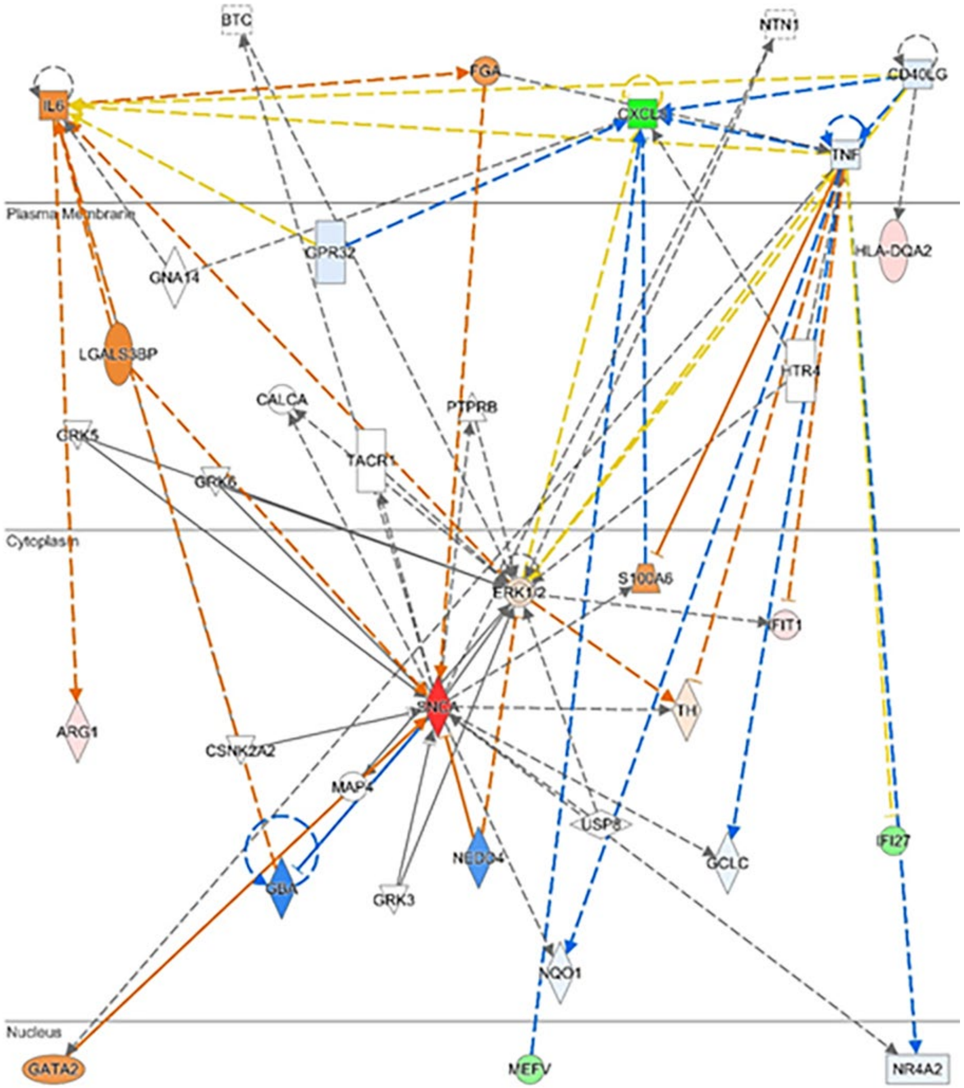


**Supplementary Figure 5. Network map shows interactions between differentially expressed genes in severe versus mild COVID-19 patients.** IPA was used to generate this plot based on scRNA-seq data from blood buffy coat samples of GSE154567. Orange color represents predicted activation and blue represents predicated inhibition.

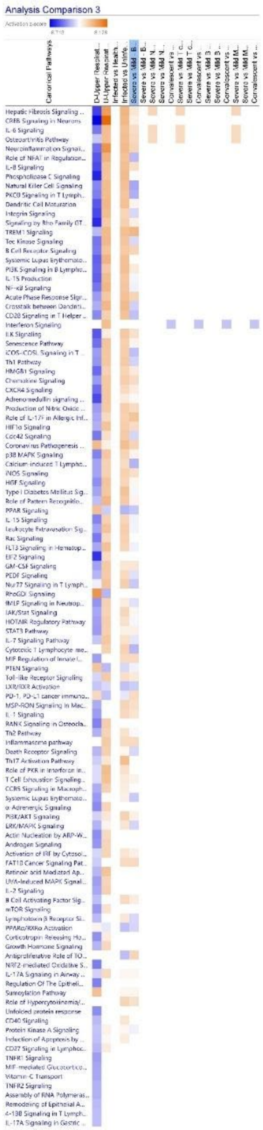


**Supplementary Figure 6. Comparison analysis across the four analyzed datasets reveals canonical pathways consistently significant in COVID-19 patient samples.** Heatmap was generated in IPA. Blue represents inhibition and orange represents activation, based on z-score.

**Supplementary Table 1. Descriptions of genes mentioned in paper.** Genes are referred to by their GeneCards code within the paper, and descriptions were retrieved from the Gene Cards database (https://www.genecards.org/).

| GeneCard Code | Description |
|---|---|
| ARG1 | Arginase1 |
| CACNA2D3 | Calcium Voltage-Gated Channel Auxiliar Subunit Alpha2delta 3 |
| CAMP | Cathelicidin Antimicrobial Peptide |
| CASC | Cancer Susceptibility 8 |
| CCL2 | C-C Motif Chemokine Ligand 2 |
| CCL3L1 | C-C Motif Chemokine Ligand 3 Like 1 |
| CCL6 | C-C Motif Chemokine Ligand 6 |
| CCL7 | C-C Motif Chemokine Ligand 7 |
| CD3E | CD3e Molecule |
| CD3G | CD3g Molecule |
| CD52 | CD52 Molecule |
| CD8A | CD8a Molecule |
| CD8B | CD8b Molecule |
| CES1 | Carboxylesterase 1 |
| CSF2 | Colony Stimulating Factor 2 |
| CTSW | Cathepsin W |
| CXCL10 | C-X-C Motif Chemokine Ligand 10 |
| CXCL8 | C-X-C Motif Chemokine Ligand 8 |
| CXCR3 | C-X-C Motif Chemokine Receptor 3 |
| CXCR6 | C-X-C Motif Chemokine Receptor 6 |
| DHRS9 | Dehydrogenase/Reductase 9 |
| DPM3 | Dolichyl-Phosphate Mannosyltransferase Subunit 3, Regulatory |
| E2F7 | E2F Transcription Factor 7 |

| | |
|---|---|
| eIF2α | Eukaryotic Translation Initiation Factor Alpha |
| eIF2γ | Eukaryotic Translation Initiation Factor Gamma |
| FASLG | Fas Ligand |
| FOSB | FosB Proto-Oncogene, AP-1 Transcription Factor Subunit |
| HLA-DBQ1 | Major Histocompatibility Complex, Class II, DQ Beta 1 |
| HLA-DBQ2 | Major Histocompatibility Complex, Class II, DQ Beta 2 |
| HLA-DPB1 | Major Histocompatibility Complex, Class II, DP Beta 1 |
| HLA-DQA2 | Major Histocompatibility Complex, Class II, DQ Alpha 2 |
| HRI | Heme-Regulated eIF2α kinase |
| IDO1 | Indoleamine 2,3-Dioxygenase 1 |
| IFI27 | Interferon Alpha Inducible Protein 27 |
| IFI44L | Interferon Induced Protein 44 Like |
| IFIT1 | Interferon Induced Protein With Tetratricopeptide Repeats 1 |
| IFITM2 | Interferon Induced Transmembrane Protein 2 |
| IFITM3 | Interferon Induced Transmembrane Protein 3 |
| IFN-gamma | Inteferon Gamma |
| IGHG1 | Immunoglobulin Heavy Constant Gamma 1 (G1m Marker) |
| IGHV1-46 | Immunoglobulin Heavy Variable 1-46 |
| IGHV4-59 | Immunoglobulin Heavy Variable 1-59 |
| IGKC | Immunoglobulin Kappa Constant |
| IKZF2 | IKAROS Family Zinc Finger 2 |
| IL-1A | Interleukin 1 Alpha |
| IL-1B | Interleukin 1 Beta |
| IL-6 | Interleukin 6 |
| IL1R2 | Interleukin Receptor Type 2 |
| ISG15 | ISG15 Ubiquitin Like Modifier |
| JAML | Junction Adhesion Molecule Like |
| LAG3 | Lymphocyte Activating 3 |

| | |
|---|---|
| LINC00877 | Long Intergenic Non-Protein Coding RNA 877 |
| MEFV | MEFV Innate Immunity Regulator, Pyrin |
| MME | Membrane Metalloendopeptidase |
| MRPL53 | Mitochondrial Ribosomal Protein L53 |
| MT3 | Metallothionein 3 |
| mTORC1 | Mechanistic Target of Rapamycin Kinase Complex 1 |
| MX1 | MX Dynamin Like GTPase 1 |
| OAS3 | 2'-5'-Oligoadenylate Synthetase 3 |
| Pax6 | Paired Box 6 |
| PERK | Protein Kinase R (PKR)-like endoplasmic reticulum kinase |
| PKR | Protein Kinase R |
| RB1 | RB Transcriptional Corepressor 1 |
| RELB | RELB Proto-Oncogene, NFkB Subunit |
| ROMO1 | Reactive Oxygen Species Modulator |
| RSAD2 | Radical S-Adenosyl Methionine Domain Containing 2 |
| S100A12 | S100 Calcium Binding Protein A12 |
| SCGB3A1 | Secretoglobin Family 3A Member 1 |
| SLAMF7 | SLAM Family Member 7 |
| SLC25A37 | Solute Carrier Family 25 Member 37 |
| SNCA | Synuclein Alpha |
| SPP1 | Secreted Phosphoprotein 1 |
| TNF | Tumor Necrosis Factor |
| TNFSF12 | TNF Superfamily Member 12 |
| TREM1 | Triggering Receptor Expressed On Myeloid Cells 1 |
| VMO1 | Vitelline Membrane Outer Layer 1 Homolog |
| XCL1 | X-C Motif Chemokine Ligand 1 |
| ZNF683 | Zinc Finger Protein 683 |

**Supplementary Table 2. Classification counts for GSE145926 quantifies immune cell populations in each BALF sample.** All the cells grouped in each immune cell subset per sample are listed. N/A defines the cells not classified in an immune cell subset.

| Sample name | # Cells | Disease State | B cells | Macrophages | NK cells | T cells | N/A |
|---|---|---|---|---|---|---|---|
| Healthy Control_C100 | 8972 | Uninfected | 13 | 604 | 152 | 138 | 1321 |
| Healthy Control_C51 | 11115 | Uninfected | 14 | 2419 | 70 | 917 | 5632 |
| Healthy Control_C52 | 10366 | Uninfected | 21 | 2591 | 16 | 532 | 5163 |
| Mild COVID-19 Patient_C141 | 6249 | Mild COVID-19 Patient | 24 | 434 | 576 | 516 | 500 |
| Mild COVID-19 Patient_C142 | 10269 | Mild COVID-19 Patient | 25 | 666 | 392 | 288 | 884 |
| Mild COVID-19 Patient_C144 | 3716 | Mild COVID-19 Patient | 10 | 84 | 111 | 68 | 373 |
| Severe COVID-19 Patient_C143 | 20857 | Severe COVID-19 Patient | 73 | 9073 | 811 | 202 | 8167 |
| Severe COVID-19 Patient_C145 | 18044 | Severe COVID-19 Patient | 78 | 4467 | 656 | 457 | 10508 |
| Severe COVID-19 Patient_C146 | 4111 | Severe COVID-19 Patient | 55 | 704 | 50 | 56 | 1905 |
| Severe COVID-19 Patient_C148 | 3920 | Severe COVID-19 Patient | 68 | 748 | 61 | 44 | 656 |
| Severe COVID-19 Patient_C149 | 2879 | Severe COVID-19 Patient | 35 | 1143 | 173 | 81 | 725 |
| Severe COVID-19 Patient_C152 | 7732 | Severe COVID-19 Patient | 157 | 801 | 362 | 111 | 1755 |
| All samples | N/A | N/A | 573 | 23734 | 3430 | 3410 | 37589 |

**Supplementary Table 3. Classification counts for GSE154567 quantifies immune cell populations in each blood buffy coat sample.** All the cells grouped in each immune cell subset per sample are listed. N/A defines the cells not classified in an immune cell subset.

| Sample name | # Cells | Disease State | B cells | Macrophages | NK Cells | T Cells | N/A |
|---|---|---|---|---|---|---|---|
| Mild COVID-19 Patient_CM1_CM2 | 7523 | Mild COVID-19 Patient | 183 | 1381 | 1913 | 1165 | 2881 |
| Mild COVID-19 Patient_CM3_CM4 | 9341 | Mild COVID-19 Patient | 123 | 1037 | 3546 | 1269 | 3366 |
| Mild COVID-19 Patient_CM5 | 7828 | Mild COVID-19 Patient | 479 | 868 | 2295 | 1996 | 2190 |
| Recovered COVID19 Patient_CR13_CR14 | 12442 | Recovered COVID-19 Patient | 455 | 1929 | 2257 | 1678 | 6123 |
| Recovered COVID19 Patient_CR15_CR16 | 10583 | Recovered COVID-19 Patient | 591 | 753 | 2670 | 398 | 6171 |
| Recovered COVID19 Patient_CR17_CR18 | 11485 | Recovered COVID-19 Patient | 437 | 1181 | 1883 | 2562 | 5422 |
| Severe COVID-19 Patient_CS11_C12 | 10440 | Severe COVID-19 Patient | 524 | 551 | 1955 | 3307 | 4103 |
| Severe COVID-19 Patient_CS7_CS8 | 3976 | Severe COVID-19 Patient | 186 | 191 | 801 | 1187 | 1611 |
| Severe COVID-19 Patient_CS9_C10 | 11446 | Severe COVID-19 Patient | 97 | 75 | 857 | 768 | 9649 |
| All samples | N/A | N/A | 3075 | 7966 | 18177 | 14330 | 41516 |

**Supplementary Table 4. Upstream regulators of significant dataset molecules present potential host therapeutic targets for COVID-19.** Regulators were identified from the "Regulator Effects" function in IPA, applied to all pathway analyses.

| Regulators | Target Molecules in Dataset | Diseases & Functions |
|---|---|---|
| 3M-002 | CCL5,CD69,CD86,CLEC4E,CXCL10,CXCL3,TLR8,TNFSF10 | Stimulation of cells |
| ACKR2 | CCL22,CCL5,CCR2,CXCL10,DDX58,STAT1 | Cell movement of lymphoid cells |
| ACKR2 | CCL22,CCL5,CCR2,CXCL10,DDX58,STAT1 | Lymphocyte migration |
| ACKR2 | CCL22,CCL5,CCR2,CXCL10,DDX58,STAT1 | Migration of mononuclear leukocytes |
| ACTN4 | CCL2,CXCL8,IL1B,SERPINE1 | Binding of tumor cell lines |
| ACTN4 | CCL2,CXCL8,IL1B,SERPINE1 | Cell movement of breast cancer cell lines |
| ACTN4 | CCL2,CXCL8,IL1B,SERPINE1 | Migration of tumor cell lines |
| ADA | ADORA3,DCK,MMP12,SPP1 | Quantity of leukocytes |
| ADORA3 | CCL2,CCL3,CCL4,CCL5,CCL7,SPP1 | Cell movement of antigen presenting cells |

| | | |
|---|---|---|
| ADORA3 | CCL2,CCL3,CCL4,CCL5,CCL7,CCL8,CXCL8,SPP1,VEGFA | Cell movement of blood cells |
| ADORA3 | CCL3,CCL4,CCL5,CCL7,SPP1 | Cell movement of dendritic cells |
| ADORA3 | CCL2,CCL3,CCL4,CCL5,CCL7,CCL8,CXCL8,SPP1 | Cell movement of phagocytes |
| ADORA3 | CCL2,CCL3,CCL4,CCL5,CCL7,CCL8,CXCL8,SPP1,VEGFA | Chemotaxis |
| ADORA3 | CCL2,CCL3,CCL4,CCL5,CCL7 | Chemotaxis of antigen presenting cells |
| ADORA3 | CCL3,CCL4,CCL5,CCL7 | Chemotaxis of dendritic cells |
| ADORA3 | CCL2,CCL3,CCL4,CCL5,CCL7,CCL8,CXCL8,SPP1 | Chemotaxis of leukocytes |
| ADORA3 | CCL2,CCL4,CCL5,CCL8,CXCL3,IL10,SPP1 | Chemotaxis of phagocytes |
| ADORA3 | CCL2,CCL3,CCL4,CCL5,CCL7,CCL8,CXCL8,SPP1 | Chemotaxis of phagocytes |
| ADORA3 | CCL2,CCL4,CXCL3,IL10 | Degranulation of cells |
| ADORA3 | CCL2,CCL4,CCL5,CCL8,CXCL3,IL10,SPP1 | Homing of leukocytes |
| ADORA3 | CCL2,CCL3,CCL4,CCL5,CCL7,CCL8,CXCL8,SPP1,VEGFA | Leukocyte migration |
| AGT | ACE,ADIPOQ,CD86,FCGR1A,MAS1,PPARG,PTGS1 | End stage renal disease |
| aldesleukin | CD86,FCGR3A/FCGR3B,JAK2,STAT1 | Activation of cells |

| | | |
|---|---|---|
| aldosterone | CYBB,EDN1,FLT1,OLR1,PPARG,SPP1 | Cell death of kidney cells |
| alefacept | CCL5,CD3D,CD69,GNLY,IL2RA,IL7R,STAT1 | Quantity of T lymphocytes |
| Alpha catenin | ADAMTS1,BCL3,BIRC3,CXCL10,CXCL12,IGF1,IL2RG,LYN,MMP12, MMP13,TNFAIP6,WIPF1,ZEB2 | Quantity of cells |
| anakinra | CCL2,CCL4,CXCL12,EDN1 | Recruitment of myeloid cells |
| APOE | C1QA,SPP1,STAT4 | Increased Levels of ALT |
| ATM | CLU,CXCL8,DUSP1,IL6,RRM2,TNFAIP3 | Cell survival |
| ATM | CLU,CXCL8,DUSP1,IL6,RRM2,TNFAIP3 | Cell viability |
| BCR (complex) | CCL4,CD69,IL10,JAK2,REL,SELL | Homing of leukocytes |
| BIRC2 | CCL2,CXCL1,CXCL8,IL27,IL6 | Cellular homeostasis |
| bisindolylmaleimide I | ACE,FASLG,IGF1,TNFSF10 | Activation of phagocytes |
| BRD4 | BCL3,CCR1,IL2RA,IL6R,IL7R,ITGA1,KCNA3,PDGFA | Migration of cells |
| bromodeoxyuridine | CXCL10,IFI16,IFI44,IFI6,IFIH1,IFIT1,IFIT2,IFIT3,IFITM1,INHBA,IRF4,MX1,PML,SP110,STAT1,TNFSF10 | Glioma |
| BTK | CCL4,CXCL9,SPN | Adhesion of lymphocytes |
| C1q | CCL2,CD80,IL10 | Cell viability |
| C3 | C3AR1,CCL2,CCL4,CCL5,CXCL10,CXCL3,IL10,SELL | Chemotaxis of phagocytes |
| C5 | CXCL8,ICAM1,IL10,IL1B,IL6,MMP9,SERPINE1 | Cell movement of blood cells |

| | | |
|---|---|---|
| C5 | CXCL8,ICAM1,IL10,IL1B,IL6,MMP9,SERPINE1 | Leukocyte migration |
| C5 | CCL2,CCL5,CD86,CXCL10,CXCL3,FASLG,IL10 | Stimulation of leukocytes |
| C5AR1 | CD80,CD86,FCER1G,IGF1,IL10 | Quantity of lymphoid tissue |
| CAMP | CCL2,CCL3,CCL5,CCL7,CXCL8,IL10,IL1A,IL1B,IL6,TLR2 | Activation of blood cells |
| CAMP | CCL2,CCL3,CCL5,CCL7,CXCL8,IL10,IL1A,IL1B,IL6,TLR2 | Activation of leukocytes |
| CAMP | CCL2,CCL3,CCL5,CCL7,CXCL8,IL1A,IL1B,TLR2 | Activation of myeloid cells |
| CAMP | CCL2,CCL3,CCL5,CCL7,CXCL8,IL10,IL1A,IL1B,IL6,TLR2 | Activation of phagocytes |
| CAMP | CCL2,CCL3,CCL4,CCL5,CCL7,CXCL8,IL1B,IL6,TLR2 | Cell movement of carcinoma cell lines |
| CAMP | CCL2,CCL3,CCL5,CCL7,CXCL10,CXCL8,IL1B | Cell movement of eosinophils |
| CAMP | CCL2,CCL3,CCL4,CCL5,CCL7,CXCL1,CXCL8,FPR2,IL10,IL1B,IL6,TLR2 | Cellular homeostasis |
| CAMP | CCL2,CCL4,CCL5,CXCL10,CXCL13,CXCL3,CXCR2,IL10,TLR2,TLR4 | Chemotaxis |
| CAMP | CXCL1,CXCL10,CXCL8 | Response of myeloid leukocytes |
| CARD9 | CCR1,CCR5,CLEC7A,CXCL10,CXCL3,IL10,TLR2 | Organismal death |
| cardiotoxin | CASP1,CCL2,CCR2,CCR5,CSF1R,IL2RG,MMP12,SPP1 | Quantity of macrophages |

| | | |
|---|---|---|
| CASP8 | CCR5,CD69,CXCL10,IL2RA,TSLP | Quantity of T lymphocytes |
| CCL11 | AIF1,CCL18,CCL2,CCL4,CCL8,CCR2,CXCL11,CXCL12,CXCL3,CXCL9, FASLG,ITGA1,PDGFA,TNFSF14 | Cell movement |
| CCL11 | AIF1,CCL18,CCL2,CCL4,CCL8,CCR2,CXCL11,CXCL12,CXCL3,CXCL9, FASLG,ITGA1,TNFSF14 | Cell movement of blood cells |
| CCL11 | CCL18,CCL2,CCL4,CCL8,CCR2,CXCL11,CXCL12,CXCL9,TNFSF14 | Homing of T lymphocytes |
| CCL11 | AIF1,CCL18,CCL2,CCL4,CCL8,CCR2,CXCL11,CXCL12,CXCL3,CXCL9, FASLG,ITGA1,TNFSF14 | Inflammatory response |
| CCL11 | AIF1,CCL18,CCL2,CCL4,CCL8,CCR2,CXCL11,CXCL12,CXCL3,CXCL9, FASLG,ITGA1,TNFSF14 | Leukocyte migration |
| CCL11 | AIF1,CCL18,CCL2,CCL4,CCL8,CCR2,CXCL11,CXCL12,CXCL3,CXCL9, FASLG,ITGA1,PDGFA,TNFSF14 | Migration of cells |
| CCL5 | CCL2,CCL4,CCR1,IL6 | Binding of lymphocytes |
| CCL5 | CCL2,CCL3,CCL4,CCR1,IL6 | Binding of mononuclear leukocytes |
| CCL5 | CCL4,CCR1 | Binding of T lymphocytes |
| CCL5 | CCL3,CCL4 | Cell movement of monocyte-derived dendritic cells |
| CCL5 | CCL2,CCL3,CCL4,CCR1,CXCL8 | Cell movement of peripheral blood monocytes |

| | | |
|---|---|---|
| CCL5 | CCL2,CCL3,CCL4,CXCL8,DUSP6,IL1B,IL6,MMP9,NAMPT | Cell movement of tumor cell lines |
| CCL5 | CCL2,CCL3,CCL4,CCR1,CXCL8 | Chemotaxis of PBMCs |
| CCL5 | CCL2,CCL3,CCL4,CCR1,CXCL8 | Chemotaxis of peripheral blood monocytes |
| CCL5 | CCL2,STAT1 | Cytotoxicity of lymphocytes |
| CCL5 | CCL2,CCL3,CCL4,CCR1,CXCL8 | Flux of Ca2 |
| CCL5 | CCL2,CCL3,CCL4,CCR1,CXCL8 | Flux of ion |
| CCL5 | CCL2,CCL4,CCR1,IL6 | Interaction of lymphocytes |
| CCL5 | CCL2,CCL3,CCL4,CCR1,IL6 | Interaction of mononuclear leukocytes |
| CCL5 | CCL4,CCR1 | Interaction of T lymphocytes |
| CCL5 | CCL2,CCL3,CCL4,CCR1,CXCL8 | Ion homeostasis of cells |
| CCL5 | CCL2,CCL3,CXCL8,IL1B | Recruitment of leukocytes |
| CCR2 | CD86,CXCL12,IGF1,IL10 | Quantity of hematopoietic progenitor cells |

| | | |
|---|---|---|
| CCR5 | CCL4,CCL5,CXCL10,CXCL12,CXCL3,FASLG,IL10 | Chemotaxis of myeloid cells |
| CCR5 | CCL4,CCL5,CD40LG,CXCL10,CXCL12,FASLG,IL10 | Interaction of T lymphocytes |
| CD2 | CD40LG,CD80,CD86,FASLG,HLA-DMA,IL10,PTPRC,STAT1,STAT4 | Cellular homeostasis |
| CD2 | CD40LG,CD80,CD86,FASLG,HLA-DMA,IL10,PTPRC,STAT1,STAT4 | Development of mononuclear leukocytes |
| CD2 | CD40LG,CD80,CD86,FASLG,HLA-DMA,IL10,PTPRC,STAT1,STAT4 | Hematopoiesis of mononuclear leukocytes |
| CD28 | CCL3,CCL4,CXCL8,CXCR3,FASLG,FYN,IL10,IL1B,PTGS2,XCL1 | Chemotaxis |
| CD28 | GATA3,IL10,IL1B,TBX21 | Differentiation of naive lymphocytes |
| CD28 | GATA3,IL10,IL1B,TBX21 | Differentiation of naive T lymphocytes |
| CD28 | GATA3,IL10,IL1B,TBX21 | Differentiation of T lymphocytes |
| CD28 | GATA3,IL10,IL1B,NFKBIA | Quantity of lymphatic system cells |
| CD28 | GATA3,IL10,NFKBIA | Quantity of lymphocytes |
| CD28 | GATA3,IL10,IL1B,NFKBIA | Quantity of lymphoid cells |

| | | |
|---|---|---|
| CD28 | CXCL8,FYN,IL10,IL1B | Stimulation of leukocytes |
| CD3 | CCL3,CCL4,CD28,CXCL8,CXCR3,IL10,XCL1 | Migration of lymphatic system cells |
| CD3 | CCL3,CCL4,CD28,CXCL8,CXCR3,IL10,XCL1 | T cell migration |
| CD3 group | DGKA,FASLG,FOXP3,HAVCR2,IL10,IL2RA | Cellular homeostasis |
| CD3 group | DGKA,FASLG,FOXP3,HAVCR2,IL10,IL2RA | Leukopoiesis |
| CD40 | CCL2,CCL5,ICAM1,IL10,IL1A,IL6,IL6R | Binding of mononuclear leukocytes |
| CD40LG | CCL2,CCL3,CCL4,CD40,CXCL10,CXCL8,ICAM1,IL10,IL1A,IL1B,IL21R, IL6,VEGFA | Leukocyte migration |
| CD40LG | BCL6,CCL2,CD40,CXCL8,IL10,IL1A,IL1B,IL21R,IL6,TNFSF13B,VEGFA | Leukopoiesis |
| CD40LG | CASP1,CCL2,CCL5,CD69,CD80,CD86,CXCL10,FASLG,IL10,MED1,TNFSF10,TNFSF13 B | Stimulation of leukocytes |
| CD80 | CD40LG,IL10 | Priming of T lymphocytes |
| cigarette smoke | CCL2,CCL4,CXCL10,CXCL3,IL10,TLR4,TSLP | Chemotaxis of mononuclear leukocytes |
| CLEC4E | CCL4,CLEC5A | Recruitment of macrophages |
| CLEC4E | CXCL3,IL10 | Stimulation of leukocytes |
| COL18A1 | CCL2,ETS1,IL6,JUN,NRP1,PLAU,PTGS2,VEGFA | Invasion of tumor cell lines |

| | | |
|---|---|---|
| COL18A1 | ETS1,IL6,PLAU,PTGS2,VEGFA | Synthesis of reactive oxygen species |
| corticosteroid | ACE,CD69,GZMA,IL10,PPARG,PTGS1 | Inflammation of body cavity |
| corticosteroid | ACE,CD69,GZMA,IL10,PPARG,PTGS1 | Inflammation of organ |
| CpG ODN 1826 | CCL2,CCL5,CD274,CD69,CD80,CD86,CXCL10,CXCL3,CYBB,IL10,REL, TLR4 | Leukopoiesis |
| CRNDE | DUSP6,NR4A1,VEGFA | Colony formation |
| CRNDE | DUSP6,NR4A1,VEGFA | Migration of tumor cell lines |
| CRNDE,IL6 | CLU,DUSP6,IL6R,NFKBIA,NR4A1,PLAU,PTGS2,VEGFA | Colony formation |
| CRP | IL1B,IL6R,S100A12,SERPINE1 | Inflammatory response |
| CSF2 | CCL2,CD40,CEACAM1,CXCL8,DUSP6,IL1B,IL6,ITGA4,MMP9,PTGS2, TLR2 | Migration of tumor cell lines |
| CTLA4 | CD69,CD80,CD86,CXCL10,FASLG,IL10 | Stimulation of leukocytes |
| CXCL12 | BCL3,CCL5,CD40LG,CD69,IL2RA,LCP2 | Differentiation of T lymphocytes |
| CXCL2 | CCL4,CCL5,CXCL10,CXCL3,CXCR2 | Activation of cells |
| CXCL2 | CCL4,CCL5,CXCL10,CXCL3,CXCR2 | Adhesion of blood cells |
| CXCL2 | CCL4,CCL5,CXCL10,CXCL3,CXCR2 | Binding of leukocytes |

| | | |
|---|---|---|
| CXCL2 | CCL4,CCL5,CXCL10,CXCL3,CXCR2 | Cell movement of neutrophils |
| CXCL2 | CCL4,CCL5,CXCL10,CXCL3,CXCR2 | Cellular infiltration by leukocytes |
| CXCL2 | CCL4,CCL5,CXCL10,CXCL3,CXCR2 | Cellular infiltration by myeloid cells |
| CXCL2 | CCL4,CCL5,CXCL10,CXCL3,CXCR2 | Chemotaxis of myeloid cells |
| CXCL2 | CCL4,CCL5,CXCL10,CXCL3,CXCR2 | Chemotaxis of phagocytes |
| CXCL2 | CCL4,CCL5,CXCL10,CXCL3,CXCR2 | Migration of phagocytes |
| CXCL2 | CCL4,CCL5,CXCL10,CXCL3,CXCR2 | Recruitment of myeloid cells |
| CXCL2 | CCL4,CCL5,CXCL10,CXCL3,CXCR2 | Recruitment of phagocytes |
| CXCL8 | FASLG,IL1B,ITGA5,MMP9,PTGS2 | Cell movement of tumor cell lines |
| CXCL8 | ABCB1,FASLG,IL1B,ITGA5,MMP9,PTGS2 | Hematopoieti c neoplasm |
| CXCL8 | FASLG,IL1B,ITGA5,MMP9,PTGS2 | Invasion of tumor cell lines |
| CXCL8 | ABCB1,FASLG,IL1B,ITGA5,MMP9,PTGS2 | Lymphohema topoietic neoplasia |

| | | |
|---|---|---|
| CXCL8 | FASLG,IL1B,ITGA5,MMP9,PTGS2 | Migration of cells |
| CXCR4 | CCL2,CD69,CXCL10,CXCL12,CXCR2,IL2RA,TNFSF10,ZEB1 | Hematopoiesis of mononuclear leukocytes |
| CYBB | CCL5,CXCL10,CXCL12,CXCL3,PPARG | Activation of cells |
| cyclic AMP | CD80,CD86,ENTPD1,IGF1,IL10,IL2RA,STAT1 | Differentiation of T lymphocytes |
| cyclosporin A | CCL2,CCL4,CCL5,CXCL9,FLT1,IL10,LYN | Chemotaxis of monocytes |
| cytokine | CCL2,CCL22,CCL5,CD40LG,CD80,CD86,CXCL3,EDN1,TLR2,TLR4 | Activation of myeloid cells |
| cytokine | CCL2,CCL22,CCL5,CD209,CD40LG,CD69,CD80,CD86,CXCL3,LIF,LIPG,NOS1,TLR2,TLR4 | Adhesion of blood cells |
| cytokine | CCL2,CCL22,CCL5,CD209,CD40LG,CD69,CD80,CD86,CXCL3,LIF,LIPG,NOS1,TLR2,TLR4 | Adhesion of immune cells |
| cytokine | CCL2,CCL22,CCL5,CD209,CD40LG,CD69,CD80,CD86,CXCL3,LIF,LIPG,NOS1,TLR2,TLR4 | Binding of blood cells |
| cytokine | CCL2,CCL22,CCL5,CD209,CD40LG,CD69,CD80,CD86,CXCL3,LIF,LIPG,NOS1,TLR2,TLR4 | Binding of leukocytes |
| cytokine | CCL2,CCL22,CD40LG,CXCL3,TLR2,TLR4 | Response of myeloid cells |
| cytokine | CCL2,CCL22,CCL5,CD40LG,CD86,CXCL3,TLR2,TLR4 | Response of phagocytes |
| cytokine | CCL2,CCL22,CCL5,CD209,CD40LG,CD69,CD80,CD86,CXCL3,EDN1, LIF,PDCD1LG2,TLR2,TLR4 | Stimulation of cells |
| daporinad | BRIP1,CCL2,LIF,LMNB1 | Cell viability |
| DDX58 | CCL2,CCL5,IFIH1,IFNL1,IFNL2,IRF8,STAT1,TLR3,TNFSF10 | Maturation of antigen |

| | | |
|---|---|---|
| | | presenting cells |
| DDX58 | CCL2,CCL5,IFIH1,IFNL1,IFNL2,IRF8,STAT1,TLR3,TNFSF10 | Maturation of leukocytes |
| DEPTOR | CCL5,CXCL10,CXCL11,CXCL9 | Adhesion of blood cells |
| DEPTOR | CCL5,CXCL10,CXCL11,CXCL9 | Binding of lymphatic system cells |
| DEPTOR | CCL5,CXCL10,CXCL11,CXCL9 | Binding of T lymphocytes |
| DEPTOR | CCL5,CXCL10,CXCL11,CXCL9 | Cell movement of lymphoid cells |
| DEPTOR | CCL5,CXCL10,CXCL11,CXCL9 | Interaction of T lymphocytes |
| DEPTOR | CCL5,CXCL10,CXCL11,CXCL9 | Migration of mononuclear leukocytes |
| D-glucose | EDN1,IGF1,ITGA1,TLR2,TLR4 | Proliferation of mesangial cells |
| DOCK8 | CXCL10,SLFN12L,TLR3 | Quantity of antigen presenting cells |
| EDN1 | CCL2,ITPR2,MAS1,SELL | Mobilization of Ca2 |
| EGF | CXCL8,MMP9,PTGS2,SERPINE1,VEGFA | Cell movement of endothelial cells |

| | | |
|---|---|---|
| epigallocatechin-gallate | CCL2,CCL22,CCL5,CXCL10,FASLG,IL10,TLR4 | Interaction of lymphocytes |
| ERBB2 | CCL2,CXCL8,IL6,ITGA5,JUN,MUC1,PIK3CD,PLAU,PTGS2,RRM2, SERPINE1,VEGFA | Invasion of cells |
| ERK | CCL2,CXCL10,CXCL2,CXCL8,ETS1,FOSL1,IL6,JUN,MMP9,MUC1, PTGS2,SERPINE1 | Migration of tumor cell lines |
| ERK1/2 | CCL2,CCL4,CCL5,CXCL8,CXCR4,ICAM1,IL1A,IL1B,IL6,VEGFA | Adhesion of immune cells |
| ERK1/2 | CCL2,CCL4,CXCR4,IL6 | Adhesion of lymphocytes |
| ERK1/2 | CCL2,CCL4,CXCR4,ICAM1,IL1A,IL6 | Adhesion of mononuclear leukocytes |
| ERK1/2 | CCL2,CCL3,CCL4,CXCL8,CXCR4,ICAM1,IL6,JUN,VEGFA | Adhesion of tumor cell lines |
| ERK1/2 | C3,CCL2,CCL3,CCL4,CCL5,CD40,CXCL8,CXCR4,ICAM1,IL1A,IL1B,IL6,VEGFA | Binding of blood cells |
| ERK1/2 | CCL2,CCL3,CCL4,CCL5,CD40,CXCL8,CXCR4,ICAM1,IL1A,IL1B,IL6, VEGFA | Binding of leukocytes |
| ERK1/2 | CCL2,CCL3,CCL4,CXCL8,CXCR4,ICAM1,IL1B,IL6,JUN,VEGFA | Binding of tumor cell lines |
| ERK1/2 | C3,CCL2,CCL3,CCL4,CCL5,CCL8,CCR7,CD40,CXCL8,CXCR4,FOSB, ICAM1,IL1A,IL1B,IL6,JUN,PTGS2,VEGFA | Cell movement |
| ERK1/2 | CCL2,CCL3,CCL4,CCL5,CCL8,CCR7,CXCL8,CXCR4,ICAM1,IL6 | Cell movement of lymphatic system cells |

| | | |
|---|---|---|
| ERK1/2 | CCL2,CCL3,CCL4,CCL5,CCL8,CCR7,CXCL8,CXCR4,ICAM1 | Cell movement of lymphocytes |
| ERK1/2 | CCL2,CCL3,CCL4,CCL5,CCL8,CCR7,CXCL8,CXCR4,ICAM1,IL1B | Cell movement of mononuclear leukocytes |
| ERK1/2 | CCL2,CCL3,CCL4,CCL5,CCL8,CCR7,CXCL8,CXCR4,ICAM1 | Cell movement of PBMCs |
| ERK1/2 | C3,CCL2,CCL3,CCL5,CCR7,CD40,CXCL8,CXCR4,ICAM1,IL1A,IL1B,IL6, JUN,PTGS2,VEGFA | Cell viability |
| ERK1/2 | C3,CCL3,CCR7,CD40,CXCL8,CXCR4,ICAM1,IL1B,IL6 | Cell viability of blood cells |
| ERK1/2 | C3,CCR7,CD40,CXCL8,CXCR4,ICAM1,IL1B,IL6 | Cell viability of leukocytes |
| ERK1/2 | CCL2,CCL3,CCL4,CCL5,CCR7,CXCL8,CXCR4,ICAM1 | Lymphocyte migration |
| ERK1/2 | CCL2,CCL3,CCL4,CCL5,CCR7,CXCL8,CXCR4,ICAM1,IL6 | Migration of lymphatic system cells |
| ERK1/2 | CCL2,CCL3,CCL4,CCL5,CCL8,CCR7,CXCL8,CXCR4,ICAM1 | Migration of mononuclear leukocytes |
| ERK1/2 | CCL2,CCL5,CCR7,CD40,CXCL8,CXCR4,IL1A,IL1B,IL6,JUN,PTGS2, VEGFA | Migration of tumor cell lines |
| ERK1/2 | CCL2,CCL22,CCL5,CCND2,CD80,CD86,CXCL10,CXCL3,EDN1,IL10, MED1,SPP1,TSLP | Stimulation of cells |
| ERK1/2 | CCL2,CCL3,CCL4,CCL5,CCR7,CXCL8,CXCR4,ICAM1 | T cell migration |
| estrogen | BCL3,CXCL12,IGF1,IL10,LIF,SPP1,STAT1 | Quantity of stem cells |

| | | |
|---|---|---|
| **ethanol** | **CCL2,CCL5,CD40LG,CXCL3,EDN1,FASLG,IL10,TLR1,TLR2,TLR4,TLR8** | **Stimulation of cells** |
| **F2** | **CXCL8,ICAM1,IL6,MMP9,OSM,PTGS2** | **Migration of cells** |
| **F2** | **CCL2,CCL8,CXCL10,CXCL13,CXCL3,EDN1** | **Mobilization of Ca2** |
| **F7** | **CXCL2,CXCL8,FOSL1,IL1B,VEGFA** | **Invasion of cells** |
| **FAS** | **CCL2,CCL5,CD80,CD86,CXCL10,CXCL11,CXCL3,FASLG,IL10,LILRB2, LILRB4,TLR2,TNFSF10** | **Stimulation of leukocytes** |
| **FASLG** | **BIRC2,CCL2,CXCL10,IL10,STAT4** | **Quantity of leukocytes** |
| **FAT1** | **IL1B,IL6,JUN,PLAU,PTGS2** | **Cell movement of tumor cell lines** |
| **FAT1** | **IL1B,IL6,JUN,PLAU,PTGS2** | **Invasion of tumor cell lines** |
| **FAT1** | **IL1B,IL6,JUN,PLAU,PTGS2** | **Migration of tumor cell lines** |
| **Fcer1** | **CCL18,CCL2,CCL4,CCL5,CXCL3,IL10,INHBA,LIF** | **Chemotaxis** |
| **FGF2** | **CXCR4,MMP9,PLAU,SERPINE1,VEGFA** | **Cell movement of breast cancer cell lines** |
| **FGF2** | **CXCR4,MMP9,PLAU,SERPINE1,VEGFA** | **Migration of tumor cell lines** |
| **FLT3LG** | **CD80,CD86,FOXP3,IL10,IL7R,PTPRC** | **Quantity of lymphoid tissue** |

| | | |
|---|---|---|
| FOXM1 | ACE,ACE2,BRCA2,BRIP1,CCND2,CDK2,FLT1,IGF1,MCM8,NBN, NEDD4,PDGFA,TGFBR2,ZEB1,ZEB2 | Brain tumor |
| FOXM1 | ACE,BRCA2,BRIP1,CCND2,CDK2,CXCL12,FLT1,IGF1,NBN,PDGFA, TGFBR2,ZEB1,ZEB2 | Cell viability |
| FOXM1 | ACE,ACE2,BRCA2,BRIP1,CCND2,CDK2,FLT1,IGF1,MCM8,NBN, NEDD4,PDGFA,TGFBR2,ZEB1,ZEB2 | Intracranial lesion |
| GDF2 | CRYAB,CXCL12,EDN1,IGF1,MDFI,SPP1,TLR4 | Quantity of cells |
| GFI1 | BBC3,BCL3,CASP1,ENTPD1,GFI1B,GIMAP4,GPR183,IL10,IL6R,IL7R,MAFB,REL,ST AT1,STAT4 | T cell development |
| GFI1 | BBC3,BCL3,CASP1,CASP8,ENTPD1,GFI1B,GIMAP4,GPR183,IL10,IL6R,IL7R,MAFB,R EL,STAT1,STAT4 | T cell homeostasis |
| GLI1 | CDH1,EGR3,MMP9,SPP1,VEGFA | Angiogenesis |
| GNAI3 | CCL2,CCL4,CCL5,CCL8,CXCL3,SPP1 | Cell movement of lymphoid cells |
| GNAI3 | CCL2,CCL4,CCL5,CCL8,CXCL3,SPP1 | Cell movement of neutrophils |
| GNAI3 | CCL2,CCL4,CCL5,CCL8,CXCL3,SPP1 | Chemotaxis of myeloid cells |
| GNAI3 | CCL2,CCL4,CCL5,CCL8,CXCL3,SPP1 | Chemotaxis of phagocytes |
| GNAI3 | CCL2,CCL4,CCL5,CCL8,CXCL3,SPP1 | Lymphocyte migration |
| GNAI3 | CCL2,CCL4,CCL5,CCL8,CXCL3,SPP1 | Migration of mononuclear leukocytes |
| GNAI3 | CCL2,CCL4,CCL5,CCL8,CXCL3,SPP1 | Migration of phagocytes |
| GPR39 | CLU,LIPE,PNPLA2,RGS16 | Necrosis |

| | | |
|---|---|---|
| Hbb-b2 | CCR2,CCR5,CLEC7A,CXCL10,CXCR2,CYBB,IL10RA,PRLR,TLR2 | Migration of cells |
| Hdac | BCL6,CXCL8,IL1B,IL6,JUN,NR4A1,SPP1 | Cell viability |
| Hdac | BCL6,CXCL8,IL6,JUN,NR4A1,SPP1 | Colony formation |
| Hdac | CXCL8,IL1B,IL6,JUN,NR4A1,SPP1 | Invasion of cells |
| HDL | ADIPOQ,CCL2,CCL5,CCR2,CD40LG,CD80,CD86,CX3CR1 | Activation of myeloid cells |
| HGF | CDK2,IL10,INHBA,LGMN,MDM2 | Increased Levels of Red Blood Cells |
| HMGB1 | CCL2,CXCL8,ICAM1,IL6 | Adhesion of tumor cell lines |
| HMGB1 | CCL2,CCL4,CXCL3,IL10,TLR2,TLR4 | Chemotaxis |
| HMGB1 | CCL2,CXCL8,ICAM1,IL6 | Vasculogenesi s |

| | | |
|---|---|---|
| HTT | ABCC10,ADAM23,AGRN,AGT,AKT1,AKT2,ALDOA,ANTKMT,AP1S1, ARHGEF7,ATP1A1,ATP5PO,B4GAT1,BAIAP2,BASP1,BTG3,CAMK4, CBR1,CD9,CDH11,CDKN1A,CEBPA,CEBPB,CHKA,CLDN6,COL16A1, COL6A1,CORO2B,CSN3,CSNK1E,CTNNB1,CTSD,CTTN,CYC1,DBP, DDIT3,DDX50,DHCR7,DLL1,DLST,DNAI1,DRD2,ECI1,ECRG4,EEF1A1, EEF2,ENO2,FASN,FDFT1,FDPS,FGFR1,FHL1,FOLR1,FOS,GAP43, GNAL,GPC1,GPX3,GRK2,GSN,GSS,HMGA1,HRAS,HSPA5,HSPD1, HTRA2,IGFBP5,IL11RA,JUN,JUNB,JUP,KAT2A,KLF4,KLF5,LDHB,LTBP2,LTF,MAST3,MCAM,MDH2,MTCO3,NAB2,NDUFA11,NDUFA3,NDUFS3,NDUFS4,NDUFS7,NEFL,NFIC,NGEF,NGFR,NR4A1,NTRK2,NTS, OPRL1,PABPC4,PC,PCNA,PDLIM4,PDXK,PER1,PITPNM1,PLOD3, POLR2A,POR,PPP1R1A,PPP1R1B,PPP1R7,PPP5C,PRKACA,PROM1, RAP1GAP,RARA,RELA,RGS14,RGS16,RRS1,SART1,SATB1,SCAMP5, SCN4B,SDHA,SEPHS1,SERPINH1,SLC2A1,SOD1,SORT1,SQSTM1, TAGLN,TAGLN3,TCF3,THRA,TIMP3,TNNI2,TNNT3,TPI1,TRAP1, UCHL1,UCP1,VAPA,WNT3A | Development of malignant tumor |
| ICAM1 | CCL4,CCL5,CD69,CD80,CD86,CXCL3,IL2RA | Cellular homeostasis |
| ICAM1 | CCL5,CD69,CD80,CD86,CXCL3,IL2RA,ZEB2 | Leukopoiesis |
| ICAM1 | CCL4,CCL5,CD69,CD80,CD86,IL2RA,ZEB2 | Quantity of T lymphocytes |
| IFIH1 | CCL2,CCL5 | Maturation of dendritic cells |
| IFN alpha/beta | CCL2,CCL5,CCR2,CD69,CD80,CD86,CXCL10,IL10,IL20RB,IRF8,SERPINB9,STAT1,TLR1,TLR2,TLR3,TLR7,TNFSF10,TNFSF13B | Activation of lymphocytes |

| | | |
|---|---|---|
| IFN alpha/beta | CCL2,CCR2,CXCL10,IFI16,IL10,IRF8,STAT1,TLR1,TLR2,TLR7,TNFSF10 | Differentiation of phagocytes |
| IFN alpha/beta | CCL2,CCL5,CD69,CD80,CD86,CXCL10,IL10,TLR1,TLR2,TLR3,TNFSF10,TNFSF13B | Stimulation of cells |
| IFN Beta | CCL3,HLA-G,IDO1,IL27,IL6 | Cellular homeostasis |
| IFNA1/IFNA13 | CD80,CD86,IFIH1,STAT1,TLR3 | Maturation of dendritic cells |
| IFNA10 | CCL8,CXCL10,CXCL9,STAT4,ZBP1 | Activation of cells |
| IFNA14 | CCL8,CXCL10,CXCL9,STAT4,ZBP1 | Activation of cells |
| IFNA16 | CCL8,CXCL10,CXCL9,STAT4,ZBP1 | Activation of cells |
| IFNA2 | IL10,LILRB2,LILRB4 | Differentiation of regulatory T lymphocytes |
| IFNA21 | CCL8,CXCL10,CXCL9,STAT4,ZBP1 | Activation of cells |
| IFNA4 | CXCL10,IFIH1,OASL,RSAD2,ZBP1 | Replication of Herpesviridae |
| IFNA5 | CCL8,CXCL10,CXCL9,STAT4,ZBP1 | Activation of cells |
| IFNA6 | CCL8,CXCL10,CXCL9,STAT4,ZBP1 | Activation of cells |
| IFNA7 | CCL8,CXCL10,CXCL9,STAT4,ZBP1 | Activation of cells |
| IFNA8 | CCL8,CXCL10,CXCL9,STAT4,ZBP1 | Activation of cells |

| | | |
|---|---|---|
| **IFNB1** | **CCL2,IL10,STAT1,STAT4,TNFSF10** | **Cytotoxicity of natural killer cells** |
| **IFNG** | **AIM2,BCL3,BIRC2,BIRC3,CCL22,CCL4,CCL5,CCL8,CD40LG,CD86, CLEC4E,CXCL10,CXCL12,DDX58,DDX60,DTX3L,DUOX2,EIF2AK2, FCER1G,FCGR1A,IFI16,IFI44,IFI44L,IFI6,IFIH1,IFIT1,IFIT2,IFIT3,IFIT5,IFITM1,IFITM2,IFITM3,IL10,IRF8,ISG15,MICA,MMP12,MSR1,MX1, MX2,OAS1,OAS2,OAS3,OASL,PARP9,PLSCR1,RSAD2,RTP4, SERPINB9,STAT1,STAT2,TLR2,TLR3,TLR4,TLR7,TLR8,TRIM22,UBD** | **Antimicrobial response** |
| **IFNG** | **CCL2,CCL3,CCL4,CCL5,CXCL8,CXCR4,IL1B,IL6,MUC1,NFKBIA,PTGS2,TLR2** | **Cell movement of carcinoma cell lines** |
| **IFNG** | **CD40LG,CD80,CD86,CEACAM1,FCER1G,IL10,IRF4,IRF8,PDCD1LG2, PRKCQ,SERPINB9,TLR3,TLR4** | **Priming of T lymphocytes** |
| **IFNG** | **CCL2,CCL5,CD2,IL1B,IL6** | **Stimulation of tumor cell lines** |
| **IFNK** | **CD80,CD86,CXCL10,CXCL9,EIF2AK2,STAT1,STAT4,TLR3,ZBP1** | **Activation of cells** |
| **IFNL2** | **CCR5,CXCL10,IFNL1** | **Cell movement of phagocytes** |
| **IFNL2** | **CCR5,CXCL10,IFNL1** | **Migration of antigen presenting cells** |
| **IFNL2** | **CCR5,IFNL1** | **Proliferation of lymphocytes** |
| **IFNL2** | **CXCL10,IFNL1,RSAD2** | **Replication of Herpesviridae** |

| | | |
|---|---|---|
| IFNW1 | CD80,CD86,CXCL10,CXCL9,MARCHF1,STAT4,TLR3,ZBP1 | Activation of cells |
| IFNW1 | CD80,CD86,CXCL10,STAT4,TLR3 | Proliferation of blood cells |
| IFNW1 | CD80,CD86,STAT4,TLR3 | Proliferation of immune cells |
| IFNW1 | CD80,CD86,STAT4,TLR3 | Proliferation of mononuclear leukocytes |
| Ige | CCL2,CCL22,CCL4,CCL5,CCR5,CEACAM1,CXCL3,EDN1,ELMO1, FCER1G,FLT1,IL10,LIF,PDGFA,PENK,PPARG,PTPN6,SPP1,TLR2, TNFSF14,TSLP | Chemotaxis |
| IGF1R | CCL5,CDH1,IL16,PTGS2 | Migration of cells |
| IHH | CXCL10,FLT1,SPP1 | T cell development |
| IKBKB | BRCA2,CCL2,CCL4,CCR1,CDC6,CSF1R,CXCL10,CXCL12,EDN1,FASLG,GBP2,IFI16,IL10,MDM2,REL,TGFBR2 | Malignant neoplasm of retroperitoneum |
| IKZF3 | DDX58,TLR4 | Quantity of lymphoid tissue |
| IL1 | CCL2,CXCL8,IL10,IL6,MMP9,PTGS2 | Cell proliferation of tumor cell lines |
| IL13 | CCL18,CCL2,CCL22,CCL4,CCL5,CCR5,CXCL10,CXCL3,CXCR2,CYSLTR1,IL10,OPRM1,SPP1,TLR4,TSLP | Chemotaxis of mononuclear leukocytes |

| | | |
|---|---|---|
| IL13 | CCL18,CCL2,CCL22,CCL4,CCL5,CCR5,CXCL10,CXCL3,CXCR2,CYSLTR1,IL10,OPRM1,SPP1,TLR4,TSLP | Homing of mononuclear leukocytes |
| IL15 | CD40LG,CD80,CD86,CEACAM1,IL10 | Priming of T lymphocytes |
| IL15RA | CD226,EDN1,IL10,IL2RA,PTPRC | Organismal death |
| IL17A | CCL2,CCL22,CCL4,CCL5,CXCL10,CXCL12,CXCL13,IL10,LIF,PECAM1, TLR2,TLR4 | Interaction of lymphocytes |
| IL17RA | CCR1,CXCL12,CXCR2,MMP13 | Quantity of cells |
| IL18 | CCL2,CD244,FASLG,HAVCR2,ULBP2 | Cytotoxicity of cells |
| IL18 | CCL2,CCL3,CCL4,CD244,CXCL10,CXCL8,FASLG,ICAM1,IL10,IL1A,IL1B,IL6,MMP9 | Leukocyte migration |
| IL18 | CXCL8,IL10,IL1B,IL6 | Stimulation of leukocytes |
| IL1A | CCL2,CCL5,IL1B,IL6 | Stimulation of tumor cell lines |
| IL1B | CCL2,CCL22,CCL4,CD40LG,CD80,CXCL11,CXCL12,CXCL13,CXCL9,IL6R,LIF,SPP1,TLR4 | Adhesion of lymphocytes |
| IL1B | CCL2,CD40,CXCR4,IL10,IL6,MUC1,PTGS2,THBS1,VEGFA | Growth of tumor |
| IL1B | CXCL10,CXCL9,EDN1,IGF1,ITGA1,TLR2,TLR4,TSLP | Proliferation of glomerular cells |
| IL1B | CXCL10,CXCL9,EDN1,IGF1,ITGA1,TLR2,TLR4,TSLP | Proliferation of mesangial cells |
| IL1B | CD40,CXCR4,IL10,IL6,MUC1,PTGS2,THBS1,VEGFA | Proliferation of tumor cells |

| | | |
|---|---|---|
| IL1B | C3,CCL2,CCL3,CCL4,CCL5,CCL7,CXCL10,CXCL8,CXCR4,FPR2,IL6, VEGFA | Quantity of Ca2 |
| IL1B | CCL2,CCL3,CCL5,CCL7,CXCL8,CXCR4,IL6R,VEGFA | Recruitment of cells |
| IL1B | CCL2,CCL5,IL6,VEGFA | Stimulation of tumor cell lines |
| IL21 | BCL6,IL10,IL6 | Cell viability of lymphocytes |
| IL21 | BCL6,IL10,IL6 | Cell viability of mononuclear leukocytes |
| IL21 | IL10,IL6,TBX21 | Differentiatio n of naive T lymphocytes |
| IL23 | IL10,IL2RA,TLR2,TLR4 | Inflammatory Bowel Disease |
| IL3 | CD40,IL1B,IL6 | Growth of malignant tumor |
| IL3 | CD40,IL1B,IL6 | Proliferation of cancer cells |
| IL33 | CCL2,CCL4,CCL5,CCR1,CCR2,CD69,CXCL10,CXCL3,CXCR2,DOCK2, FLT1,IL10,PTPRC,REL,TLR2,TSLP | Chemotaxis of myeloid cells |
| IL33 | CCL2,CCL4,CCL5,CCR1,CCR2,CD69,CXCL10,CXCL3,CXCR2,DOCK2, FLT1,IL10,REL,TLR2,TSLP | Chemotaxis of phagocytes |
| IL4 | CCL3,CXCL8,CXCR4,FLT3LG,ICAM1,IL10,IL1B,IL6 | Cell viability of blood cells |

| | | |
|---|---|---|
| IL5 | A2M,BCL3,BIRC2,CCL2,CCND2,CCR1,CCR2,CD40LG,CD80,CEACAM1,CX3CR1,HLX,IL10,IL2RG,IL6R,LMNB1,PTPRC,SKA1,SPN,TNFRSF9 | Cell survival |
| IL5 | A2M,BCL3,BIRC2,CCL2,CCND2,CCR1,CCR2,CD40LG,CD80,CEACAM1,CX3CR1,IL10,IL2RG,IL6R,LMNB1,PTPRC,SKA1,SPN,TNFRSF9 | Cell viability |
| IL5 | BCL3,CD40LG,CD80,CX3CR1,IL10,IL2RG,PTPRC,SPN,TNFRSF9 | Cell viability of leukocytes |
| imiquimod | CD274,CD80,IL10,IL2RA,RSAD2,STAT1,TLR7 | Differentiation of T lymphocytes |
| Immunoglobulin | CCL2,CCL3,CCL4,CCL7,CCL8,CXCL10,CXCL2,IL1B,IL6,NFKBIA | Inflammatory response |
| infliximab | EDN1,GNLY,INHBA,TNFRSF9 | Activation of phagocytes |
| Interferon alpha | AIM2,APOBEC3A,APOBEC3F,BCL3,CCL22,CCL5,CD86,CXCL10, DDX58,EIF2AK2,FCGR1A,IFI16,IFI44,IFI44L,IFI6,IFIH1,IFIT1,IFIT2, IFIT3,IFITM1,IFITM2,IFITM3,IFNL1,ISG15,MX1,MX2,NT5C3A,OAS1,OAS2,OAS3,PARP9,PLSCR1,RNASEL,RSAD2,RTP4,STAT1,STAT2, TLR3,TLR7,TLR8,TRIM22,TRIM5,USP25,ZC3HAV1 | Antiviral response |
| Interferon alpha | APOBEC3F,DDX58,FCGR1A,IFIH1,IFITM2,IFITM3,IFNL1,IFNL2,OAS1,PARP12,PML,SP110,STAT2,TLR3 | Replication of Rhabdoviridae |
| ionomycin | CD40LG,CD69,CXCL10,FASLG,IL10,TLR2,TNFSF10,TNFSF14 | Stimulation of leukocytes |
| IRF5 | CCL4,CCL5,CXCL10,CXCL11,DDX58,STAT1 | Lymphocyte migration |
| isoproterenol | EDN1,FASLG,IL10,LIF | Stimulation of cells |

| | | |
|---|---|---|
| ITGB1 | CCL2,CXCL8,ITGA5,MMP9,PTGS2,VEGFA | Angiogenesis |
| ITGB1 | CXCL8,ITGA5,MMP9,PLAU,VEGFA | Cell spreading |
| ITGB1 | CCL2,CXCL8,ITGA5,MMP9,PLAU,VEGFA | Leukocyte migration |
| ITK | CCL5,CX3CR1,CYSLTR2,FCER1G,IL10,LYN | Mobilization of Ca2 |
| JAK1 | BCL3,EIF2AK2,IL10,LIF,STAT1 | Quantity of stem cells |
| JAK2 | CCND2,ICAM1,LYN,PTGS2 | Cell viability |
| JAK2 | ACE,CASP1,CCL2,CCL5,CXCL10,CYBB,LYN,PECAM1,PRKCB,STAT1 | Inflammatory response |
| JAK2 | CASP1,CCL2,CCL5,CXCL10,CYBB,LYN,PECAM1,PRKCB,SLC1A3, STAT1 | Migration of cells |
| Jnk | CCL2,CCL3,CCL5,MMP9,PTGS2,SERPINE1 | Cell movement of muscle cells |
| JQ1 | IL10,TLR2,TNFSF10 | Stimulation of leukocytes |
| JUN | ABCB1,CCL2,CXCL8,DUSP1,DUSP6,FASLG,FOSL1,IL10,IL1A,IL1B,IL6,NAMPT,NFKB2,NFKBIA,PTGS2,VEGFA | Cell viability of tumor cell lines |
| JUNB | FOSL1,IL1B,NAMPT,NFKB2,SERPINE1 | Expression of RNA |
| KLF4 | ACOD1,ADAMTS1,ADGRE1,AIF1,CCL2,CCND2,CLIC4,CSF1R,DLX3, EMX2,EOMES,FLT1,GBP4,GREB1,IFITM3,IL10,INHBA,LAMA1, MMP13,MRC1,MYH11,MYOCD,PECAM1,TF,TPK1,USP18 | Head and neck tumor |
| lactic acid | CCL2,CLEC4E,CXCL10,CXCL12,IGF1,LY96 | Inflammatory response |
| lactic acid | CCL2,CLEC4E,CXCL10,CXCL12,IGF1,PCK1 | Quantity of cells |

| | | |
|---|---|---|
| leukotriene D4 | ADAMTS1,ARID5B,CCL2,CCL4,CYSLTR1,HLX | Migration of cells |
| LGALS3 | ABCB1,CCL2,DUSP6,ICAM1,IL6,SPP1 | Cell viability |
| LGALS8 | CCL2,CCL5,CXCL3,FASLG | Adhesion of blood cells |
| LGALS8 | CCL2,CCL5,CXCL3,FASLG | Adhesion of immune cells |
| LGALS8 | CCL2,CCL5,CXCL1,CXCL8,FASLG,IL6 | Cell movement of tumor cell lines |
| LGALS8 | CCL2,CCL5,CXCL3,FASLG | Interaction of phagocytes |
| LGALS8 | CCL2,CCL5,CXCL3,FASLG | Invasive tumor |
| LGALS8 | CCL2,CCL5,CXCL3,FASLG | Leukopoiesis |
| LGALS8 | CCL2,CCL5,CXCL3,FASLG | Stimulation of leukocytes |
| lipoarabinomannan | CCL4,CCR5,STAT1,TLR4 | Activation of leukocytes |
| lipoteichoic acid | CCL2,CD80,CD86,CXCL10,CYBB,IL10,TLR2,TLR3,TLR4,TLR7 | Leukopoiesis |
| losartan potassium | CXCL10,CXCL3,CYBB,EDN1,EDNRA,SPP1,TGFBR2 | Cell movement of granulocytes |
| losartan potassium | CXCL10,CXCL3,CYBB,EDN1,EDNRA,SPP1,TGFBR2 | Cell movement of neutrophils |
| LRBA | CCL5,CXCL10,IFIT2 | Inflammation of organ |
| lysophosphatidic acid | CCL2,CCL4,CCL5,CXCL10,EDN1,MSR1 | Activation of antigen |

| | | |
|---|---|---|
| | | presenting cells |
| lysophosphatidylcholine | CCL2,CCL5,TNFRSF9 | Cell movement of dendritic cells |
| MALP-2s | CCL4,CD86,CXCL3,IL10,TLR2,TLR4 | Cellular homeostasis |
| MAPK1 | ANPEP,ATP2A1,CCDC82,CCL2,CCL4,CRYBG1,DDX58,FASLG,GBP2, IFI16,IFIT3,IFITM1,IFITM3,IL2RA,PLSCR1,ROS1,SUN2,TNFSF10, TRIM22,TRIM5 | Renal cancer |
| MAPK3 | FOXP3,FURIN,IL10,IL2RA,SIRT6,SPP1 | Organismal death |
| MAPK8 | ETS1,FOSL1,JUN,MMP9,PTGS2 | Cell movement of tumor cell lines |
| MAPK8 | ETS1,FOSL1,JUN,MMP9,PTGS2 | Cell proliferation of tumor cell lines |
| MAPK8 | ETS1,FOSL1,JUN,MMP9,PTGS2 | Invasion of tumor cell lines |
| MAPK8 | ETS1,FOSL1,JUN,MMP9,PTGS2 | Migration of tumor cell lines |
| MAPK9 | CCL2,CCL5,EDN1,FASLG,FOXP3,IFI16,LIF,PPARG,TLR3 | Activation of cells |
| MASTL | ASAP1,CYP1B1,E2F3,RAP2C,SH2B3 | Cell movement |

| | | |
|---|---|---|
| MAVS | DDX58,IFITM3,IFNL1,IFNL2,OAS1,OASL,PARP12,STAT2 | Replication of vesicular stomatitis virus |
| MED1 | ADIPOQ,BCL3,CCND2,CDK2,IGF1,IKZF1,IRF8,LIF,MDM2,PPARG | Organismal death |
| MESP1 | MYOCD,ZEB1,ZEB2 | Organismal death |
| MET | EDN1,IGF1,INHBA,SPP1 | Chemotaxis |
| MIF | CCL2,CCL4,CCR1,CCR5,CXCL3,IL10,TLR4 | Chemotaxis of mononuclear leukocytes |
| mir-1 | CCL2,CXCL1,MMP9,VEGFA | Angiogenesis |
| mir-1 | CCL2,CXCL1,MMP9,VEGFA | Cell proliferation of tumor cell lines |
| mir-135 | CASP1,CXCL12,EDN1,MMP13,SPP1 | Quantity of cells |
| miR-141-3p (and other miRNAs w/seed AACACUG), MIR17HG | AKR1C1/AKR1C2,ALPL,ATF2,BAP1,BMP4,CCM2,CDH2,CDKN1A, CDKN1B,CELSR1,CLIC1,CTBP2,CTNNB1,DLL1,DLX5,ECE1,ECSIT, EPCAM,FZD1,FZD4,GEMIN2,GJB3,GRB7,HSPG2,KCNK1,LAMA5, LRP1,LRP4,LSR,LTBP1,MSLN,NCOR2,PITX1,PITX2,PKD1,PTK7, PTPRU,RNF128,SATB1,SKI,SNAI1,SORT1,SP5,TAX1BP1,TOB2,TP73, TWSG1,ZNF703,ZNRF3 | Intestinal tumor |
| mir-146 | CCL3,CXCL10,IL1B,IL6,IRAK1 | Expression of RNA |

| | | |
|---|---|---|
| miR-146a-5p (and other miRNAs w/seed GAGAACU) | CAMP,CCL2,CD40,CXCL8,IL10,IL1R1,PRF1,S100A12 | Activation of cells |
| miR-146a-5p (and other miRNAs w/seed GAGAACU) | CCL2,CXCL3,IL10,IL1RL2,STAT1,TLR1,TLR4 | Leukopoiesis |
| mir-15 | CCL2,CCL5,IL1B,IL6 | Cell movement of carcinoma cell lines |
| mir-15 | CCL2,CCL5,IL1B,IL6 | Cellular homeostasis |
| mir-15 | CCL2,CCL5,IL1B,IL6 | Inflammatory response |
| miR-155-5p (miRNAs w/seed UAAUGCU) | CASP1,CCL4,CD69,CSF1R,MAFB,RAB27B,RCOR1,SLA | Quantity of cells |
| mir-17 | BIRC2,BIRC3,MDM2,MMP13,PURA | Quantity of blood cells |
| MIR17HG | ALPL,APCDD1,BMP4,CCM2,CDKN1A,CELSR1,CPE,DLL1,ECE1,FZD1,HSPG2,KLF2,LAMA5,LRP1,LRP4,LTBP1,NCOR2,PITX1,PITX2,PKD1,PTK7,PTPRU,RELA,SKI,SORT1,SP5,TAGLN,TAX1BP1,TWSG1,ZNF703,ZNRF3 | Frequency of tumor |

| | | |
|---|---|---|
| MIR17HG | ALPL,APCDD1,BMP4,CCM2,CDKN1A,CELSR1,CPE,DLL1,ECE1,FZD1,HSPG2,KLF2,LAMA5,LRP1,LRP4,LTBP1,NCOR2,PITX1,PITX2,PKD1,PTK7,PTPRU,RELA,SKI,SORT1,SP5,TAGLN,TAX1BP1,TWSG1,ZNF703,ZNRF3 | Incidence of tumor |
| mir-181 | CD69,IRF8,PTPN22,ZEB2 | Lymphopoiesis |
| mir-204 | CXCL1,CXCL2,CXCL8,IL1B,IL6 | Inflammatory response |
| mir-204 | CXCL1,CXCL2,CXCL8,IL1B,IL6 | Invasion of cells |
| mir-223 | CLEC4E,MMP13,MMP8,SOAT1,TLR4,TLR7 | Quantity of phagocytes |
| mir-25 | ADM,BBC3,E2F3,EDNRB,MDM2,REV3L | Organismal death |
| mir-29 | COL4A2,EOMES,IDO1,IGF1,IL2RA,ITGA11,PDGFA,PMP22 | Organismal death |
| miR-450a-5p (and other miRNAs w/seed UUUGCGA) | ITGA5,MMP9,SERPINE1,SPP1 | Cell movement of tumor cell lines |
| MMP1 | CXCL8,JUN,MMP9,PLAU,SERPINE1 | Cell movement of breast cancer cell lines |
| MMP1 | CXCL8,JUN,MMP9,PLAU,SERPINE1 | Migration of tumor cell lines |
| MRTFB | ADAMTS1,ADM,ARID5B,CDK2,DAB2,EDN1,EDNRA,FLI1,INHBA | Organismal death |
| MUC1 | CCND2,IL10,MMP13,MRC1,PDGFA,STAT1 | Organismal death |

| | | |
|---|---|---|
| MUC1 | CCND2,IFIT3,IFITM1,IL10 | Renal cancer |
| MYD88 | CCL2,CCL4,CCL5,CCRL2,CXCL10,CXCL12,CXCL13,CXCL3,EDNRB, FCER1G,IGF1,IL10,INHBA,PDE4B,SPP1,TLR2 | Chemotaxis |
| N-acetylmuramyl-L-alanyl-D-isoglutamine | CASP1,CCL2,CD80,CD86,CXCL11,CXCL3,IL10 | Stimulation of leukocytes |
| NAMPT | CXCL10,CXCL8,IL6,MMP9 | Cell movement of tumor cell lines |
| NCOA3 | CDC6,EOMES,IGF1,PPARG,SLC16A3 | Invasive tumor |
| NCSTN | CD68,CSF1R,FCER1G,GFI1B,GRN,IRF8 | Organismal death |
| NFATC1 | CCND2,CD40LG,CX3CR1,CYBB,EDN1,FASLG,FOXP3,IL2RA,ITPR2, MYH11,PPARG,SPP1,TNFSF10,UTRN | Organismal death |
| NFkB (complex) | C3,CAMP,CCL2,CCL3,CCL3L1,CCL4,CCL5,CCL8,CCR7,CD40,CLU,CXCL1,CXCL10, CXCL2,CXCL8,CXCR4,EDNRB,FASLG,ICAM1,IL10,IL1B,IL6, LTA,MMP9,NAMPT,NFKB2,NFKBIA,PLAU,PTGS2 | Cell movement |
| NFkB (complex) | C3,CAMP,CCL2,CCL3,CCL4,CCL5,CCL8,CCR7,CD40,CLU,CXCL1,CXCL10,CXCL2,CXCL8,CXCR4,EDNRB,FASLG,IL1B,IL6,MMP9,NAMPT, NFKBIA,PLAU,PTGS2 | Cell movement of tumor cell lines |
| NFkB (complex) | CCL2,CCR7,CLU,CXCL1,CXCL2,CXCL8,CXCR4,EDNRB,FASLG,IER3,IL1B,IL6,MMP9,NAMPT,NFKBIA,PLAU,PTGS2 | Invasion of cells |
| NFkB (complex) | CCL2,CCL5,CXCL10,CXCL8,CXCR4,FASLG,IL1B,IL6,MMP9,PLAU | Migration of breast cancer cell lines |

| | | |
|---|---|---|
| **NFkB (complex)** | **C3,CAMP,CCL2,CCL3,CCL3L1,CCL4,CCL5,CCL8,CCR7,CD40,CLU,CXCL1,CXCL10, CXCL2,CXCL8,CXCR4,EDNRB,FASLG,ICAM1,IL10,IL1B,IL6, LTA,MMP9,NFKB2,NFKBIA,PLAU,PTGS2** | **Migration of cells** |
| **NLRP12** | **CASP1,CXCL12,CXCL13,LY96,TLR4** | **Inflammatory response** |
| **NLRP12** | **BCL3,CASP1,CXCL12,CXCL13,TLR4** | **Quantity of mononuclear leukocytes** |
| **NOD2** | **CCL2,CCL5,CXCL10,CXCL3,IL10,PDE4B** | **Chemotaxis of phagocytes** |
| **NOD2** | **CCL2,CCL5,CXCL10,CXCL3,IL10,PDE4B** | **Homing of leukocytes** |
| **NONO** | **PDE10A,PDE1B,PDE3A,PDE3B,PDE4B,PDE4D** | **Development of genital tumor** |
| **NONO** | **PDE3A,PDE3B,PDE4B,PDE4D** | **Lymphoid cancer** |
| **NONO** | **PDE3A,PDE3B,PDE4B,PDE4D** | **Lymphoprolif erative disorder** |
| **NONO** | **PDE10A,PDE1B,PDE3A,PDE3B,PDE4B,PDE4D** | **Neoplasia of cells** |
| **NONO** | **PDE3A,PDE3B,PDE4B,PDE4D** | **Urinary tract cancer** |
| **NOX4** | **CCL2,CDH1,CXCL8,ICAM1,VEGFA** | **Adhesion of tumor cell lines** |
| **NOX4** | **CCL2,CDH1,CXCL8,ICAM1,VEGFA** | **Vasculogenesi s** |
| **NPC1** | **CCR5,MMP8,STAT1,STAT4,TLR4** | **Cell death of immune cells** |
| **NPC2** | **PPARG,PTGS1,STAT1,STAT2,STAT4,TCF21,TLR4** | **Morbidity or mortality** |

| | | |
|---|---|---|
| ORMDL3 | CXCL10,RNASEL,TNFSF13B | Cell death of immune cells |
| OSM | CASP4,CCL2,CCL5,CCND2,CXCL10,CXCL12,CXCL13,CXCL3,IL10,LIF, SPP1,TLR2,TLR3 | Stimulation of cells |
| P38 MAPK | CCL2,CXCL2,ICAM1,IL1B,IL6,JUN,VEGFA | Binding of lymphoma cell lines |
| P38 MAPK | CCL2,CCL5,CXCL10,CXCL8,CXCR4,ETS1,IL1B,IL32,IL6,JUN,MMP9, SERPINE1,VEGFA | Cell movement of breast cancer cell lines |
| P38 MAPK | CCL2,CCR7,CXCL8,CXCR4,ETS1,IL1B,IL32,IL6,JUN,MMP9,PTGS2, VEGFA | Invasion of tumor cell lines |
| P38 MAPK | CCL2,CCL5,CCR7,CXCL10,CXCL8,CXCR4,ICAM1,IL10 | Lymphocyte migration |
| P38 MAPK | CCL2,CCL5,CCL8,CCR7,CD36,CD40,CXCL1,CXCL10,CXCL2,CXCL8,CXCR4,ETS1,ICAM1,IL10,IL1B,IL32,IL6,JUN,MMP9,PTGS2,S100A12, SERPINE1,TLR2,VEGFA | Migration of cells |
| P38 MAPK | CCL2,CD40,CXCL1,CXCL8,ETS1,ICAM1,PTGS2,SERPINE1,VEGFA | Migration of endothelial cells |
| PDCD1 | CD3D,CD80,CD86,GNLY,HLA-DMA,IL10,IL2RA,STAT1 | Quantity of T lymphocytes |
| PDGF BB | CCL2,CXCL8,DUSP1,DUSP6,IER3,IL1B,IL6,JUN,NAMPT,NR4A1,PLAU | Invasion of cells |
| pentoxifylline | ACE,CCL5,CCR5,CXCL3,CYBB,IL10,REL | Neoplasia of cells |
| PF4 | CCL2,CCL3,CCL4,CXCL8,IL1A,IL1B,IL6 | Cell movement of blood cells |

| | | |
|---|---|---|
| PF4 | CCL4,CXCL3,FLI1,INHBA,REL,STAT1 | Development of hematopoietic cells |
| PF4 | CCL4,CXCL3,FLI1,INHBA,REL,STAT1 | Development of hematopoietic system |
| PF4 | CCL2,CCL3,CCL4,CXCL8,IL1A,IL1B,IL6 | Inflammatory response |
| PF4 | CCL2,CCL3,CCL4,CXCL8,IL1A,IL1B,IL6 | Leukocyte migration |
| phospholipid | CCL4,CCL5,CXCL10,CXCL3 | Activation of myeloid cells |
| phospholipid | CCL4,CCL5,CXCL10,CXCL3 | Activation of phagocytes |
| phospholipid | CCL4,CCL5,CXCL10,CXCL3 | Adhesion of blood cells |
| phospholipid | CCL4,CCL5,CXCL10,CXCL3 | Binding of leukocytes |
| phospholipid | CCL4,CCL5,CXCL10,CXCL3 | Recruitment of myeloid cells |
| phospholipid | CCL4,CCL5,CXCL10,CXCL3 | Recruitment of phagocytes |
| phytohemagglutinin | ADORA3,CCL2,CCL5,CD69,FASLG,GZMB,IL10,PTPRC,STAT1,STAT4, TNFSF10 | Cytotoxicity |
| PI3K (complex) | CCL3,CCL4,CXCL8,FOSL1,IL1B,IL6,PTGS2,VEGFA | Chemotaxis |
| PLAG1 | AKAP8L,CDKN1C,CNKSR1,MAPK11,PIR,PTP4A3,ROM1,SMARCD3 | Expression of RNA |
| PLK4 | CXCL10,EDN1,MMP13,RGS1 | Quantity of cells |

| | | |
|---|---|---|
| PML | IFIH1,OAS1 | Replication of Rhabdoviridae |
| poly rI:rC-RNA | DDX58,FCGR1A,IFIH1,IFITM3,IFNL1,IFNL2,OAS1,OASL,PARP11,SP110,STAT2,TLR3 | Replication of Rhabdoviridae |
| PPP2R5C | BCL2A1,CXCL8,FASLG,TNFAIP3 | Cell viability |
| PRKCA | CCL2,ETS1,JUN,MMP9,PTGS2,VEGFA | Invasion of tumor cell lines |
| PRKCD | CXCL8,DUSP6,FOSL1,IL1B,IL32,IL6,MMP9,NFKBIA | Invasion of tumor cell lines |
| PRKD1 | CCL4,CCL5,CXCL10,CXCL3,IL10 | Mobilization of Ca2 |
| PRL | CDH1,CXCL10,JUN | Adhesion of tumor cell lines |
| PROCR | CCL22,IRF8,LIF | Binding of blood cells |
| PTEN | CCL5,CXCL8,IL6,PTGS2 | Cell movement of carcinoma cell lines |
| PTEN | CD274,CXCL8,IL6,PTGS2 | Cell proliferation of tumor cell lines |
| PTEN | CCL5,CXCL8,IL6,PTGS2 | Cell viability |
| PTEN | CCL5,CXCL8,IL6,PTGS2 | Chemotaxis |
| PTEN | CD274,CXCL8,IL6,PTGS2 | Invasion of tumor cell lines |

| | | |
|---|---|---|
| PTEN | CCL5,CXCL8,IL6,PTGS2 | Migration of tumor cell lines |
| PTPRE | CXCL3,INHBA | Activation of myeloid cells |
| PTPRE | CXCL3,INHBA | Activation of phagocytes |
| PTPRE | CXCL3,INHBA | Invasive tumor |
| PTX3 | CCL2,CCL4,CCL5,CCR2,CCR5,CX3CR1,TLR2,TLR4 | Cell movement of myeloid cells |
| PTX3 | CCL2,CCL4,CCL5,CCR2,CCR5,CX3CR1,TLR2,TLR4 | Cell movement of phagocytes |
| PTX3 | CCL2,CCL4,CCL5,CCR2,CCR5,CX3CR1,TLR2,TLR4 | Cellular infiltration |
| PTX3 | CCL2,CCL4,CCL5,CCR2,CCR5,CX3CR1,TLR2,TLR4 | Cellular infiltration by blood cells |
| PTX3 | CCL2,CCL4,CCL5,CCR2,CCR5,CX3CR1,TLR2,TLR4 | Cellular infiltration by leukocytes |
| PTX3 | CCL2,CCL4,CCL5,CCR2,CCR5,CX3CR1,IGF1,TLR2,TLR4 | Migration of cells |
| reactive oxygen species | CASP1,FASLG,GZMB | Cytolysis |
| REL | CCL2,CCL4,CD40LG,CD80,CD86,CXCL10,IRF8 | Binding of blood cells |
| RELA | CCL2,CCL3,CCL5,CXCL10,CXCL2,CXCL8,CXCR4,ICAM1,IL10,IL1A,IL1B,IL6,PLAU, TLR2,VEGFA | Binding of blood cells |

| | | |
|---|---|---|
| **REST** | **CXCL12,EOMES,GRIN2B,KCNJ6,NTRK3,OPRM1,PENK,PTPRN, SLC12A5** | **Organismal death** |
| **RETNLB** | **CCL2,CCL5,CCR1,CCR2,CCR5,CD86,IGF1,LMNB1,MMP12,TLR2** | **Quantity of cells** |
| **RIPK1** | **CCL2,CXCL1,CXCL8,IL6** | **Cell proliferation of tumor cell lines** |
| **RIPK1** | **CCL2,CXCL1,CXCL8,IL6** | **Inflammatory response** |
| **RIPK2** | **CCL5,CD69,CD86,CLEC4E,CXCL10,PDE4B,RASGRP3,TSLP** | **Quantity of leukocytes** |
| **RIPK2** | **CCL5,CD69,CLEC4E,CLEC5A,CXCL10,CXCL3,MCOLN2,PDE4B** | **Recruitment of myeloid cells** |
| **RIPK2** | **CCL5,CD69,CLEC4E,CLEC5A,CXCL10,CXCL3,PDE4B** | **Recruitment of neutrophils** |
| **RIPK2** | **CCL5,CD69,CLEC4E,CLEC5A,CXCL10,CXCL3,MCOLN2,PDE4B** | **Recruitment of phagocytes** |
| **RNASE2** | **CCL2,CCL22,CCL4,CCL5,CCL8,CXCL10,CXCL9,IL10** | **Chemotaxis of blood cells** |
| **RNASE2** | **CCL2,CCL22,CCL4,CCL5,CCL8,CXCL10,CXCL9,IL10** | **Chemotaxis of leukocytes** |
| **RNASE2** | **CCL2,CCL22,CCL4,CCL5,CCL8,CXCL10,CXCL9,IL10** | **Chemotaxis of phagocytes** |
| **RNASE2** | **CCL2,CCL22,CCL4,CCL5,CCL8,CXCL10,CXCL9,IL10** | **Homing of blood cells** |
| **RNASE2** | **CCL2,CCL22,CCL4,CCL5,CCL8,CXCL10,CXCL9,IL10** | **Homing of leukocytes** |
| **SELP** | **CCL2,CCL4,IGF1,SERPINB9** | **Activation of mononuclear leukocytes** |

| | | |
|---|---|---|
| SELPLG | CCL2,CCL3,CCL4,CXCL2,CXCL8,CXCR4,IL1B,IL1R2 | Binding of tumor cell lines |
| SELPLG | CCL2,CCL3,CCL4,CXCL2,CXCL8,CXCR4,IL10,IL1B | Cell movement of leukocytes |
| SELPLG | CCL2,CCL3,CCL4,CXCL2,CXCL8,CXCR4,IL10,IL1B | Cell movement of myeloid cells |
| SELPLG | CCL2,CCL4,IGF1,IL10 | Chemotaxis |
| SMARCA4 | ACE,ADIPOQ,CASP1,CCL2,CCR2,CEACAM1,IFI16,IGF1,IL20RB,INHBA,MARCHF1, MICB,PPARG,SPP1,TLR2,TNFRSF9,TNFSF10 | Activation of cells |
| SMPD1 | CCL2,CCL4,CCL5,CXCL9,JAK2,TLR2,TNFSF10 | Activation of cells |
| SMPD1 | CCL2,CCL4,CCL5,CXCL11,CXCL9,JAK2,TLR2 | Cell movement of T lymphocytes |
| SOCS1 | CCL2,CD69,CD86,CXCL10,DDX58,FCER1G,STAT1,TLR3 | Activation of mononuclear leukocytes |
| SOD3 | EDN1,EDNRA,EDNRB,MC1R,WNT7A | Migration of cells |
| ST8SIA1 | C1QA,C3AR1,CDK2 | Quantity of blood cells |
| STAR | CCL5,CD86,CLEC7A,SPP1 | Cellular homeostasis |
| STAT3 | CCL2,CD40,CEACAM1,CXCL8,EGR3,MMP9,NFATC2,PTGS2, SERPINE1,VEGFA | Cell movement of endothelial cells |

| | | |
|---|---|---|
| STAT4 | CCL2,CD40LG,IL10,STAT1 | Cytotoxicity of lymphocytes |
| STAT5a/b | CD69,FOXP3,IGF1,IL2RA,LCP2,TSLP | Differentiation of T lymphocytes |
| SYVN1 | BCAT1,CYP1B1,DAB2,DPYD,GPRC5A,HERC5,IFI44,IFITM2,IL7R,OLR1,SETX,SLC2A3,ZC3HAV1 | Neoplasia of cells |
| TCF | ALDH1A1,BCL2L1,BMP4,EPCAM,FGF18,FOS,JUNB,MYC,SFN | Colony formation of cells |
| TCF3 | ADIPOQ,CD3D,DCK,GFI1B,IL2RA,IL7R,LMO2 | Development of hematopoietic cells |
| TCR | CCL22,CCL5,CD40LG,CD69,CXCL10,CXCL11,CXCL13,EIF4E,FASLG,IL10,P2RX7,PTPRC,SLC2A3,SPP1,TLR3,TLR4,TNFSF13B | Stimulation of cells |
| TEAD2 | ADM,ARID5B,EDN1,EDNRA,TNNT2 | Organismal death |
| telmisartan | ADIPOQ,MAS1,PPARG | Chronic renal failure |
| telmisartan | ADIPOQ,MAS1,PPARG | End stage renal disease |
| TFEB | ARSB,ATP6V1B2,CCL5,MCOLN1,PPARG | Cellular homeostasis |
| TGM2 | C3,CCL2,CCL3,CXCL10,CXCL8,IL1B,JAML,MMP9,S100A8,SPP1 | Cell movement of myeloid cells |
| TGM2 | CCL2,CCL3,CXCL8,IL1B,MMP9 | Migration of myeloid cells |
| THRB | CCR1,CCR6,CXCR4,LEF1,MMP9,PTGS2,SPP1 | Migration of cells |
| Tlr | CCL2,CXCL8,IL10,IL6 | Cell viability |

| Tlr | CASP4,CCL2,CCL5,CD80,CD86,CXCL10,CXCL3,IL10,TNFSF13B | Stimulation of leukocytes |
|---|---|---|
| TLR2 | CCL2,CCL5,IL10,IL1B,IL6 | Activation of mononuclear leukocytes |
| TLR2 | CCL2,CCL5,CXCL8,GATA3,IL1B,IL6,PTGS2 | Cell movement of carcinoma cell lines |
| TLR2 | CCL2,CXCL8,GATA3,IL10,IL1B,IL27,IL6 | Differentiatio n of mononuclear leukocytes |
| TLR2 | CCL2,GATA3,IL10,IL1B,IL27,IL6 | Hematopoiesi s of mononuclear leukocytes |
| TLR3 | C3,CXCL10,CXCL8,IL10,IL1B,IL6 | Immune response of cells |
| TLR4 | CCL2,CCL4,CCL5,CCR5,CD80,CD86,CXCL10,FASLG,IL10,RGS1,SPP1, TLR2 | Interaction of lymphocytes |
| TLR4 | C3,CCL2,CCL5,CCL8,CCR7,CXCL10,CXCL8,ICAM1,IL10,IL1B,IL6,MMP9 | Leukocyte migration |
| TLR7 | CCL2,CD40,CXCL8,IL1B,IL21R,IL6,ZAP70 | Differentiatio n of mononuclear leukocytes |
| TLR9 | CCL4,CCL5,CCR7,CD40,CD5,CXCL10,CXCL8,IL10,IL21R,IL6,LTA, NAMPT,ZAP70 | Cell movement |
| TLR9 | CD40,CD5,CXCL8,IL10,IL21R,IL6,ZAP70 | Differentiatio n of |

| | | |
|---|---|---|
| | | mononuclear leukocytes |
| TLR9 | CD40,CD5,IL10,IL21R,IL6,ZAP70 | Hematopoiesis of mononuclear leukocytes |
| TLR9 | CD40,CD5,IL10,IL21R,IL6,ZAP70 | Lymphopoiesis |
| TNF | C3,CCL2,CCL3,CCL5,CXCL8,IL1A,IL1B,TLR2 | Activation of myeloid cells |
| TNF | C3,CCL2,CCL3,CCL5,CXCL8,IL10,IL1A,IL1B,IL6,LTB,RNASE2,TLR2 | Activation of phagocytes |
| TNF | CXCL2,CXCL8,ICAM1,IL1B,ITGA4,ITGA5,PLAU,S100A8,TLR2,VEGFA | Adhesion of granulocytes |
| TNF | CCL2,CD40,CXCL1,CXCL8,CXCR4,ETS1,ICAM1,ITGA4,MMP9,PTGS2, SERPINE1,VEGFA | Cell movement of endothelial cells |
| TNF | CCL2,CCL3,CCL4,CCL5,CXCL10,CXCL8,CXCR3,CXCR4,ICAM1,ITGA4, LYN,MMP9 | Cell movement of hematopoietic progenitor cells |
| TNF | C3,CCL2,CCL4,CCL5,CXCL1,CXCL8,CXCR4,ITGA4,LYN,MMP9,PLAU, VEGFA | Cell movement of leukemia cell lines |
| TNF | C3,CCL2,CCL3,CCL4,CCL5,CXCL10,CXCL8,CXCR3,CXCR4,ICAM1, ITGA4,LYN,MMP9 | Cell movement of stem cells |
| TNF | C3,CD40,CXCL8,IL10,IL6,TNFSF13B | Cell viability of lymphatic system cells |

| | | |
|---|---|---|
| TNF | CCL2,CCL3,CCL4,CCL5,CXCL1,CXCL8,CXCR4,JUN,PIK3CD,PLAU | Chemotaxis of tumor cell lines |
| TNF | CCL2,CD274,CD40,CXCR4,IER3,IL10,IL1B,IL1R1,IL6,MUC1,PIK3CD, VEGFA | Growth of malignant tumor |
| TNF | CCL2,CCL3,CCL4,CCL5,CXCL1,CXCL8,CXCR4,JUN,PIK3CD,PLAU, VEGFA | Homing of tumor cell lines |
| TNF | CD274,CD40,CXCR4,IER3,IL10,IL1B,IL1R1,IL6,MUC1,PIK3CD,VEGFA | Proliferation of cancer cells |
| TNFRSF18 | CCL2,CXCL8,ICAM1,IL6,MMP9 | Cell movement of blood cells |
| TNFRSF18 | CCL2,CXCL8,ICAM1,IL6,MMP9 | Leukocyte migration |
| TNFRSF1A | CCL2,CXCL1,CXCL8,IL1B | Cell movement of myeloid cells |
| TNFRSF1A | CCL2,CXCL1,CXCL8,IL1B | Cell movement of neutrophils |
| TNFRSF1A | CCL2,CXCL1,CXCL8,IL1B | Cell proliferation of tumor cell lines |
| TNFRSF1A | CCL2,CXCL1,CXCL8,IL1B | Chemotaxis |
| TNFRSF1A | CCL2,CXCL1,CXCL8,IL1B | Inflammatory response |
| TNFRSF1A | CCL2,CD83,CXCL8,IL1B | Quantity of cells |

| | | |
|---|---|---|
| TNFSF11 | AIF1,CCL2,CCL4,CCL5,CCR1,CXCL3,IL10,SPP1,STAT1 | Cell movement of monocytes |
| TNFSF12 | CCL2,CCL5,CCR5,CLEC4D,CLEC4E,CXCL10,CXCL3,MMP13,TSLP | Activation of cells |
| TNFSF12 | CCL2,CCL5,CXCL8,ICAM1,IL6 | Adhesion of immune cells |
| TNFSF12 | CCL2,CXCL8,ICAM1,IL6 | Angiogenesis |
| TNFSF12 | CCL5,CXCL8,ICAM1 | Binding of granulocytes |
| TNFSF12 | CCL2,CCL5,CCR1,CCR5,CXCL10,CXCL3,MMP12,TSLP | Cell movement of mononuclear leukocytes |
| TNFSF12 | CCL2,CCL5 | Cytotoxicity |
| TNFSF12 | CCL2,CXCL8,ICAM1,IL6 | Vasculogenesi s |
| TNFSF13B | ADIPOQ,CCL2,CCL4,CD40LG,CD80,CD86,GPR183,IL10,IL2RG,SPP1 | Cellular homeostasis |
| TNFSF14 | CCL5,CXCL10,CXCL11,CXCL12,CXCL3,FLT1,INHBA | Chemotaxis of mononuclear leukocytes |
| TNFSF14 | CCL5,CXCL10,CXCL11,CXCL12,CXCL3,FLT1,INHBA | Homing of mononuclear leukocytes |
| TRAF6 | CCL2,CD274,CXCL8,IL10,IL1B,IL6 | Cell proliferation of tumor cell lines |
| TRAF6 | BIRC3,CCL2,CD274,CYBB,IL10,MMP13,UBD | Organismal death |
| tributyrin | ADM,BIRC3,CCL2,CD274,CSGALNACT2,CXCL3,EDN1,INHBA,MMP13 | Organismal death |

| | | |
|---|---|---|
| TSC2 | CDKN1A,CDKN1B,CTNNB1,FOS,RELA,S100A6 | Proliferation of fibroblast cell lines |
| U0126 | CCL2,CCL4,CCL5,CCR2,CD40LG,CXCL10,CXCL12,CXCL3,CXCL9,IL10, INHBA,MMP12,PECAM1,PPARG,PRKCB,SPP1,STAT1 | Cell movement of monocytes |
| uric acid | CASP1,CCL2,CCL5,CD80,CD86,CXCL3,IL10 | Stimulation of leukocytes |
| USP18 | CCL5,CXCL10,TNFSF10 | Activation of antigen presenting cells |
| USP18 | CCL5,CXCL10,TNFSF10 | Activation of myeloid cells |
| USP18 | CCL5,CXCL10,TNFSF10 | Activation of phagocytes |
| USP22 | CCND2,CYBB,FES,MRC1,MSR1,PTPN6,PTPRC | Organismal death |
| USP7 | FOXP3,MDM2,PLIN2,PPARG,TNNT2 | Organismal death |
| VCP | CCL2,CCL5,CXCL8,NFKB2,TNFSF13B | Cell viability |
| Vegf | CXCL12,EDN1,MMP13 | Activation of hepatic stellate cells |
| VEGFA | CCL2,CXCL10,CXCL12,FLT1,IL10 | Chemotaxis of monocytes |
| VIPR1 | CCL4,CCL5,CXCL10,CXCL3,FASLG,IL10 | Chemotaxis of phagocytes |
| VIPR1 | CCL4,CCL5,CXCL10,CXCL3,FASLG,IL10 | Homing of leukocytes |
| W7 | CCL2,CD80,CD86,IL10 | Cellular homeostasis |

| | | |
|---|---|---|
| WNT5A | CCL2,CXCL1,CXCL8,IL10,IL6,PTGS2 | Cell proliferation of tumor cell lines |
| WNT5A | CCL2,CXCL1,CXCL8,IL10,IL6,PTGS2 | Chemotaxis |
| WNT5A | CCL2,CXCL1,CXCL8,IL10,IL6,PTGS2 | Inflammatory response |
| WNT5A | CCL2,CXCL1,CXCL8,IL10,IL6,PTGS2 | Migration of cells |
